# Long non-coding RNA Neat1 and paraspeckle components are translational regulators in hypoxia

**DOI:** 10.1101/2021.02.10.430272

**Authors:** Anne-Claire Godet, Emilie Roussel, Florian David, Fransky Hantelys, Florent Morfoisse, Joffrey Alves, Françoise Pujol, Isabelle Ader, Edouard Bertrand, Odile Burlet-Schiltz, Carine Froment, Anthony K. Henras, Patrice Vitali, Eric Lacazette, Florence Tatin, Barbara Garmy-Susini, Anne-Catherine Prats

## Abstract

Internal ribosome entry sites (IRESs) drive translation initiation during stress. In response to hypoxia, (lymph)angiogenic factors responsible for tissue revascularization in ischemic diseases are induced by the IRES-dependent mechanism. Here we searched for IRES *trans*-acting factors (ITAFs) active in early hypoxia in mouse cardiomyocytes. Using knock-down and proteomics approaches, we show a link between a stressed-induced nuclear body, the paraspeckle, and IRES-dependent translation. Furthermore, smiFISH experiments demonstrate the recruitment of IRES-containing mRNA into paraspeckle during hypoxia. Our data reveal that the long non-coding RNA Neat1, an essential paraspeckle component, is a key translational regulator, active on IRESs of (lymph)angiogenic and cardioprotective factor mRNAs. In addition, paraspeckle proteins p54^nrb^ and PSPC1 as well as nucleolin and Rps2, two p54^nrb^-interacting proteins identified by mass spectrometry, are ITAFs for IRES subgroups. Paraspeckle thus appears as a platform to recruit IRES-containing mRNAs and possibly host IRESome assembly. Polysome PCR array shows that Neat1 isoforms regulate IRES-dependent translation and, more widely, translation of mRNAs involved in stress response.

**Highlights:**

- Paraspeckle formation correlates with activation of translation via internal ribosome entry sites (IRES) in mouse hypoxic cardiomyocytes as well as in tumoral cells.
- The long non-coding RNA Neat1, an essential paraspeckle component, is a key translational regulator of (lymph)angiogenic and cardioprotective factor expression in this process.
- IRES-containing mRNA is recruited into paraspeckles during hypoxia.
- Paraspeckle proteins p54^nrb^ and PSPC1 as well as two p54^nrb^-interacting proteins, nucleolin and RPS2, contribute to this process.
- Paraspeckle appears as a platform for IRESome formation in the nucleus.
- The Neat1 isoforms widely regulate the translation of mRNAs containing IRESs and of genes involved in the stress response.

**GRAPHICAL ABSTRACT:** 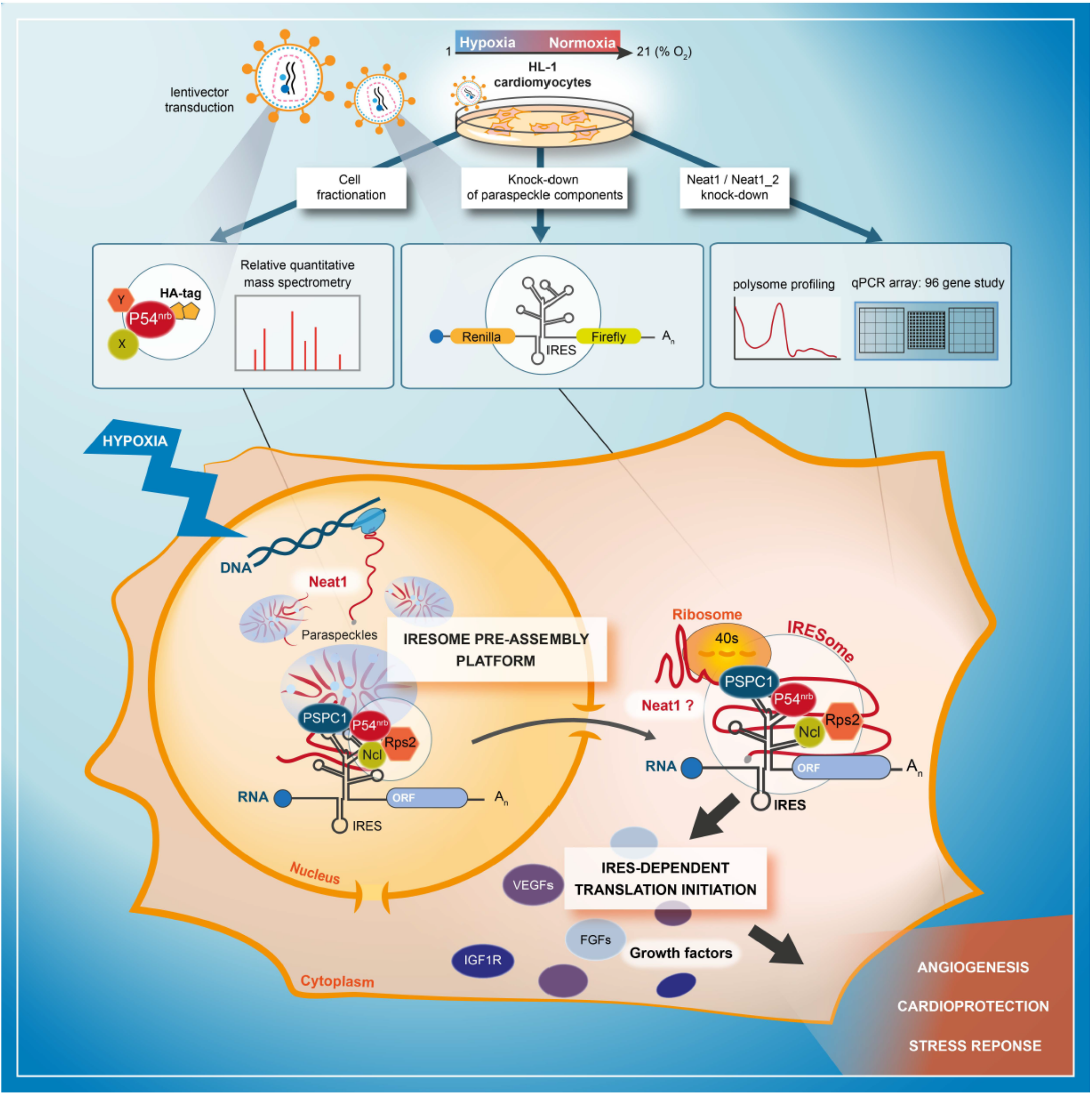

## INTRODUCTION

Cell stress triggers major changes in the control of gene expression at the transcriptional and post-transcriptional levels. One of the main responses to stress is the blockade of global translation allowing cells to save energy. This process results from inactivating the canonical cap-dependent mechanism of translation initiation (Holcik and Sonenberg, 2005). However, translation of specific mRNAs is maintained or even increased during stress via alternative mechanisms of translation initiation. One of these mechanisms involves internal ribosome entry sites (IRES), structural elements mostly present in the 5’ untranslated regions of specific mRNAs, which drive the internal recruitment of ribosomes onto mRNA and promote cap-independent translation initiation (Godet et al., 2019).

Hypoxia, or the lack of oxygen, is a major stress in pathologies such as cancer and cardiovascular diseases (Pouysségur et al., 2006). In particular, in ischemic heart failure disease, coronary artery branch occlusion exposes cardiac cells to hypoxic conditions. The cell response to hypoxia induces angiogenesis and lymphangiogenesis to reperfuse the stressed tissue with new vessels and allow cell survival (Morfoisse et al., 2014; Pouysségur et al., 2006; Tatin et al., 2017). The well-known response to hypoxia is the transcriptional induction of specific genes under the control of the hypoxia-induced factors 1 and 2 (HIF1, HIF2) (Hu et al., 2003; Koh et al., 2011). However, we have recently reported that most mRNAs coding (lymph)angiogenic growth factors are induced at the translatome level in hypoxic cardiomyocytes (Hantelys et al., 2019). Expression of these factors allows the recovery of functional blood and lymphatic vasculature in ischemic diseases, including myocardial infarction (Tatin et al., 2017; Ylä-Herttuala & Baker, 2017). The mRNAs of the major (lymph)angiogenic growth factors belonging to the fibroblast growth factor (FGF) and vascular endothelial growth factor (VEGF) families all contain IRESs that are activated in early hypoxia (Morfoisse et al., 2014; Hantelys et al., 2019).

IRES-dependent translation is regulated by IRES trans-acting factors (ITAFs) that are in most cases RNA-binding proteins acting as positive or negative regulators. A given ITAF can regulate several IRESs, while a given IRES is often regulated by several ITAFs (Godet et al., 2019), depending on the cell type or physiology. This has led to the concept of IRESome, a multi-partner ribonucleic complex allowing ribosome recruitment onto the mRNA via the IRES. ITAFs often exhibit several functions in addition to their ability to control translation. Many of them play a role in alternative splicing, transcription, ribosome biogenesis or RNA stability (Godet et al., 2019). Clearly, a large part of ITAFs are nuclear proteins able to shuttle between nucleus and cytoplasm. Previous data have also shown that a nuclear event is important for cellular IRES activity, leading to the hypothesis of IRESome formation in the nucleus (Ainaoui et al., 2015; Semler & Waterman, 2008; Stoneley et al., 2000).

Interestingly, several ITAFs are components of a nuclear body, the paraspeckle, formed in response to stress (Choudhry et al., 2015; Fox et al., 2002). These ITAFs include several hnRNPs, as well as major paraspeckle proteins such as P54^nrb^ nuclear RNA binding (P54^nrb^/NONO) and splicing factor proline and glutamine-rich (SFPQ/PSF). P54^nrb^ and SFPQ belong to the family of *drosophila melanogaster* behavior and human splicing (DBHS) proteins whose third member is the paraspeckle protein C1 (PSPC1). P54^nrb^ and SFPQ are essential for paraspeckle formation while PSPC1 is not. These three DBHS proteins are known to interact with each other and function in heteroduplexes (Fox et al., 2005; Lee et al., 2015; Passon et al., 2012). In addition, P54^nrb^ and SFPQ interact with the long non-coding RNA (lncRNA) Neat1 (nuclear enriched abundant transcript 1), that constitutes the skeleton of the paraspeckle (Clemson et al., 2009; Sunwoo et al., 2009). This lncRNA, a paraspeckle essential component, is present as two isoforms Neat1_1 and Neat1_2 whose sizes in mouse are 3.2 and 20.8 kilobases, respectively (Sunwoo et al., 2009). Its transcription is induced during hypoxia by HIF2 and promotes paraspeckle formation (Choudhry et al., 2015). Neat1 is overexpressed in many cancers (Yang et al., 2017). Recently, its induction by hypoxia has been shown in cardiomyocytes where it plays a role in cell survival (Kenneweg et al., 2019).

According to previous reports, paraspeckle is able to control gene expression via the retention of edited mRNAs and transcription factors (Hirose et al., 2014; Imamura et al., 2014; Prasanth et al., 2005). In 2017, Shen et al. have also shown that the paraspeckle might inhibit translation by sequestering p54^nrb^ and SFPQ which are ITAFs of the c-myc IRES (Shen et al., 2017).

In this study we were interested in finding new ITAFs responsible for activating (lymph)angiogenic factor mRNA IRESs in HL-1 cardiomyocytes, during early hypoxia. We have previously shown that the two paraspeckle proteins p54^nrb^ and hnRNPM are ITAFs, activators of the FGF1 IRES during myoblast differentiation (Ainaoui et al., 2015). This incited us to investigate the potential role of the paraspeckle and of Neat1 in the control of IRES-dependent translation in hypoxic cardiomyocytes. We show here that Neat1 expression and paraspeckle formation correlate with the activation of the FGF1 IRES during hypoxia, in cardiomyocytes and breast cancer cells. The knock-down of p54^nrb^, PSPC1 or Neat1 generates a decrease in FGF1 IRES activity and in endogenous FGF1 expression. Furthermore, our data revealed that IRES-containing mRNA is colocalized with Neat1 in paraspeckle during hypoxia. By quantitative mass spectrometry analysis of the p54^nrb^ interactome, we identified two additional ITAFs able to control the FGF1 IRES activity: nucleolin and ribosomal protein Rps2. Analysis of IRESs in the knock-down experiments showed that p54^nrb^ and PSPC1 are activators of several but not all IRESs of (lymph)angiogenic and cardioprotective factor mRNAs whereas Neat1 appears as a strong activator of all the cellular IRESs tested. These data suggest that the paraspeckle, via Neat1 and several protein components would be the site of IRESome assembly in the nucleus. In addition, a polysome PCR array reveals that Neat1 affects the translation of most IRES-containing mRNAs and of several mRNA families involved in hypoxic response, angiogenesis and cardioprotection.

## RESULTS

### FGF1 IRES activation during hypoxia correlates with paraspeckle formation and with Neat1 induction in different cell types

In order to analyze the regulation of IRES activity during hypoxia, HL-1 cardiomyocytes were transduced with the “Lucky Luke” bicistronic lentivector validated in our previous reports, containing the *renilla* luciferase (LucR) and firefly luciferase (LucF) genes separated by the FGF1 IRES **) (**Fig. 1A). In this construct, the first cistron LucR is expressed in a cap-dependent manner and the second cistron LucF is under the control of the IRES. The ratio LucF/LucR reflects the IRES activity.

**Figure 1.**
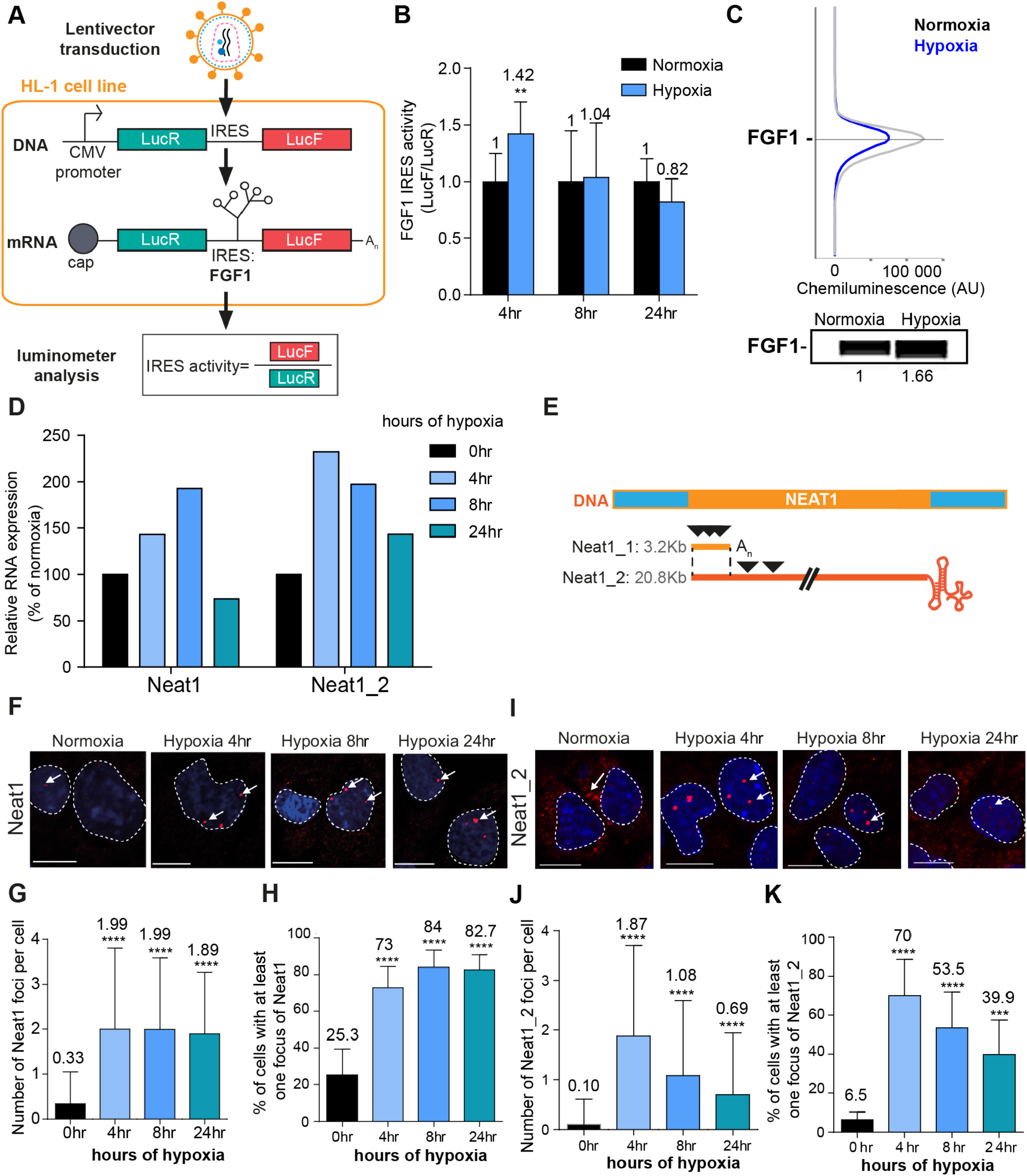
FGF1 IRES activation during hypoxia correlates with Neat1 induction and paraspeckle formation. (A) Schema depicting the Lucky Luke bicistronic construct and HL-1 cells transduced by a lentivector carrying the transgene. The LucF/LucR ratio indicates the IRES activity. (B) Activity of the human FGF1 IRES in HL-1 cardiomyocytes at 4 hr, 8 hr or 24 hr of hypoxia normalized to normoxia. The corresponding luciferase values are presented in Fig. 1-figure supplement 1, Suppl. file 1. (C) Detection of endogenous mouse FGF1 by capillary Simple Western in normoxic and hypoxic (2 hr) cardiomyocytes. The curve corresponds to the chemiluminescence signal detected with FGF1 antibody. A numerical blot is represented. Below the blot is shown the quantification of FGF1 normalized to total proteins and to control gapmer. Total proteins are detected by a dedicated channel in capillary Simple Western. (D) HL-1 cells were subjected to normoxia (0 hr) or to hypoxia during 4 hr, 8 hr and 24 hr. Neat1 and Neat1_2 expression was analyzed by droplet digital PCR (Primer sequences in Supplementary file 2). RNA expression is normalized to the normoxia time point. (E) Schema depicting the Neat1 mouse gene and the Neat1_1 and Neat1_2 RNA isoform carrying a poly(A) tail or a triple helix, respectively. Black arrowheads represent FISH probes against Neat1 and Neat1_2 (sequences in Suppl. file 2). (F-K) Neat1 (F) or Neat1_2 (I) FISH labeling in HL-1 cardiomyocytes in normoxia or at 4 hr, 8 hr and 24 hr of 1% O_2_. DAPI staining is represented in blue and Neat1 or Neat1_2 cy3 labeling in red. Nuclei are delimited by dotted lines. Scale bar=10μm. (G and J) Quantification of Neat1 (G) or Neat1_2 (J) foci per cell by automated counting (ImageJ). (H and K) Percentage of cell harboring at least one focus of Neat1 (H) or Neat1_2 (K); Histograms correspond to means + standard deviation, with Mann-Whitney (n=12) (B) or one way ANOVA (G-H, n=269-453) and (J-K, n=342-499); **p<0.01, ***<0.001, ****p<0.0001. **Figure supplement 1**. LucR and LucF expression from bicistronic vectors in hypoxic HL-1 cells. **Figure supplement 2**. Detection of Neat1 and Neat1_2 in hypoxic HL-1 by FISH. **Figure supplement 3**. FGF1 IRES activity is activated by hypoxia and inactivated after Neat1 knock-down in 67NR cells.

LucR and LucF activities were measured in HL-1 cells subjected to hypoxia for 4 hr, 8 hr or 24 hr (Fig. 1-figure supplement 1, Suppl. file 1). We previously showed that eIF2α is phosphorylated after 4 hr of hypoxia and that global protein synthesis decreases in these conditions (Hantelys et al., 2019). Both activities increased after 4 hr of hypoxia and decreased at 24 hr. However LucF increased more than LucR (2.5 times versus 1.5 times, respectively). Thus the ratio LucF/LucR revealed a significant activation of the FGF1 IRES in early hypoxia, correlated to induction of endogenous FGF1 as previously shown (Hantelys et al., 2019) (Fig. 1B and 1C). Neat1 and Neat1_2 expression in cells was measured by reverse transcription and droplet digital PCR (RT ddPCR), showing an increase of Neat1 and Neat1_2 at 4 hr with a peak of expression of Neat1 at 8 hr of hypoxia, while the peak of expression of Neat1_2 was observed after 4 hr of hypoxia (Fig. 1D). The same data were also obtained by classical RT-qPCR (data not shown), in agreement with our previous report showing Neat1 induction by hypoxia in HL-1 cells (Hantelys et al., 2019).

In parallel, paraspeckle formation was studied by fluorescent *in situ* hybridization (FISH) targeting the non-coding RNA Neat1, considered as the main marker of paraspeckles. The fluorescent probes targeted either the common part of the two isoforms Neat1_1 and Neat 1_2, or only the large isoform Neat1_2 (Fig. 1E). After 4 hr of hypoxia, the number of foci increased and reached 2 foci per cell on average, while the number of cells containing at least one focus shifted from 20% to 70% (Fig. 1F-K, Fig. 1-figure supplement 2). This was observed with both Neat1 and Neat1_2 probes. The values observed at 4 hr did not change after 8 hr and 24 hr of hypoxia with the Neat1 probe (Fig. 1F-H). In contrast, the number of foci containing Neat1_2 decreased after longer times of hypoxia: at 8 hr and 24 hr, the number of foci per cell reached 1 and 0.5 while only 50% and 40% of the cells contained at least one focus, respectively (Fig. 1I-K). Surprisingly, Neat1_2 was detected in the cytoplasm in normoxia and after 24 hr of hypoxia (Fig. 1I, Fig. 1-figure supplement 2).

These data revealed that FGF1 IRES activation correlates with increased Neat1 expression and paraspeckle formation after 4 hr of hypoxia in HL-1 cardiomyocytes. To determine whether such a correlation also occurs in other cell types, similar experiments were performed in a mouse breast tumor cell line 67NR (Fig. 1-figure supplement 3). In these cells, known to be more resistant to hypoxia, Neat1 increased only after 24 hr of hypoxia. In particular, we observed a strong and significant induction of Neat1_2 (Fig. 1-figure supplement 3B). As regards the IRES activity (LucF/LucR ratio), it also increased after 24 hr of hypoxia (Fig. 1- figure supplement 3C).

These data indicate that the correlation between Neat1_2 isoform induction and IRES activation under hypoxia exists in different cell types.

### LncRNA Neat1 knock-down drastically affects the FGF1 IRES activity and endogenous FGF1 expression

To determine whether Neat1 could have a role in the regulation of FGF1 IRES activity, we depleted HL-1 for this non-coding RNA using locked nucleic acid (LNA) gapmers, antisense modified oligonucleotides described for their efficiency in knocking-down nuclear RNAs. HL-1 cells transduced with the bicistronic vector were transfected with a pool of gapmers targeting Neat1 and with a control gapmer (Supplementary file 2). The knock-down efficiency was measured by smiFISH (single molecule inexpensive FISH) and ddPCR and showed a decrease in the number of paraspeckles, correlated to the decrease of Neat1 RNA, which shifted from 5 to 2 foci per cell (Fig. 2A-B, Fig. 2-figure supplement 1A) (Tsanov *et al*. 2016). In these experiments performed in normoxia, the number of paraspeckles was high (almost 5 foci per cell), suggesting that cells were already stressed by the gapmer treatment, before being submitted to hypoxia. Alternatively, it could also be explained by the high sensitivity of the smiFISH method used here, whereas paraspeckles were detected by FISH in Figure 1.

**Figure 2.**
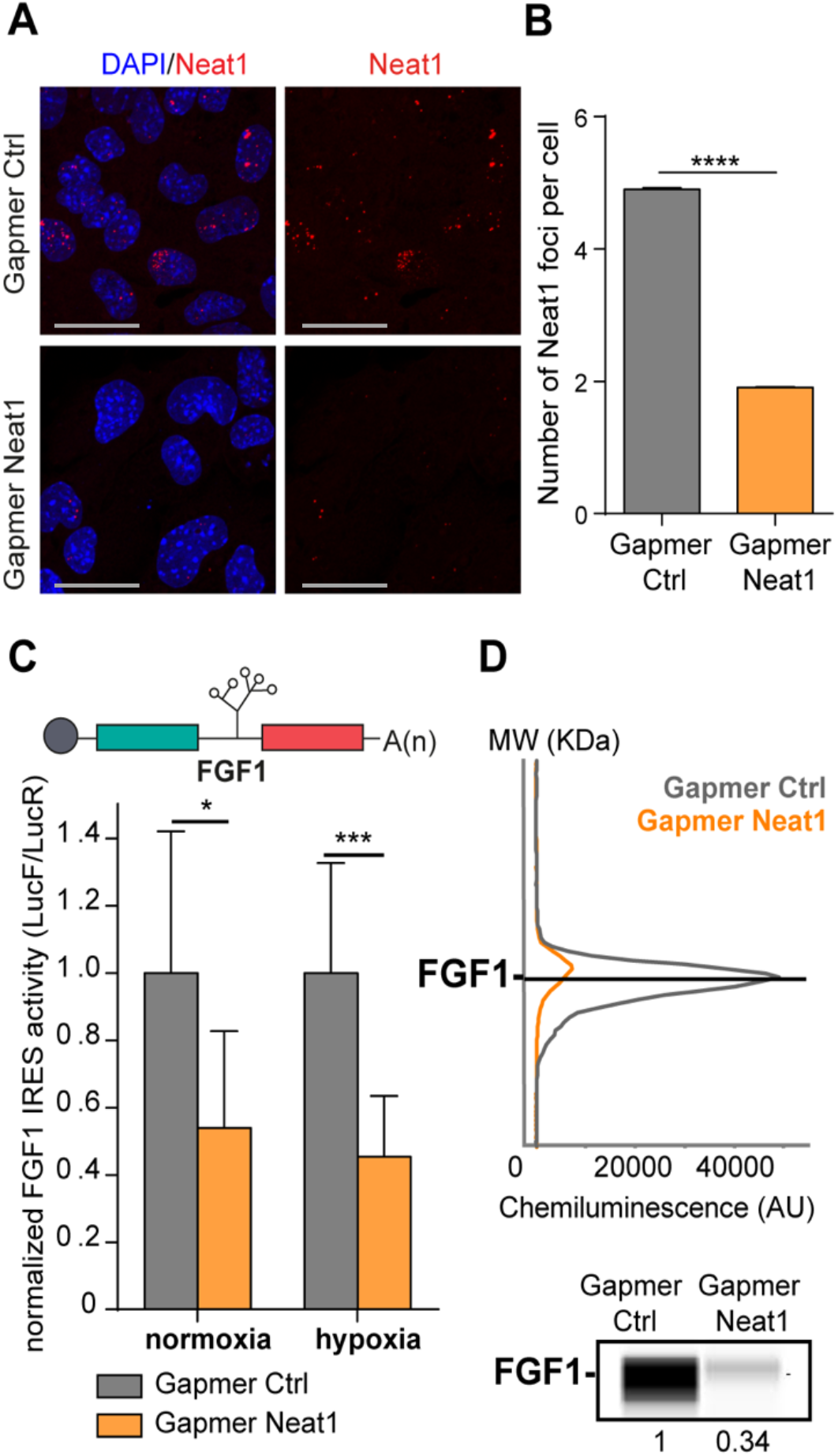
LncRNA NEAT1 knock-down drastically affects the FGF1 IRES activity and endogenous FGF1 expression. (A) SmiFISH imaging of Neat1 knock-down by a pool of LNA gapmers targeting both isoforms (Sequences in Supplementary file 2C). Cells were treated during 48 hr with the gapmers. Scale bar=10 μm (B) Neat1 foci counting per cell for the control gapmer and Neat1 LNA gapmer pool, using unpaired two-tailed student t-test with n=249 for control and 187 for Neat1 LNA gapmer. (C) FGF1 IRES activities in HL-1 cells transduced with Lucky Luke bicistronic reporter and treated with gapmer Neat1 or control during normoxia or hypoxia (1% O_2_). Histograms correspond to means + standard deviation of the mean. Non-parametric Mann-Whitney test was performed with n=9. *p<0.05, ***<0.001, ****p<0.0001. For each IRES the mean has been calculated with nine cell culture biological replicates, each of them being already the mean of three technical replicates (27 technical replicates in total). Detailed values of biological replicates are presented in Supplementary file 3. (D) Detection of endogenous mouse FGF1 by capillary Simple Western. The curve corresponds to the chemiluminescence signal detected with FGF1 antibody. A numerical blot is represented. Below the blot is shown the quantification of FGF1 normalized to total proteins and to control gapmer. Total proteins are detected by a dedicated channel in capillary Simple Western. **Figure supplement 1**. Knock-down of Neat1 and Neat1_2 in HL-1 cardiomyocytes and effect on eIF2α phosphorylation. **Figure supplement 2**. Effect of Neat1 _2 knock-down on FGF1 IRES activity. **Figure supplement 3**. FGF1 half-life in response to Neat1_2 knock-down.

To evaluate the IRES activity, the ratio LucF/LucR was measured in normoxia or after 4 hr of hypoxia, revealing that the IRES activity decreased by two times upon Neat1 depletion (Fig. 2C, Supplementary file 3). This effect was also observed on endogenous FGF1 protein expression, measured by capillary Simple Western, which decreased by three times (Fig. 2D). Neat1_2 knock-down was then performed to evaluate the contribution of the long Neat1 isoform. Data showed a decrease of FGF1 IRES activity following Neat1_2 depletion, however less important than with the knock-down of the two isoforms (Fig. 2-figure supplement 2), suggesting an involvement of both Neat1 isoforms. Capillary Western experiments indicated an increase of eIF2α phosphorylation upon Neat1_2 depletion (Fig. 2-figure supplement 1C). FGF1 half-life was superior to 24 hr and was not affected by Neat1 knock-down (Fig. 2-figure supplement 3), suggesting that Neat1 regulates FGF1 mRNA translation, directly or indirectly.

### The IRES-containing mRNA is colocalized with Neat1 during hypoxia

The effect of Neat1 on FGF1 IRES activity suggested an interaction (direct or indirect) between these two RNAs. SmiFISH experiments were performed with two sets of 48 primary probes targeting Neat1 or the bicistronic mRNA, respectively. As a control, we also used a bicistronic construct with a hairpin instead of the IRES. The two secondary probes were coupled to different fluorophores to detect Neat and the bicistronic mRNA separately and look for a putative colocalization (Fig. 3). Data clearly show that the IRES containing bicistronic mRNA is colocalized with Neat1 and that this colocalization significantly increases during hypoxia, which is not the case for the hairpin control (Fig. 3 C and D). These data suggested that the IRES-containing mRNA is recruited into paraspeckles during hypoxia.

**Figure 3.**
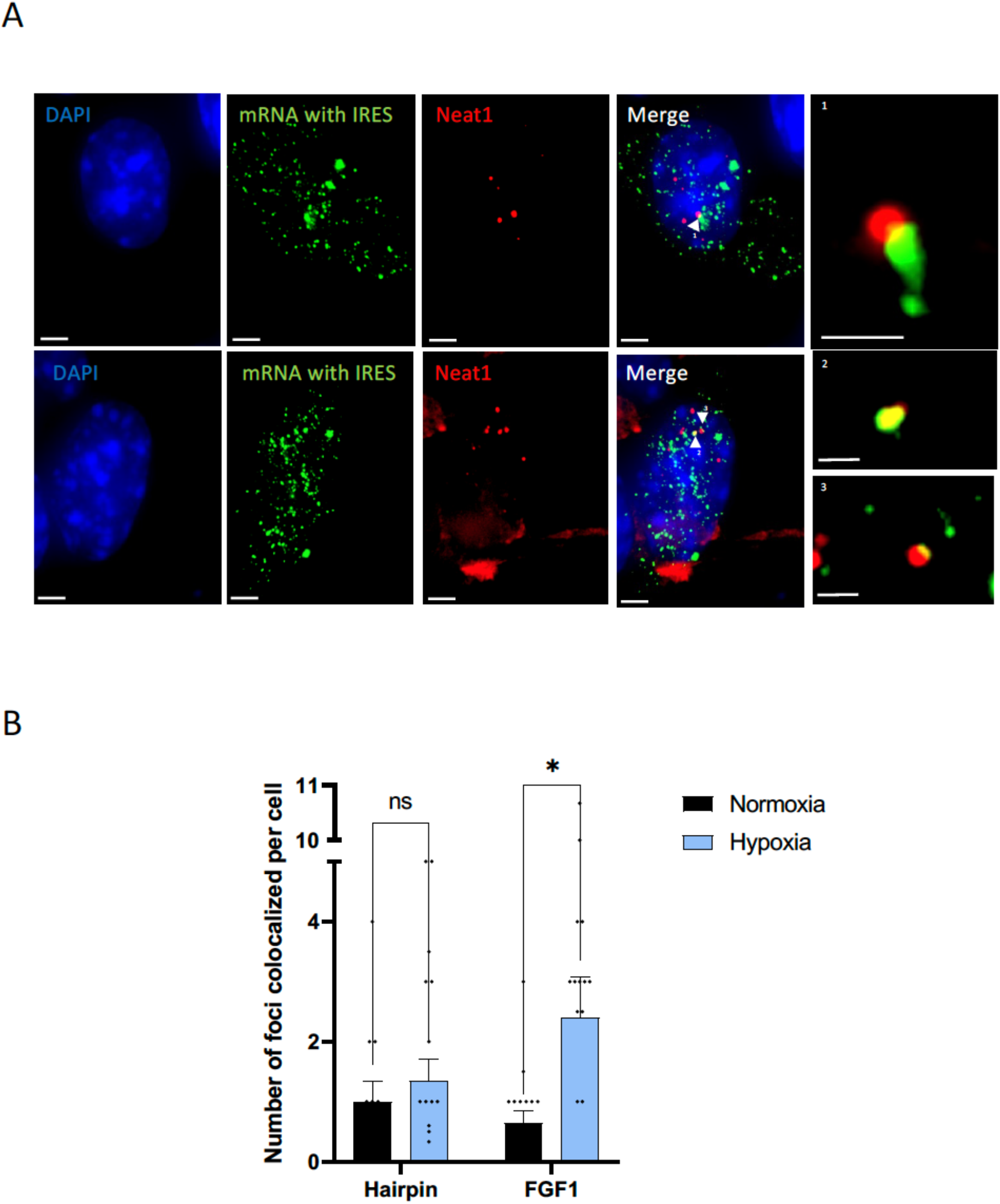
IRES-containing mRNA is colocalized with Neat1 in hypoxic HL-1 cells. Cells were transduced with lentivectors carrying bicistronic Lucky Luke constructs with the FGF1 IRES or a hairpin (control), subjected or not to 4 hr hypoxia. SmiFISH experiments were performed. (A) SmiFISH images showing the bicistronic mRNA carrying the FGF1 IRES (green) colocalized with Neat1 RNA (red) in hypoxia condition. Two representative cells are presented. (B) Quantification of colocalized spots per cell (n= 30). Unpaired two-tailed Student T-test was performed.

### Paraspeckle proteins P54^nrb^ and PSCP1, but not SFPQ, are ITAFs of the FGF1 IRES

The correlation between paraspeckle formation and FGF1 IRES activation, together with the probable recruitment of IRES-containing mRNA into paraspeckles during hypoxia, incited us to study the role of other paraspeckle components in the control of IRES activity. Three major paraspeckle proteins were chosen, the DBHS proteins, SFPQ, p54^nrb^ and PSPC1 (Fig. 4A). SFPQ and p54^nrb^ have been previously described for their ITAF function (Ainaoui et al., 2015; Cobbold et al., 2008; Lampe et al., 2018; Sharathchandra et al., 2012; Shen et al., 2017). In particular, p54^nrb^ regulates the FGF1 IRES activity during myoblast differentiation (Ainaoui et al., 2015).

**Figure 4.**
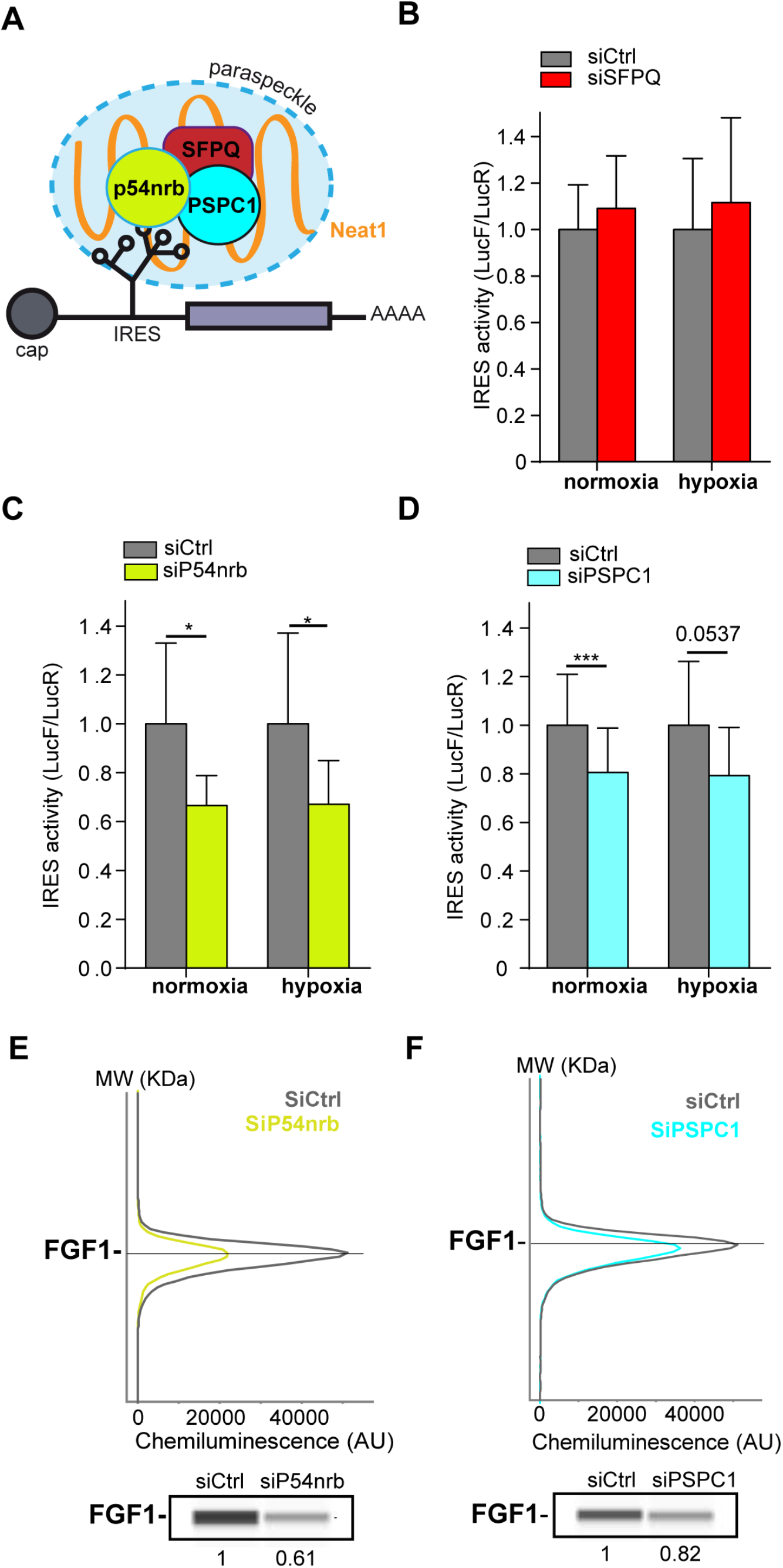
Paraspeckle proteins p54^nrb^ and PSCP1, but not SFPQ, are ITAFs of the FGF1 IRES. (A) Schema of paraspeckle and DBHS proteins possibly bound to an IRES-containing mRNA. (B-D) FGF1 IRES activity upon knock-down of SFPQ (B), P54^nrb^ (C) or PSPC1 (D) in HL-1 cell transduced with Lucky Luke bicistronic reporter during normoxia or hypoxia was measured as in Figure 2. Cells were harvested 72 hr after siRNA treatment. The IRES activity values have been normalized to the control siRNA. Histograms correspond to means + standard deviation of the mean, with a non-parametric Mann-Whitney test with n=9; *p<0.05, ***<0.001. For each IRES the mean has been calculated with nine cell culture biological replicates, each of them being already the mean of three technical replicates (27 technical replicates in total). Detailed values of biological replicates are presented in Supplementary files 3, 4 and 5. (E and F) Capillary Simple Western detection of endogenous FGF1 protein with P54^nrb^ (E) or PSPC1 (F) knockdown. **Figure supplement 1**. Knockdown of p54^nrb^, PCPC1 and SFPQ in HL-1 cardiomyocytes. **Figure supplement 2**. FGF1 half-life in response to p54^nrb^ and PSPC1 knock-down.

HL-1 cells transduced by the “Lucky Luke” bicistronic construct were transfected with siRNA smartpools targeting each of the three proteins. The knock-down efficiency was checked by capillary Simple Western, classical Western or RT qPCR (Fig. 4-figure supplement 1). SFPQ knock-down did not affect the IRES activity (Fig. 4B, Supplementary file 4). In contrast, we observed a decrease in IRES activity with p54^nrb^ and PSPC1 knock-down, both in normoxia and in hypoxia (Fig. 4C-D, supplementary files 4 and 5), despite a knock-down efficiency below 50%. p54^nrb^ and PSPC1 knock-down also inhibited the expression of endogenous FGF1 protein (Fig. 3E-F). These data confirmed the ITAF role of p54^nrb^ in HL-1 cardiomyocyte, and indicated that PSPC1 is also an ITAF of the FGF1 IRES. The ability of three paraspeckle components, Neat1, p54^nrb^ and PSPC1, to regulate the FGF1 IRES activity, together with the colocalization of the bicistronic mRNA with Neat1 observed in Figure 3, led us to the hypothesis that the paraspeckle might be involved in the control of IRES-dependent translation.

### P54^nrb^ interactome in normoxic and hypoxic cardiomyocytes

The moderate effect of p54^nrb^ or PSPC1 depletion on FGF1 IRES activity, possibly due to the poor efficiency of knock-down (> 50%), also suggested that other proteins may be involved. Previous data from the literature support the hypothesis that the IRESome is a multi-partner complex. In order to identify other members of this complex, we analysed the p54^nrb^ interactome in HL-1 cell nucleus and cytoplasm using a label-free quantitative mass spectrometry approach. For this purpose, cells were transduced by a lentivector expressing an HA-tagged p54^nrb^ (Fig. 5A). After cell fractionation (Fig. 5B), protein complexes from normoxic and hypoxic cells were immunoprecipitated with anti-HA antibody. Immunoprecipitated interacting proteins (three to four biological replicates for each group) were isolated by SDS-PAGE, in-gel digested with trypsin and analyzed by nano-liquid chromatography-tandem mass spectrometry (nanoLC-MS/MS), leading to the identification and quantification of 2013 proteins (Supplementary file 7). To evaluate p54^nrb^ interaction changes, pairwise comparisons based on MS intensity values were performed for each quantified protein between the four groups, cytoplasmic and nuclear complexes from cells subjected to normoxia or hypoxia (Fig. 5C). Enriched proteins were selected based on their significant protein abundance variations between the two compared group (fold-change (FC) > 2 and < 0.5, and Student t test P < 0.05) (see STAR Method for details) (Fig. 5D-E and Fig. 4- supplement 1). Globally, the HA-tag capture revealed an enrichment of hnRNP proteins in nucleus and of ribosomal proteins in the cytoplasm (Fig 5-supplement 1A and 1B). In nucleus P54^nrb^ interacted with itself (endogenous mouse Nono), PSPC1 and SFPQ, as well as with other paraspeckle components: in total P54^nrb^ interaction was identified with 22 proteins among 40 paraspeckle components listed in previous reports (Table 1) (Naganuma et al., 2012, Yamamoto et al., 2021). Six of these paraspeckle components exhibit an ITAF function (FUS, hnRNPA1, hnRNPK, hnRNPM, hnRNPR and SFPQ (Fig. 5-supplement 1, Table 1). Two additional ITAFs interact with p54: hnRNPC and hnRNPI (Godet et al., 2019).

**Figure 5.**
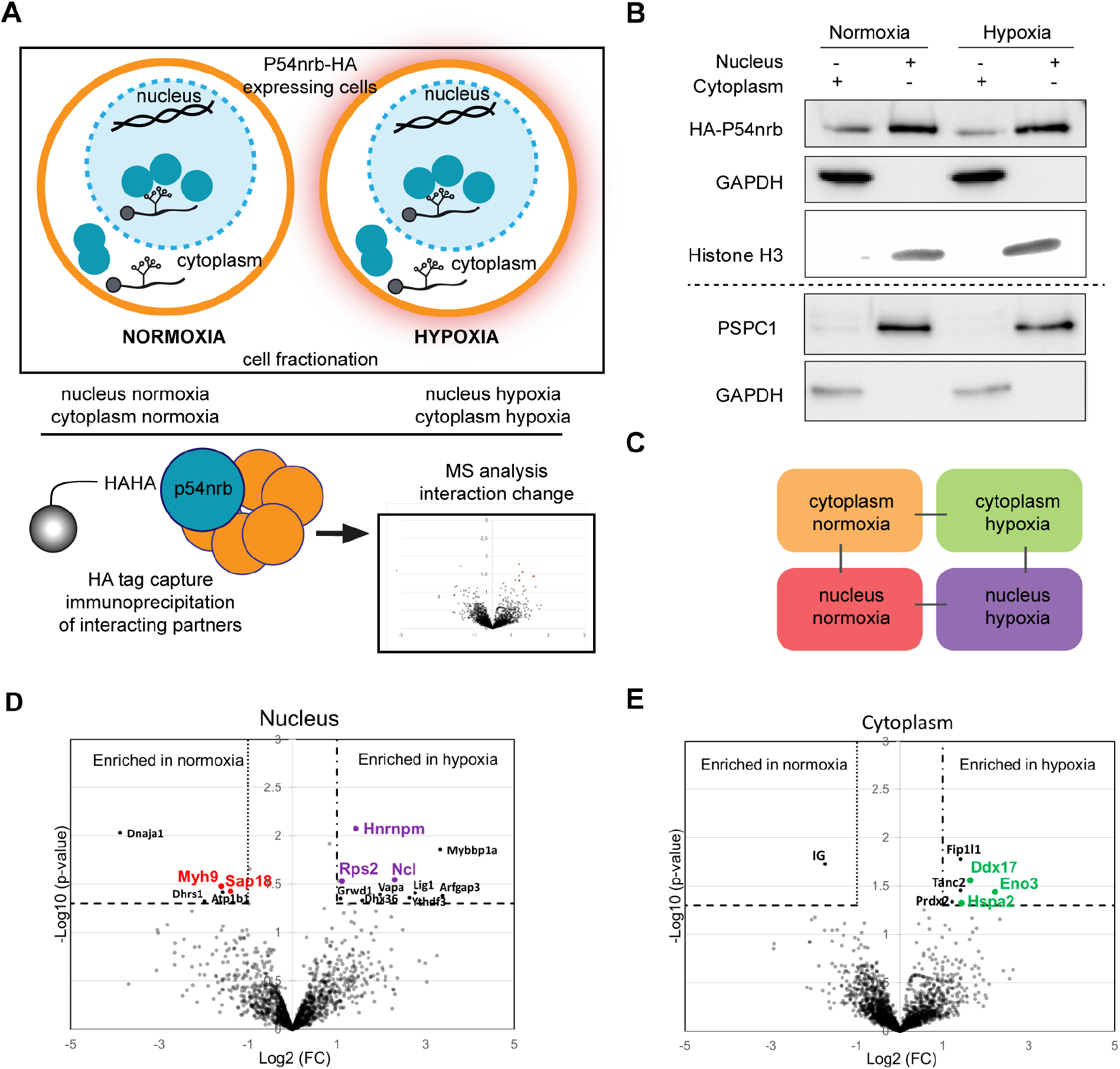
P54^nrb^ interactome in normoxic and hypoxic cardiomyocytes. (A) Experimental workflow: p54^nrb^-HA transduced HL-1 cells were subjected to normoxia or hypoxia, then nucleus and cytoplasm fractionation was performed and extracts were immunoprecipitated using anti-HA antibody. Enriched interacting proteins were identified by using a label-free quantitative mass spectrometry approach. (B) Western blot of fractionation experiment of HL-1 cells in normoxia and hypoxia. Histone H3 was used as a nuclear control and GAPDH as a cytoplasm control. The dotted line delineates two different blots of the same fractionation experiment. (C) Schema of the four pairwise comparisons submitted to statistical analysis. (D and E) Volcano plots showing proteins enriched (bold black) and significantly enriched (after elimination of false-positive hits from quantitation of low-intensity signals) in the nucleus for hypoxia (purple) versus normoxia (red) (D) or in the cytoplasm for hypoxia (green) versus normoxia (E). An unpaired bilateral student t-test with equal variance was used. Enrichment significance thresholds are represented by an absolute log2-transformed fold-change (FC) greater than 1 and a -log10-transformed (p-value) greater than 1.3. Details are provided in supplementary file 7. **Figure supplement 1**. Label-free quantitative analysis of HA-P54^nrb^-bound proteins identified by mass spectrometry in different conditions. **Figure supplement 2**. P54^nrb^ is co-immunoprecipitated by anti-nucleolin antibody.

**Table 1.**
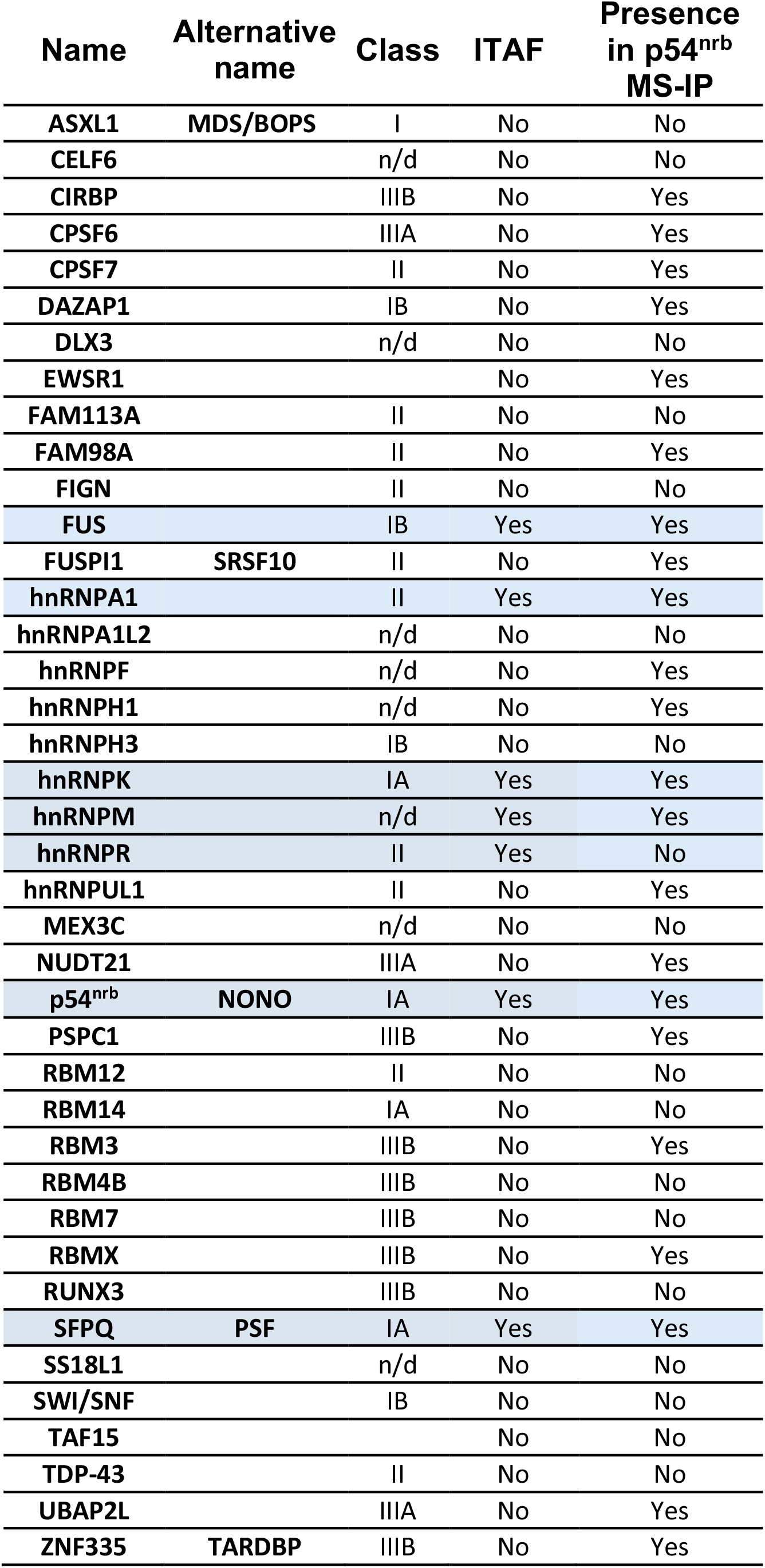
The p54^nrb^ interactome includes 22 among 40 proteins described as paraspeckle components. The paraspeckle components listed in the reports by Naganuma et al (2012) and by Yamamoto et al (2021) is presented here with their ITAF function and their presence in the p54^nrb^ interactome. Their belonging to class I, II or III of the paraspeckle proteins is indicated. Class I proteins are essential for paraspeckle formation.

As regards cytoplasmic proteins, we identified RPS25, a ribosomal protein previously described as an ITAF for many IRESs (Fig. 5-supplement 1A) (Hertz et al., 2013). Interestingly, p54^nrb^ also interacted with RPS5, RPS18 and RPS19, and other RPs, mainly from the small ribosomal subunit.

Only few proteins were significantly enriched when comparing hypoxic versus normoxic extracts. In hypoxic nucleus the significantly enriched proteins are hnRNPM, nucleolin (both previously described as ITAFs) (Hertz et al., 2013; Shi et al., 2016, 2017) and the ribosomal protein RPS2/uS5 (Fig. 5D), while the helicase Ddx17, the enolase Eno3 and the heat shock protein Hspa2 are enriched in hypoxic cytoplasm (Fig 5E). Interaction of nucleolin with p54^nrb^ was also validated by co-immunoprecipitation (Figure 5-figure supplement 2).

These data showed that p54^nrb^ interacts in normoxia and hypoxia with several ITAFs known as paraspeckle components, suggesting that the paraspeckle might be involved in the formation of the IRESome. Its interaction with numerous RPs also suggests that it interacts with the small ribosomal subunit in the cytoplasm.

### p54^nrb^-interacting proteins, nucleolin and RPS2, control the FGF1 IRES activity

The three candidates identified in nuclear extracts of hypoxic cardiomyocytes, hnRNPM, nucleolin and RPS2 represent potential candidates as ITAFs of the FGF1 IRES in hypoxia. Among them, hnRNPM has been previously described as an ITAF during myoblast differentiation while nucleolin is an ITAF of several IRESs including p53 and VEGFD IRESs but has never been described for FGF1 (Ainaoui et al., 2015; Chen et al., 2012; Godet et al., 2019; Morfoisse et al., 2016; Peddigari et al., 2013; Takagi et al., 2005).

HL-1 cardiomyocytes transduced by the Lucky Luke lentivector with the FGF1 IRES were transfected as above with siRNA smartpools targeting RPS2, hnRNPM or nucleolin (Fig. 6). The knock-down was effective, but only 50-60%, for the three mRNAs (Fig. 6A-D). This moderate knock-down was probably due to a weak transfection efficiency of HL-1 cells with the siRNAs. Nevertheless, we observed a decrease in IRES activity upon depletion of RPS2 and nucleolin, significant in normoxia but with the same trend in hypoxia while no effect was observed upon hnRNPM depletion (Fig. 6E, Supplementary file 4). Nucleolin depletion inhibited endogenous FGF1 protein expression (Fig. 6F). These data suggest that nucleolin and RPS2 are new ITAFs of the FGF1 IRES. Their nuclear localization and interaction with p54^nrb^ indicate that they could be components of the paraspeckle. RPS2 has never been described as an ITAF before the present study.

**Figure 6.**
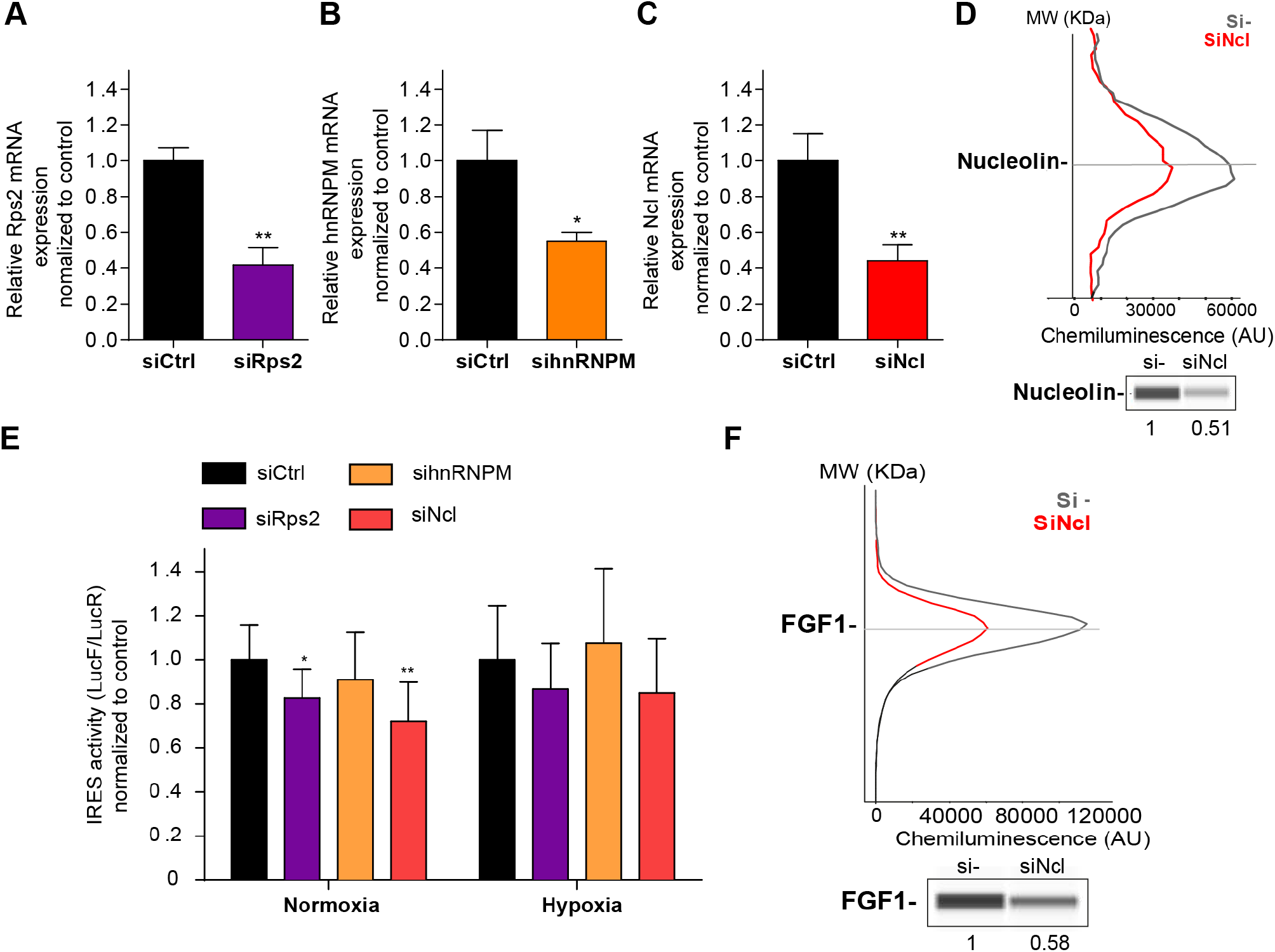
p54^nrb^-interacting proteins, nucleolin and Rps2, control the FGF1 IRES activity. (A-C) Quantification of Rps2 (A), hnRNPM (B) and nucleolin (C) RNA expression in HL-1 cells transfected with siRNAs against Rps2, hnRNPM or nucleolin, respectively. RNA expression was measured by RT-qPCR and normalized to control siRNA. One representative experiment is shown with n=3 biological replicates. Student two-tailed t-test was performed with n=3 of for E Mann-Whitney test with n=9; *p<0.05, **p<0.01, ***<0.001, ****p<0.0001. (D) Capillary Simple Western of nucleolin following nucleolin knock-down. (E) FGF1 IRES activity with knock-down by siRNA interference of candidate ITAF nucleolin in HL-1 in normoxia or hypoxia 1% O_2_ was performed as in Figure 2. The IRES activity values have been normalized to the control siRNA. Histograms correspond to means + standard deviation of the mean, with a non-parametric Mann-Whitney test *p<0.05, **p<0.01. For each IRES the mean has been calculated with nine cell culture biological replicates, each of them being already the mean of three technical replicates (27 technical replicates in total but the M-W test was performed with n=9). Detailed values of biological replicates are presented in Supplementary file 6. (F) Capillary Simple Western of endogenous FGF1 following nucleolin knock-down. Histograms correspond to means + standard deviation.

### Neat1 is the key activator of (lymph)angiogenic and cardioprotective factor mRNA IRESs

We have shown above that three main paraspeckle components, Neat1, p54^nrb^ and PSPC1, control the FGF1 IRES activity in HL-1 cardiomyocytes. To determine if the role of paraspeckle in translational control may be generalized to other IRESs, we used Lucky Luke lentivectors containing a set of other IRESs between the two luciferase genes (Fig. 7). HL-1 cells were transduced by the different lentivectors and transfected either by the siRNA smartpools to deplete p54^nrb^ and PSPC1, or by the gapmer pool to deplete Neat1. The data revealed that p54^nrb^ or PSPC1 depletion affected several IRESs but not all (Fig. 6A-B, Supplementary files 4 and 5), whereas Neat1 depletion clearly affected all cellular IRESs but not the viral EMCV IRES (Fig. 7C, Supplementary file 3).

**Figure 7.**
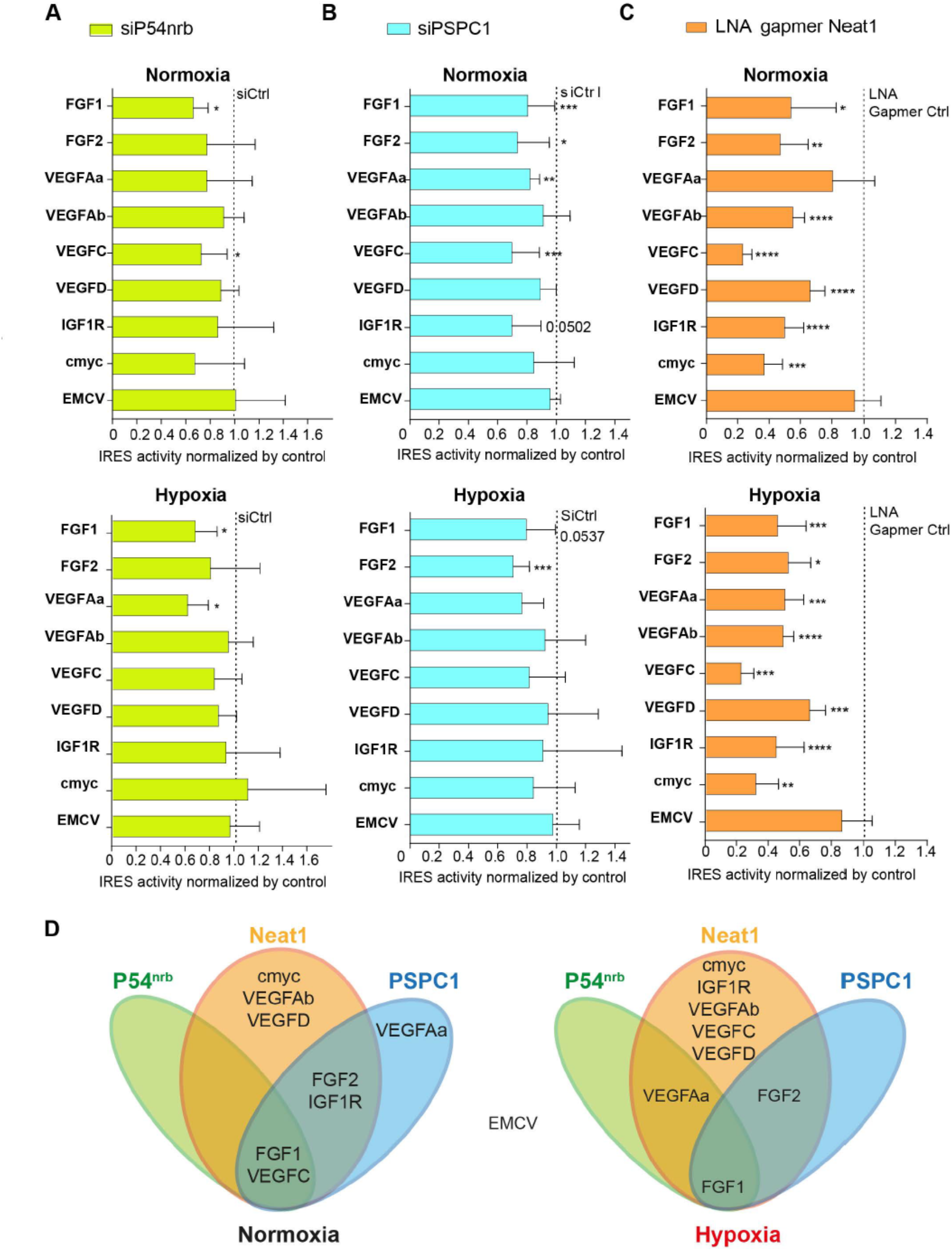
Neat1 is the key activator of (lymph)angiogenic and cardioprotective factor mRNA IRESs. (A-C) HL-1 subjected to normoxia or 1% O_2_ hypoxia were transduced by Lucky Luke bicistronic lentivectors with FGF1, FGF2, VEGFAa, VEGFAb, VEGFC, VEGFD, IGF1R, c-myc or EMCV IRES, then the knock-down of p54^nrb^(A) PCPC1 (B) and Neat1 (C) was performed as in Fig. 2 and Fig. 4. IRES activities were measured and normalized to activities in normoxia. IRES activity in normoxia is represented by a dotted line at 1. Histograms correspond to means + standard deviation, and Mann-Whitney test with n=9 or n=12 for FGF1 IRES; *p<0.05, **p<0.01, ***<0.001, ****p<0.0001. For each IRES the mean has been calculated with nine cell culture biological replicates, each of them being already the mean of three technical replicates (27 technical replicates in total). Detailed values of biological replicates are presented in Supplementary files 3, 4 and 5. (D) Schema depicting groups of IRESs regulated by Neat1, PSPC1 or P54nrb in normoxia or hypoxia.

These data allowed us to group the IRESs in different “regulons” in normoxia and in hypoxia (Fig. 7D). According to our data, P54^nrb^ is an activator of the FGF1 and VEGFC IRESs in normoxia, and of the FGF1 and VEGFAa IRESs in hypoxia. PSPC1 is an activator of the FGF1, FGF2, VEGFAa, VEGFC and IGF1R IRESs in normoxia and of the FGF1 and FGF2 IRESs in hypoxia. Neat1 is an activator of the FGF1, FGF2, VEGFAb, VEGFC, VEGFD, IGF1R and c-myc IRESs but not of the VEGFAa IRES in normoxia while it activates all the cellular IRESs in hypoxia. The EMCV IRES does not belong to any of these groups as it is not regulated by these three ITAFs, suggesting that this viral IRES is not regulated by the paraspeckle.

In conclusion, these data suggest that IRESome composition varies for each IRES and with the normoxic or hypoxic conditions. The long non-coding RNA Neat1 appears as the key ITAF for the activation of all the cellular IRESs, suggesting a crucial role of the paraspeckle in IRESome formation and in the control of IRES-dependent translation, at least for cellular IRESs.

### Neat1 isoforms impact the recruitment into polysomes of mRNAs involved in the stress response

The role of Neat1 on translatome was then studied using a Fluidigm Deltagene PCR array targeting 96 genes coding IRES-containing mRNAs, ITAFs or proteins involved in angiogenesis and cardioprotection (Supplementary file 2E). HL-1 cells were treated with gapmers targeting the two Neat1 isoforms or only Neat1_2 before analyzing the recruitment of mRNAs into polysomes compared to the control gapmer. Recruitment into polysomes decreased for 49% of IRES-containing mRNAs following Neat1 invalidation, and increased for the other 51%. In contrast this decrease concerned 95% of these mRNAs after Neat1_2 knockdown (Fig. 8A and 8B, Supplementary file 8). In contrast, the global level of translation was not affected (Fig. 8- figure supplement 1). This suggested that Neat1 has an effect on translation of most IRES-containing mRNAs, but that the two isoforms may have distinct effects. Interestingly, a similar effect was observed for the other genes tested in the PCR array: Neat1 or Neat1_2 knock-down inhibited translation of ITAF-coding genes by 71% or 87%, respectively (Fig. 8C and 8D, Supplementary file 8). This inhibition concerned 57% or 89% of the remaining genes involved in angiogenesis and cardioprotection for Neat1 or Neat1_2 knock-down, respectively (Fig 8- figure supplement 2). In total, 92% of the genes of the PCR array were less recruited into polysomes after Neat1_2 knock-down, versus only 56% after Neat1 knock-down. These data strongly suggest that Neat1_2 might be a translational activator of families of genes involved in the response to hypoxic stress in cardiomyocytes.

**Figure 8.**
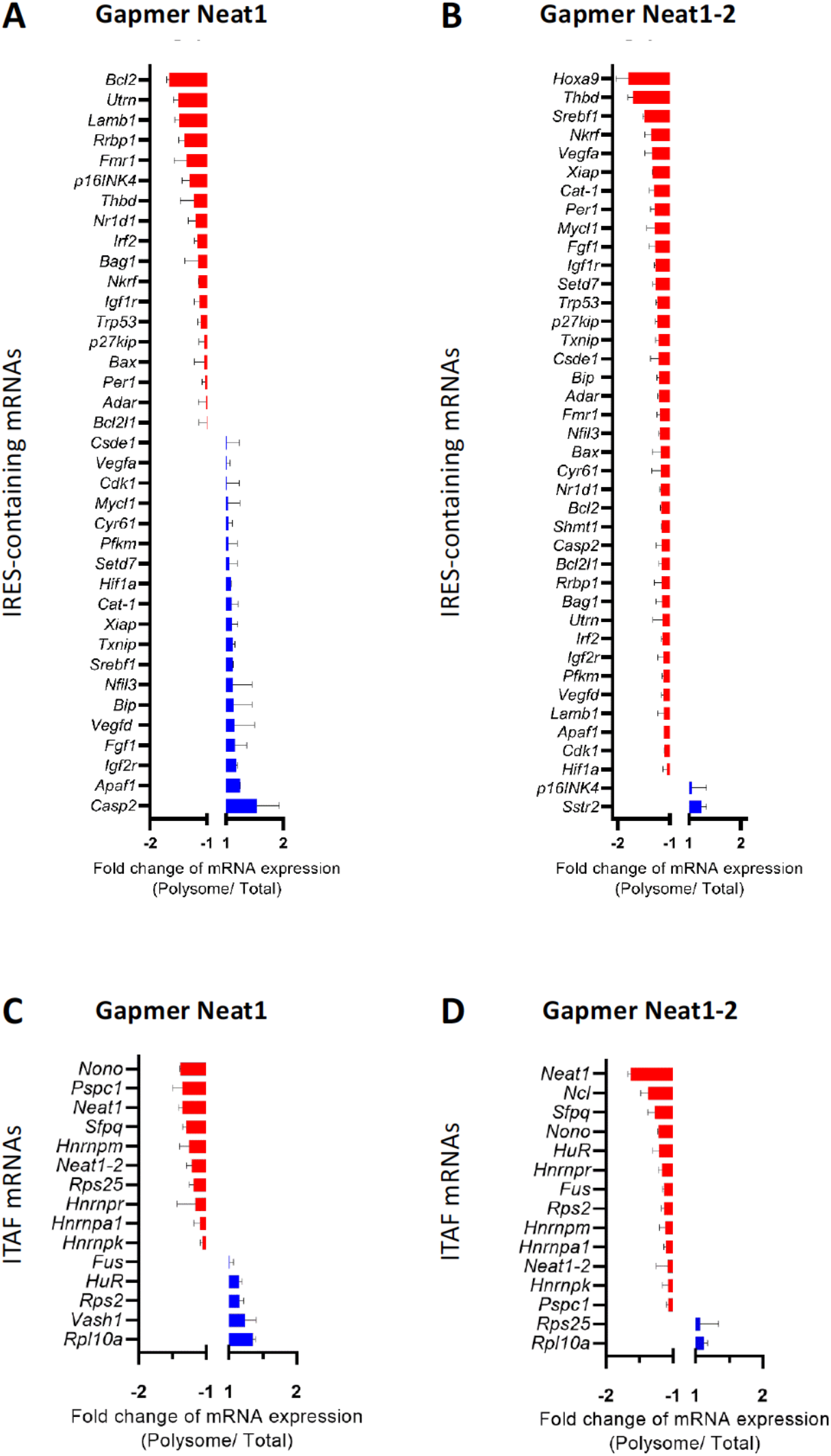
Neat1_2 knock-down down-regulates translation of most IRES-containing RNAs as well as mRNAs coding ITAFs. HL-1 cardiomyocytes were transfected with gapmer Neat1, Neat1_2, or control. Polysomes were purified on sucrose gradient as described in Star Methods. The polysome profile is presented in Fig. 8-figure supplement 1. RNAs were purified from cytoplasmic extracts and from pooled polysomal fractions and analyzed on a Fluidigm deltagene PCR array from two biologicals replicates (cell culture dishes and cDNAs), each of them measured in three technical replicates (PCR reactions) (Supplementary file 8). IRES-containing mRNAs (A-B) and ITAF mRNA levels in polysomes (C-D) (polysomal RNA/ total RNA) were analyzed. Relative quantification (RQ) of mRNA level was calculated using the 2– ΔΔCT method with normalization to GAPDH mRNA and to HL-1 tranfected by gapmer control, and is shown as fold change of repression (red) or induction (blue). **Figure-supplement 1**. Polysome profiles of HL-1 cardiomyocytes treated by gapmer Neat1 or Neat1_2, compared to gapmer control. **Figure-supplement 2**. Effect of Neat-1 and Neat1_2 knockdown on translation of mRNAs coding angiogenic and cardioprotective factors.

## DISCUSSION

The present data demonstrate a link between the paraspeckle and the control of IRES-dependent translation during hypoxia in mouse cardiomyocytes. We show that three major paraspeckle components regulate IRES-dependent translation: p54^nrb^, PSPC1 and Neat1, as well as by two proteins present in the p54^nrb^ nuclear interactome, nucleolin and RPS2. Neat1 appears as the key to this paraspeckle-related activation of translation in response to hypoxia. This lncRNA is an activator of all cellular IRESs tested, but not of the viral EMCV IRES. More broadly, Neat1 isoforms impact the recruitment into polysomes of most IRES-containing mRNAs and several families of mRNAs involved in the response to hypoxia. The colocalization of IRES-containing mRNA with Neat1 RNA in paraspeckles increased in hypoxia conditions, suggesting that the paraspeckle may be a recruitment platform for IRES-containing mRNAs during stress and that the IRESome could be assembled in the paraspeckle before mRNA export from the nucleus.

It may be noted that the inhibition of IRES activities resulting from ITAF depletion is quite moderate for the different proteins while stronger for the lncRNA Neat1. This cannot be explained only by differences in knock-down efficiency. We hypothesize is that several proteins are present in the IRESome complex and that there may be a certain redundancy between them. Thus, the depletion of a single ITAF would not be sufficient to abolish the IRES activity completely. Also, to explain why paraspeckle ITAFs such as p54^nrb^ and PSPC1 do not inhibit all the IRESs, we propose that the paraspeckle IRESome protein composition varies depending on the IRES and the hypoxic or normoxic condition, while Neat1 remains the main actor of the process. Several observations suggest that Neat1_2 may be the main isoform involved. However, the knock-down of Neat1-2 isoform with a specific gapmer does not affect IRES activity as much as the knock-down of both Neat1 isoforms (Fig. 2, Figure supplement 2). We were not successful in knocking down the isoform Neat1_1, as its sequence is entirely contained in Neat1_2. Thus at this stage we conclude that the two isoforms are probably involved. The fluidigm PCR array suggests that they may affect translation differently (Figure 8).

We searched for an ITAF able to regulate a set of IRESs during hypoxia and found the lncRNA Neat1 as a wide activator of IRES-dependent translation. However, our data show that Neat1 also regulates IRES activities both in normoxia and hypoxia. One explanation may be that Neat1 is already expressed in normoxia in HL-1 cells, which are transformed cells despite their cardiomyocyte beating phenotype (Claycomb et al., 1998). Although Neat1 expression and paraspeckle number increase in response to hypoxia, a significant percentage of cells already contain paraspeckles in normoxia, which may explain why IRESs are already active in normoxia. It has been reported that Neat1_2 is not expressed in all tissues in vivo, whereas it is found in all transformed or immortalized cell lines (data not shown) (Nakagawa et al., 2011). In concordance with this observation, previous reports show that cellular IRESs are active in all cultured cell lines while inactive or tissue-specific in mice (Créancier et al., 2000, 2001).

Our data contrast with the study of Shen *et al*. who showed that Neat1 depletion allows redistributing p54^nrb^ and SFPQ/PSF onto the c-myc mRNA, in correlation with an increase in c-Myc protein (Shen et al., 2017). Several reasons may explain this lack of concordance. Firstly, different cell lines were used: HL-1 cardiomyocytes and 67NR breast tumor cells in the present study, HeLa and MCF7 tumor cells in the report by Shen *et al*. The regulation of IRES-dependent translation varies depending on cell lines. Secondly, they worked with human cell lines while our report is focused on mouse cells. In human, c-myc expression is different from mouse as the c-myc gene contains an additional upstream promoter, P0, which generates a longer transcript with a second IRES (Nanbru et al., 2001). Thirdly, they have not directly analyzed the c-myc IRES activity but only the binding of p54^nrb^ and SFPQ to the c-myc endogenous mRNA. Moreover an increase in c-myc protein expression does not necessarily correspond to increased IRES activity as the c-myc mRNA is also translated by the cap-dependent mechanism (Nanbru et al., 1997). Taken together, the two studies are different rather than discordant.

A surprising result has been finding a ribosomal protein, RPS2, in the nuclear p54^nrb^ interactome. This suggests an extra-ribosomal role of this protein. Its interaction with p54^nrb^ favors the hypothesis that RPS2 would impact the IRES activity as an IRESome component in the paraspeckle. The presence of nucleolin in the complex also suggests a link of paraspeckle with nucleolus and ribosome biogenesis. Supporting this, PSPC1 was first identified in the nucleolus proteome (Fox et al., 2002). The nuclear binding of specific ribosomal proteins to IRESs might be a mechanism for forming specialized ribosomes.

Neat1 is not the first lncRNA to exhibit an ITAF function. The lncRNA TP53-regulated modulator of p27 (TRMP) has been recently described as an ITAF of the p27^kip^ IRES (Yang et al., 2018). TRMP inhibits the p27^kip^ IRES activity by competing with the IRES for pyrimidine tract binding protein (PTB) binding and prevents IRES activation mediated by PTB. Also, the lncRNA ARAP-as1 directly interacts with SFPQ, which results in release of PTB and activation of c-myc IRES (Zhang et al., 2019). We have not yet deciphered the mechanism of action of Neat1. We propose that the paraspeckle would be a recruitment platform for IRES-containing mRNAs. Neat1, by interacting with p54^nrb^ and other paraspeckle proteins/ITAFs, would thus allow IRESome formation in the paraspeckle. Is the role of Neat1 exclusively nuclear in the paraspeckle, or is it exported to the cytoplasm with the IRESome complex? Several observations argue for the presence of Neat1 in the cytoplasm: our FISH experiments clearly identify the Neat1-2 isoform in the cytoplasm (Fig.1, figure supplement 2), while a recent report shows that Neat1-1 isoform is released from nucleus to cytoplasm where it suppresses the Wnt signaling in leukemia stem cells and acts as a tumor suppressor in acute myeloid leukemia (Yan et al., 2021). Neat1-2 isoform has been detected in the cytoplasm of hematopoietic cells by other authors. Interestingly, they identified a histone modifier, ASXL1, interacting with p54^nrb^/NONO and involved in paraspeckle formation. Mutation of ASXL1 generates Neat1_2 export to the cytoplasm (Yamamoto et al., 2021). Furthermore, the role of cytoplasmic Neat1 in translation is suggested by our previous data showing that Neat1 is present in HL-1 cell polysomes and that this association with polysomes is increased in early hypoxia (Hantelys et al., 2019). The involvement of Neat1 in translation control via a cytoplasmic location is also supported by the presence of the triple helix in the 3’UTR of Neat1_2, whose role in translation activation has been demonstrated (Wilusz et al., 2012).

The model of IRESome formation mediated by Neat1 in the paraspeckle, and the absence of any impact of Neat1 on the picornaviral EMCV IRES activity, are both consistent with previous reports suggesting that the site of mRNA synthesis is crucial for IRES structure and function (Semler & Waterman, 2008). For picornaviruses whose mRNAs are synthesized in the cytoplasm, IRES elements would be able to form an IRESome RNP in the cytoplasm. In contrast, cellular mRNAs (as well as DNA viruses and retroviruses mRNAs) transcribed in the nucleus need a nuclear event (Ainaoui et al., 2015; Stoneley et al., 2000). The present data provide a mechanism for this nuclear history and reveal a new function of the paraspeckle, a nuclear body, in IRESome formation.

A role of Neat1 in ischemic heart has been recently reported showing that Neat1 downregulation would protect cardiomyocytes from apoptosis by regulating the processing of pri-miR-22 (Gidlöf et al., 2020). Surprisingly, these authors show that hypoxia down-regulates Neat1 expression in cardiomyocytes. This contradicts our data showing that Neat1 is induced by hypoxia. Our data are however in agreement with the rest of the literature showing that Neat1 is induced by hypoxia in tumors, its transcription being activated by HIF-2 (Choudhry et al., 2015). Another study also showed that Neat1 overexpression protects cardiomyocytes against apoptosis by sponging miR125a-5p, resulting in upregulation of the apoptosis repressor gene B-cell lymphoma-2-like 12 (BCL2L12) (Yan et al., 2019). These contradictory reports highlight the complex impact of Neat1 on miRNA-mediated gene regulation.

In the present study, we have uncovered a novel role of Neat1 in the translational control of several families of genes involved in stress response, angiogenesis and cardioprotection, while it does not affect global translation. The increased protein synthesis from mRNAs coding ITAFs favors a wide role of Neat1 and of the paraspeckle in activating IRES-dependent translation. Many of the genes involved in angiogenesis or cardioprotection tested here have not been described as containing an IRES in their mRNAs. We can make the hypothesis that these mRNA families either contain IRESs that have not been identified yet, or are translated by another cap-independent mechanism such as m6A-induced ribosome engagement sites (MIRES) (Prats et al., 2020).

Neat1, as a stress-induced lncRNA, plays a role in many pathologies including cancer and ischemic diseases, thus its central role in the translational control of expression of genes involved in tissue revascularization and cell survival makes it a potential therapeutic target of great interest.

## MATERIALS AND METHODS

### KEY RESOURCE TABLE

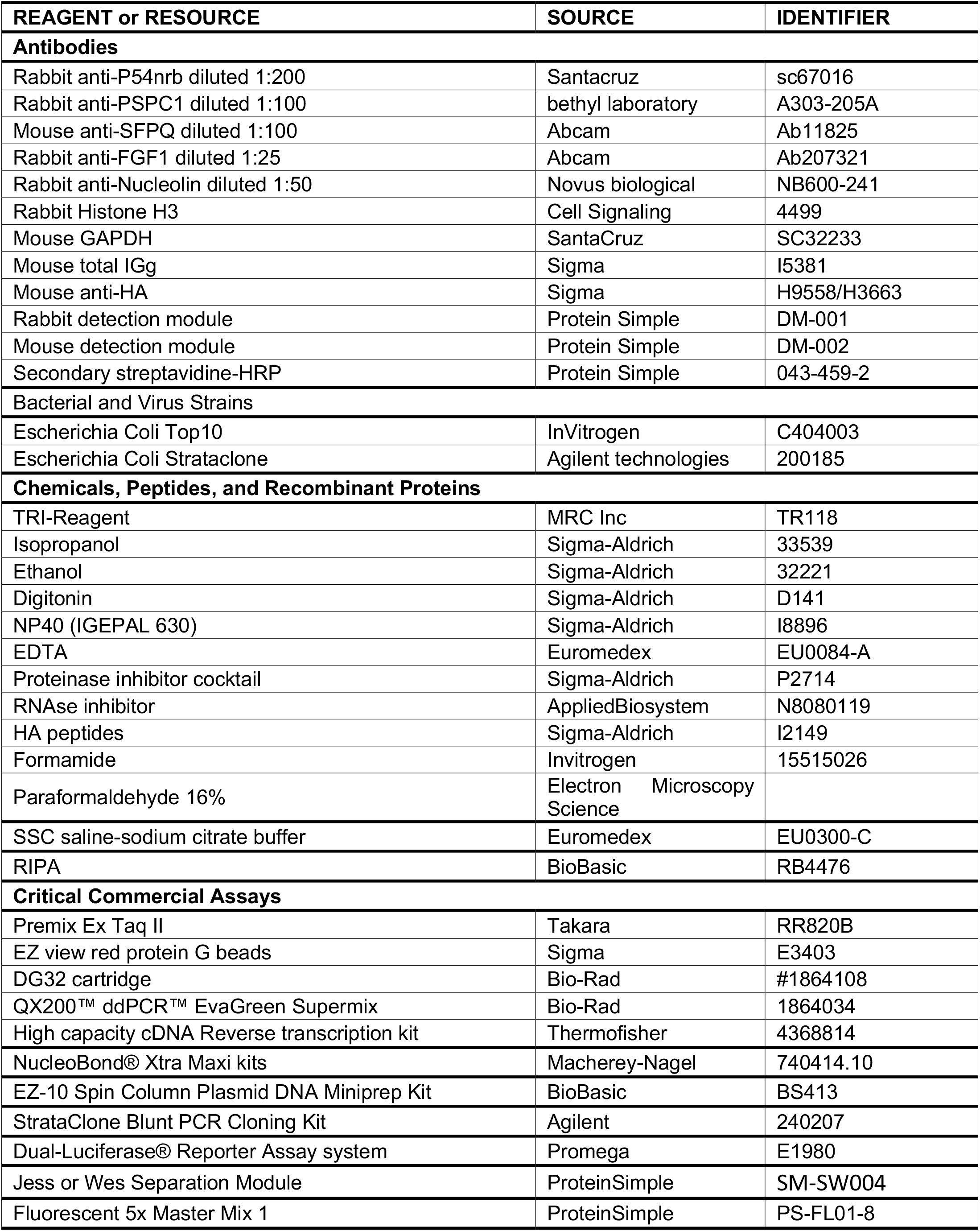
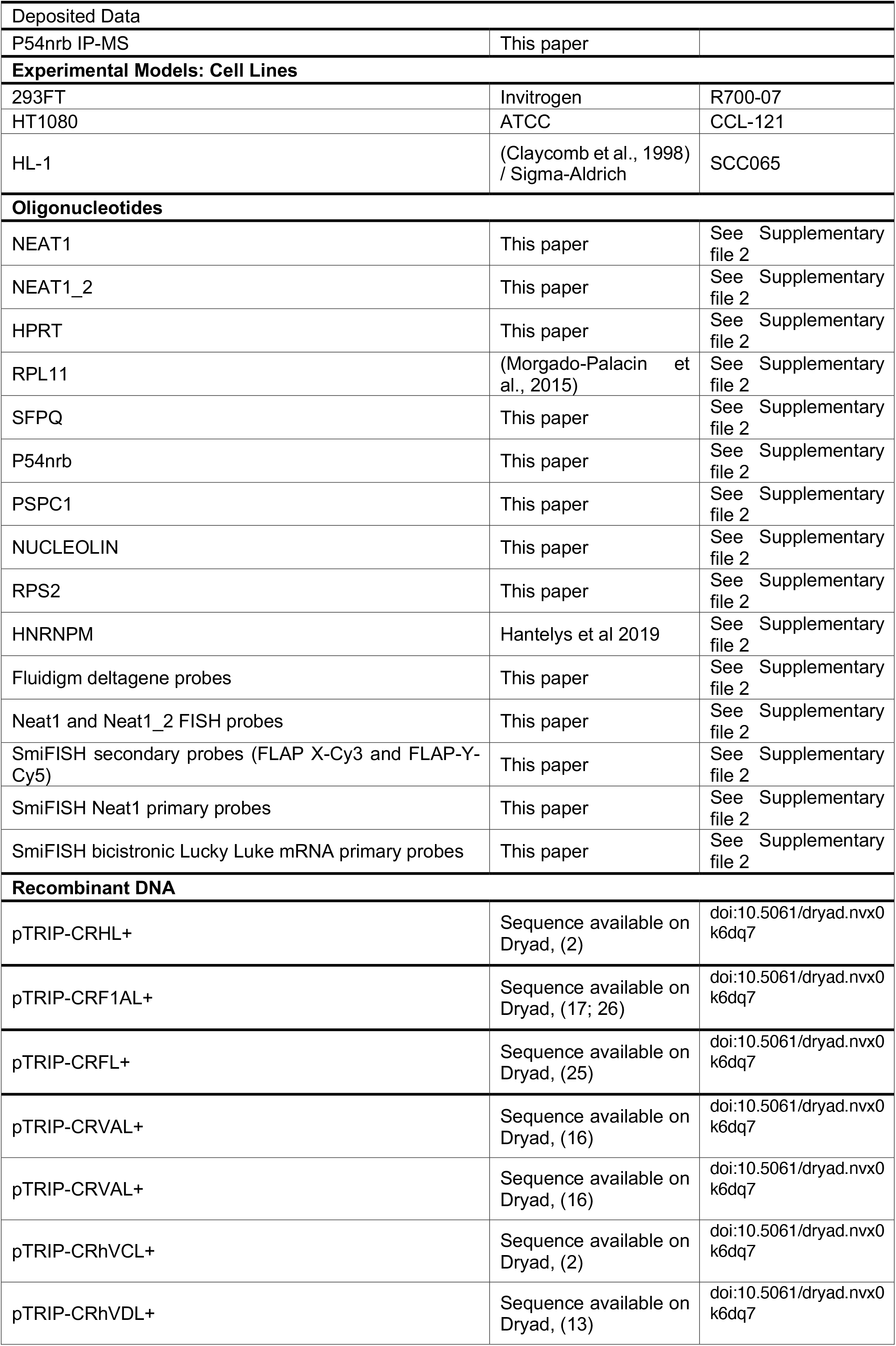
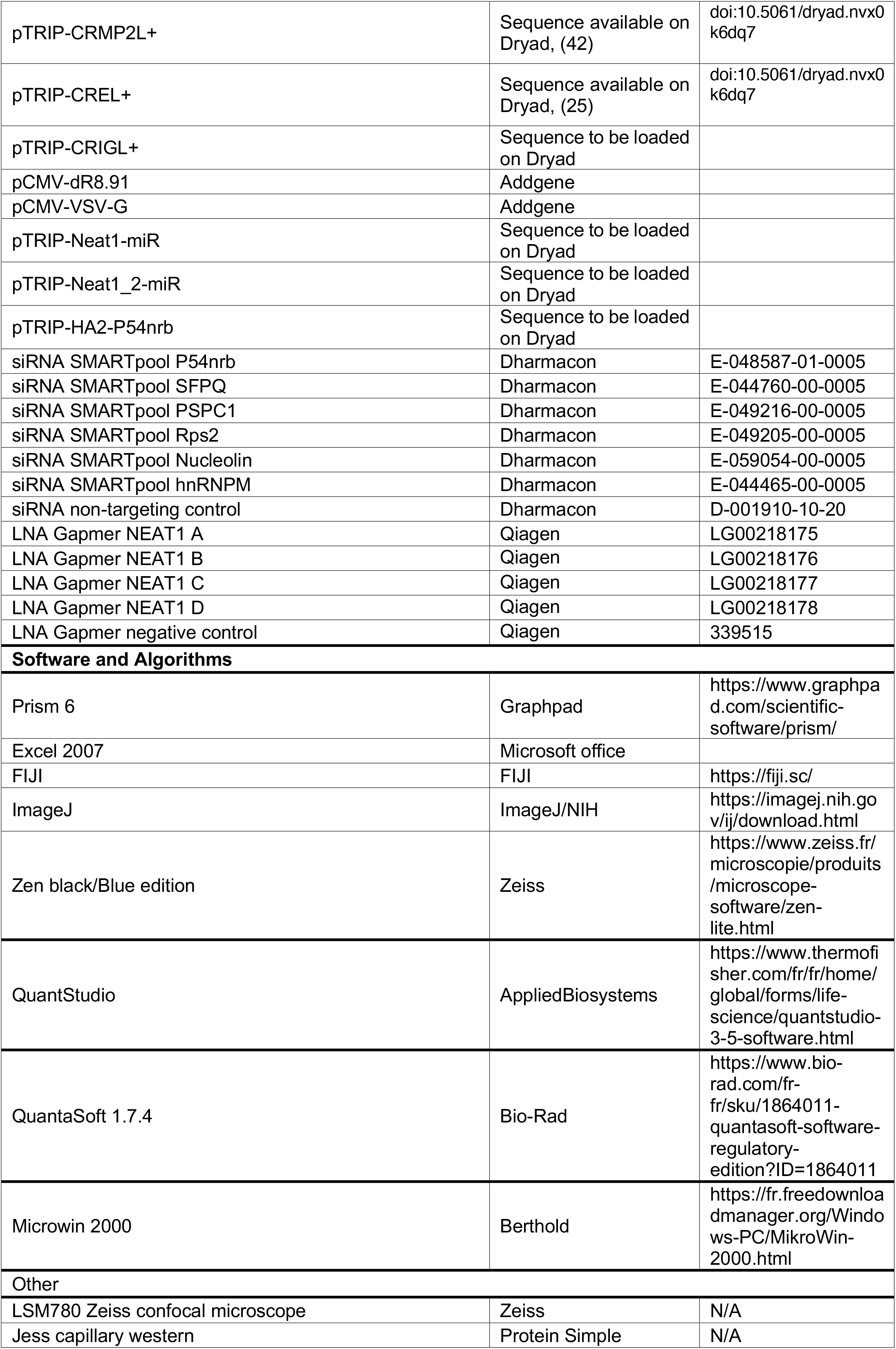
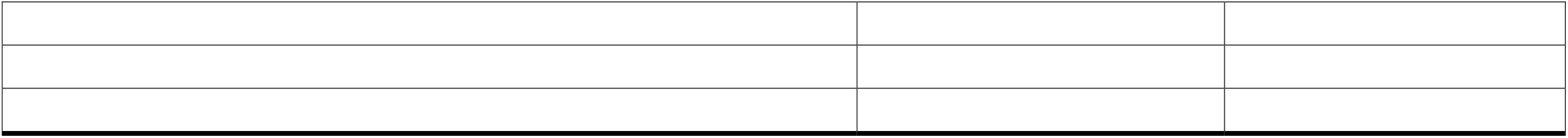

### LEAD CONTACT AND MATERIALS AVAILABILITY

- Further information and requests for resources and reagents should be directed to and will be fulfilled by the Lead Contact, Anne-Catherine Prats.
- This study did not generate new unique reagents.

### EXPERIMENTAL MODEL AND SUBJECT DETAILS

#### Cell lines

Female human embryonic kidney cells HEK-293FT and male human fibrosarcoma HT1080 cells were cultured in DMEM-GlutaMAX + Pyruvate (Life Technologies SAS, Saint-Aubin, France), supplemented with 10% fetal bovine serum (FBS), and MEM essential and non-essential amino acids (Sigma-Aldrich). Female mouse atrial HL-1 cardiomyocytes (Sigma-Aldrich) were cultured in Claycomb medium containing 10% FBS, Penicillin/Streptomycin (100U/mL-100μg/mL), 0.1 mM norepinephrine, and 2 mM L-Glutamine. Cell culture flasks were precoated with a solution of 0.5% fibronectin and 0.02% gelatin for 1 hr overnight at 37°C (Sigma-Aldrich). To keep HL-1 phenotype, cell culture was maintained as previously described (Claycomb et al., 1998). All cells were cultured in a humidified chamber at 37°C and 5% CO_2_. When subjected to hypoxia, cells were incubated at 37°C at 1% O_2_.

#### Bacterial strains

- Top 10 Escherichia coli (InVitrogen, thermofisher scientific C404003)
- Strataclone Escherichia coli (Agilent technologies, 200185)

These cells were stored at **-**80°C and grown in LB medium at 37°C. Top10 cells were used for plasmid amplification of pTRIP lentivector. Strataclone cells were used for recombination and amplification of PCR product into pSC-B-amp/kan plasmid.

### METHOD DETAILS

#### Cell transfection

siRNA treatment on transduced cells was performed 72 hr after transduction (and after one cell passage) in 24 well plate for reporter activity assay or 12 well plate for gene expression experiment. HL-1 were transfected by siRNAs as follows: one day after being plated, cells were transfected with 10 nM of small interference RNAs from Dharmacon Acell SMARTpool targeting P54^nrb^, PSPC1, SFPQ, hnRNPM, Nucleolin, Rps2, or non-targeting siRNA control (siControl), using INTERFERin (Polyplus Transfection) according to the manufacturer’s recommendations, in DMEM-GlutaMAX + Pyruvate media without penicillin-streptomycin. The media was changed 24 hr after transfection and the cells were incubated 72 hr for the time of transfection at 37°C with siRNA For Neat1 knockdown, HL-1 cells were transduced with a pool of 4 gapmers (Qiagen) at 40nM (10Nm each) and incubated 48h after transfection, proceeded essentially as described above (siRNA and gapmer sequences are provided in Supplementary file 2).

#### Cell transduction

For lentivector transduction, HL-1 cardiomyocytes were plated into a T25 flask and transduced overnight in 2.5 mL of transduction medium (OptiMEM-GlutaMAX, Life Technologies SAS) containing 5 μg/mL protamine sulfate in the presence of lentivectors (MOI 2). HL-1 cells were transduced with an 80-90% efficiency in the mean.

#### Lentivector construction

Bicistronic lentivectors coding for the renilla luciferase (LucR) and the stabilized firefly luciferase Luc+ (called LucF in the text) were constructed from the dual luciferase lentivectors described previously, which contained Luc2CP (Morfoisse et al., 2014, 2016). The LucR gene used here is a modified version of LucR where all the predicted splice donor sites have been mutated. The cDNA sequences of the human FGF1, -2, VEGFA, -C, -D, c-myc and EMCV IRESs were introduced between the first (LucR) and the second cistron (LucF) (Giraud et al. 2001; Nanbru et al., 1997; Prats et al., 2013; Vagner et al., 1995). IRES sequence sizes are : 430 nt (FGF1), 480 nt (FGF2), 302 nt (VEGFAa), 485 nt (VEGFAb), 419 nt (VEGFC), 507 nt (VEGFD), 363 nt (c-myc), 640 nt (EMCV), 973 nt (rat IGF1R) (Huez et al., 1998; Martineau et al., 2004; Morfoisse et al., 2014, 2016; Nanbru et al., 1997; Vagner et al., 1995).The two IRESs of the VEGFA have been used and are called VEGFAa and VEGFAb, respectively (Huez et al., 1998). The hairpin negative control contains a 63 nt long palindromic sequence cloned between LucR and LucF genes (Hantelys et al., 2019). This control has been successfully validated in previous studies (Créancier et al., 2000; Morfoisse et al., 2014).The expression cassettes were inserted into the SIN lentivector pTRIP-DU3-CMV-MCS vector described previously (Prats et al., 2013). All cassettes are under the control of the cytomegalovirus (CMV) promoter. Vector sequences are available on Dryad (doi:10.5061/dryad.nvx0k6dq7). The lentivectors coding artificial miRNAs miR-Neat1 and miR-Neat1_2 were constructed by inserting double-stranded oligonucleotides targeting Neat1 or Neat1_2, according to a protocol adapted from the BLOCK-iT™ technology of Life Technologies (sequences provided in Supplementary file 2). Plasmid construction and amplification were performed in the bacteria strain TOP10 (Thermofisher Scientific, Illkirch Graffenstaden, France).

#### Lentivector production

Lentivector particles were produced using the CaCl_2_ method based by tri-transfection with the plasmids pCMV-dR8.91 and pCMV-VSVG, CaCl_2_ and Hepes Buffered Saline (Sigma-Aldrich, Saint-Quentin-Fallavier, France), into HEK-293FT cells. Viral supernatants were harvested 48 hours after transfection, passed through 0.45 μm PVDF filters (Dominique Dutscher SAS, Brumath, France), and stored in aliquots at -80°C until use. Viral production titers were assessed on HT1080 cells with serial dilutions of a lentivector expressing GFP and scored for green fluorescent protein (GFP) expression by flow cytometry analysis on a BD FACSVerse (BD Biosciences, Le Pont de Claix, France).

#### Reporter activity assay

For reporter lentivectors, luciferase activities were performed *in vitro* and *in vivo* were performed using Dual-Luciferase Reporter Assay (Promega, Charbonnières-Les-Bains, France). Briefly, proteins from HL-1 cells were extracted with Passive Lysis Buffer (Promega France). Bioluminescence was quantified with a luminometer (Centro LB960, Berthold, Thoiry, France) from 9 to 12 biological replicates and with 3 technical replicates for each sample in the analysis plate.

### FISH

HL-1 cells were cultured in 12 well plates on fibronectin-gelatin coated 15 mm coverglass 1.5 thickness (Menzel-Gläser). FISH probes were produced and purchased from Sigma-Aldrich, and delivered HPLC purified at 50 nmol. The 3/2 probes used per target (NEAT1 and NEAT1_2 isoform respectively) are between 38-40 mer long and are conjugated to one Cy3 through 5’ amino acid modifications (see Supplementary file 2 for sequences).

FISH was performed as previously described (http://www.singerlab.org/protocols). Briefly, cells were fixed with 4% paraformaldehyde (electron microscopy science), rinsed twice, and permeabilized overnight in 70% ETOH. Then cells were pre-hybridized in a 15% formamide/2X SSC buffer at room temperature. The hybridization reaction was performed overnight at 37°C with a Mix of 2XSSC, 0.5 mg/mL yeast tRNA, 15% formamide, 10% dextran sulfate, and 10 ng of mixed probes. Then the coverslip was rinsed two time 10min in 2X SSC and 1XSSC for 10 min, before mounting on Moviol mounting medium supplemented with DAPI. Three-dimensional image stacks were captured on LSM780 Zeiss confocal microscope, camera lens x63 with Z acquisition of 0.45 μM, and Zen software (Zeiss).

#### SmiFISH

A set of target-specific primary probes was produced and purchased from Integrated DNA Technologies (IDT). Each probe carried an additional 28 nt-long sequence “FLAP” which is not represented in either mouse or human genomes. The primary probes against bicistronic mRNA were complementary to the fluorescent secondary probe FLAP-X, and the primary probes Neat1 were complementary to the fluorescent secondary probe FLAP-Y. The two secondary probes FLAP-X and FLAP-Y were also from IDT, conjugated to fluorophores Cy3 and Cy5, respectively. All probe sequences are presented in Supplementary file 2. SmiFISH was performed as previously described (Tsanov et al., 2016). Cells were grown to 80% confluence in 6-well plates and subjected or not to 4 hr of hypoxia,. Cells were fixed with 4% paraformaldehyde for 20 min (electron microscopy science) at room temperature, rinsed twice and permeabilized overnight in 70% ETOH. Cy3 and Cy5 labeled fluorescent FLAPs were preannealed to primary probes prior to in situ hybridization. Then cells were pre-hybridized with probes (40 pmol) in a 15% formamide/1X SSC buffer at 37°C. The hybridization reaction was performed overnight at 37°C with Mix 1 (2XSSC, 25 μg/μL yeast tRNA, 100% formamide, FLAP-structured duplex (FLAP-Y duplex + FLAP-X duplex) and H2O) + Mix 2 (20 mg/mL RNAse free BSA, 200mM VRC, 40% dextran sulfate and H2O). Then the coverslip was rinsed five times in 1X SSC 15% formamide mix at 37°C and twice in PBS before mounting on Dako mounting medium supplemented with DAPI. Three-dimensional image stacks were captured on Zeiss Axiomager Z3 Apotome confocal microscope, camera lens x63 with Z acquisition of 0.45 μM, and Zen software (Zeiss). For Figure 2, images were analyzed with a script for ImageJ. For each segmented nucleus, spots were segmented by detecting local maxima after applying a laplacien filter. For Figure 3, images were analyzed with IMARIS. For each image, spots were detected using the “spot” function and the colocalization with the “co-localize spots” function.

#### Western blot

Cells were harvested on ice, washed with cold PBS, and collected on RIPA buffer (Biobasic supplemented with protease inhibitor (Sigma). Protein concentration was measured using BCA Protein Assay Kit (Interchim), and equal amounts of proteins were subjected to SDS-PAGE (TGX Stin Free FastCast Acrylamid, 12%, Bio-Rad, 161-0185) and transferred onto nitrocellulose membrane (Transblot Turbo, Bio-Rad, 1704271). Membranes were washed in Tris-buffered saline supplemented with 0.05% Tween-20 and then saturated in Tris-buffered saline supplemented with 0.05% Tween-20 with 5% BSA, incubated overnight with primary antibodies in Tris-buffered saline supplemented with 0.05% Tween-20 with 5% BSA, washed and revealed with Clarity Western ECL Substrate (Bio-Rad, 170-5060). Western blotting was conducted using standard methods with the following antibodies: Rabbit anti-PSPC1 (bethyl laboratory, A303-205A), Rabbit anti-P54nrb (Santacruz, sc67016), Rabbit Histone H3 (Cell Signaling, 4499), Mouse GAPDH (SantaCruz, SC32233).

#### Capillary Western

Diluted protein lysate was mixed with fluorescent master mix and heated at 95°C for 5 minutes. 3 μL of protein mix (1mg/mL maximal concentration) containing Protein Normalization Reagent, blocking reagent, wash buffer, target primary antibody (rabbit anti-P54nrb, diluted 1:200 [Santacruz, sc67016], rabbit anti-PSPC1 diluted 1:100 [bethyl laboratory, A303-205A], mouse anti-SFPQ diluted 1:100 [Abcam, Ab11825]; rabbit anti-FGF1 diluted 1:25 [Abcam Ab207321], rabbit anti-Nucleolin diluted 1:50 [Novus biological, NB600-241], secondary-HRP (ready to use rabbit of mouse “detection module”, DM-001 or DM-002), and chemiluminescent substrate were dispensed into designated wells in a manufacturer-provided microplate. The plate was loaded into the instrument (Jess, Protein Simple) and proteins were drawn into individual capillaries on a 25 capillary cassette (12-230 kDa) (SM-SW004). Normalization reagent allow detecting total protein in the capillary through the binding of amine group by a biomolecule and to get rid of housekeeping protein that can arbor an inconsistent and unreliable expression. Graph plotted in Figures represent chemiluminescence value before normalization.

#### Measurement of protein half-life

To measure protein half-life, HL-1 cardiomyocytes were treated with cycloheximide (InSolution CalBioChem) diluted in PBS to a final concentration of 10 μg/mL in 6 well plates. Time-course points were taken by stopping cell cultures after 0 hr, 30 min, 1 hr, 2 hr, 4 hr, 8 hr, 16 hr, or 24 hr of incubation and subsequent capillary Western analysis of cell extracts.

#### RNA purification and cDNA synthesis

Total RNA extraction from HL-1 cells was performed using TRI Reagent according to the manufacturer’s instructions (Molecular Research Center Inc, USA). RNA quality and quantification were assessed by a Nanodrop spectrophotometer (Nanodrop 2000, Thermo Scientific). 750 ng RNA was used to synthesize cDNA using a High-Capacity cDNA Reverse Transcription Kit (Applied Biosystems, France). Appropriate no-reverse transcription and no-template controls were included in the qPCR assay plate to monitor potential reagent or genomic DNA contaminations, respectively. The resulting cDNA was diluted 10 times in nuclease-free water. All reactions for the PCR array were run in biological triplicates.

#### qPCR

7.5 ng cDNA were mixed with 2X TB green Premix Ex Taq II (Takara, RR820B), 10μM forward and reverse primers, according to manufacturer instruction. qPCR reactions were performed on Viia7 (AppliedBiosystems) and the oligonucleotide primers used are detailed in Supplementary file 2.

#### ddPCR

ddPCR reaction for NEAT1 knockdown control were performed with the Bio-Rad system. The ddPCR reaction mixture (22 μl) contained 2x QX200™ ddPCR™ EvaGreen Supermix (no dUTP) (Bio-Rad), 2 μM of a mix of forward and reverse primers (Supplementary file 2), and 2/4/6μL of cDNA depending on the target. The reaction mixture was transferred for droplet generation by AutoDG System (Bio-Rad) in individual wells of disposable DG32 Automated Droplet Generator Cartridges that were already placed in the cartridge holder. The droplet was generated by AutoDG System, between 15000-20000 droplets/well. The prepared droplet emulsions were further loaded in ddPCR 96-Well Plates (Bio-rad) by aspirating 40 μl from the DG32 cartridge by the AutoDG System. The plate was then heat sealed with pierceable foil using a PX1 PCR plate sealer 5s at 180°C (Bio-Rad), and PCR amplification was carried out in a T100 thermal cycler (Bio-Rad). The thermal consisted of initial denaturation at 95°C for 5 min followed by 40 cycles of 95°C for 30 s (denaturation) and 60°C for 1 minute (annealing/elongation) with a ramp of 2°C/s, a signal stabilization step at 4°C 5min followed by 90°C 5min. After PCR amplification the positive droplets were counted with a QX200 droplet reader (Bio-Rad).

#### Cell fractionation

HL-1 cells placed in normoxia or hypoxia and transduced by P54nrb-HA construct were trypsinized, rinsed with PBS and lysed in solution 1 (Hepes 50 mM/NaCl 150 mM pH7.3, digitonin (100 μg/mL), EDTA 1 mM, protease inhibitor cocktail) and incubated on ice. Then the lysate was centrifugated at 2000g for 5 min and the supernatant (cytosolic fraction) was aliquoted. Then the pellet was rinsed in PBS, and incubated in solution 2 (Hepes 50 mM/NaCl 150 mM pH7.3, NP40 1%, EDTA 1 mM, protease inhibitor cocktail) during 30 min at 4°C. After centrifugation at 7000g, the pellet was rinsed and resuspended in solution 3 (Tris/HCl 50 mM, NaCl 150 mM, NP40 1%, sodium deoxycholate 0.5%, SDS 0.1% (RIPA), protease inhibitor cocktail) and incubated for 10 min at 4°C. Finally, the lysate was centrifuged for 10 min at 8200g and the supernatant was aliquoted (nuclear fraction).

#### Immunoprecipitation

Immunoprecipitation experiments were realized with 150 μg of total protein amounts from the cytosolic and nuclear fraction in normoxia or hypoxia, with a HA antibody (H9558/H3663, Sigma) using EZ view red protein G beads (Sigma). The beads-antibody-protein mix was incubated overnight at 4°C and bounds protein were eluted with 35 μg HA peptides diluted in PBS (Sigma); then Laemmli buffer was added and the eluate heated at 95°C 2 min.

#### In-gel trypsin digestion and mass spectrometry analysis

For mass spectrometry analysis, immunoprecipitated samples, prepared in triple or quadruple biological replicates for each condition, were submitted to an additional protein reduction in 24.5 mM dithiothreitol for 30 min at 56°C followed by alkylation of cysteine residues in 74 mM iodoacetamide for 30 min in the dark at room temperature. Each reduced/alkylated sample was loaded onto 1D SDS-PAGE gel (stacking 4% and separating 12% acrylamide). For a one-shot analysis of the entire mixture, no fractionation was performed, and the electrophoretic migration was stopped as soon as the protein sample entered the separating gel. The gel was briefly stained using Quick Coomassie Blue (Generon). Each single slice containing the whole sample was excised and subjected to in-gel tryptic digestion using modified porcine trypsin (Promega, France) at 10 ng/μl as previously described (Shevchenko A et al., 2001). The dried peptide extracts obtained were dissolved in 17 μl of 0.05% trifluoroacetic acid in 2% acetonitrile and analyzed by online nanoLC using an Ultimate 3000 RSLCnano LC system (Thermo Scientific Dionex) coupled to an LTQ Orbitrap Velos mass spectrometer (Thermo Scientific, Bremen, Germany) for data-dependent CID fragmentation experiments. 5μl of each peptide extracts were loaded in two or three injection replicates onto 300μm ID x 5mm PepMap C18 pre-column (ThermoFisher, Dionex) at 20 μl/min in 2% acetonitrile, 0.05% trifluoroacetic acid. After 5 minutes of desalting, peptides were online separated on a 75 μm ID x 50 cm C18 column (in-house packed with Reprosil C18-AQ Pur 3 μm resin, Dr. Maisch ; Proxeon Biosystems, Odense, Denmark), equilibrated in 95% of buffer A (0.2% formic acid), with a gradient of 5 to 25% of buffer B (80% acetonitrile, 0.2% formic acid) for 80 min then 25% to 50% for 30 min at a flow rate of 300 nL/min. The LTQ Orbitrap Velos was operated in data-dependent acquisition mode with the XCalibur software (version 2.0 SR2, Thermo Fisher Scientific). The survey scan MS was performed in the Orbitrap on the 350–1,800 m/z mass range with the resolution set to a value of 60,000. The 20 most intense ions per survey scan were selected with an isolation width of 2 m/z for subsequent data-dependent CID fragmentation, and the resulting fragments were analyzed in the linear trap (LTQ). The normalized collision energy was set to 30%. To prevent repetitive selection of the same peptide, the dynamic exclusion duration was set to 60 s with a 10 ppm tolerance around the selected precursor and its isotopes. Monoisotopic precursor selection was turned on. For internal calibration the ion at 445.120025 m/z was used as lock mass.

#### MS-based protein identification

Acquired MS and MS/MS data as raw MS files were converted to the mzDB format using the pwiz-mzdb converter (version 0.9.10, https://github.com/mzdb/pwiz-mzdb) executed with its default parameters (Bouyssié et al., 2015). Generated mzDB files were processed with the mzdb-access library (version 0.7, https://github.com/mzdb/mzdb-access) to generate peak lists. Peak lists were searched against the UniProtKB/Swiss-Prot protein database with mus musculus taxonomy (16,979 sequences) in the Mascot search engine (version 2.6.2, Matrix Science, London, UK). Cysteine carbamidomethylation was set as a fixed modification and methionine oxidation as a variable modification. Up to two missed trypsin/P cleavages were allowed. Mass tolerances in MS and MS/MS were set to 10 ppm and 0.6 Da, respectively. Validation of identifications was performed through a false-discovery rate set to 1% at protein and peptide-sequence match level, determined by target-decoy search using the in-house-developed software Proline software version 1.6 (Bouyssié et al., 2020).

#### Polysomal RNA preparation

HL-1 cells were cultured in 150-mm dishes. 10 min before harvesting, cells were treated with cycloheximide at 100 mg/mL. Cells were washed with PBS at room temperature containing 100 mg/mL cycloheximide and harvested with Trypsin. After centrifugation at 500g for 3 min at 4°C, cells were washed two times in PBS cold containing 100 mg/mL cycloheximide, and cells were lysed by hypotonic lysis buffer (10 mM HEPES-KOH Ph7.5; 10 mM KCl; 1.5 mM MgCl_2_) containing 100 mg/mL cycloheximide. Cells were centrifuged at 500g for 3 min and lysed by lysis solution containing hypotonic buffer, 1 mM DTT, 0.5U/mL Rnasin, and protease inhibitor 100X. Cells were centrifuged by two times, first at 1000g for 10 min at 4°C and second at 10 000g for 15 min; the supernatants were collected and loaded onto a 10-50% sucrose gradient. The gradients were centrifuged in a Beckman SW41 Ti rotor at 39,000 rpm for 2.3 hr at 4°C without a brake. Fractions were collected using a Foxy JR ISCO collector and UV optical unit type 11. RNA was purified from pooled heavy fractions containing polysomes (fractions 12-19) as well as from cell lysate before gradient loading.

#### qPCR fluidigm array

The DELTAgene Assay™ was designed by Fluidigm Corporation (San Francisco, USA). The qPCR-array was performed on BioMark with the Fluidigm 96.96 Dynamic Array following the manufacturer’s protocol (Real-Time PCR Analysis User Guide PN 68000088). The list of primers is provided in Supplementary file 2. A total of 25 ng of cDNA was preamplified using PreAmp Master Mix (Fluidigm,100-5581, San Francisco, USA) in the plate thermal cycler at 95°C for 10 min, 16 cycles at 95°C for 15 sec, and 60°C for 4 min. The preamplified cDNA was treated with Exonuclease I in the plate thermal cycler at 37°C for 30 min, 80°C for 15 min and 10°C infinity. The preamplified cDNA was mixed with 2x TaqMan Gene Expression Master Mix (Applied Biosystems), 20 μM of mixed forward and reverse primers, and sample Loading Reagent (Fluidigm, San Francisco, USA). The sample was loaded into the Dynamic Array gene expression 96.96 IFC (Fluidigm San Francisco, USA). The qPCR reactions were performed in the BioMark RT-qPCR system. Data were analyzed using the BioMark RT-qPCR Analysis Software Version 4.5.2.

GAPDH rRNA was used as a reference gene, and all data were normalized based on GAPDH rRNA level. Relative quantification (RQ) of gene expression was calculated using the 2^-ΔΔCT^ method. When the RQ value was inferior to 1, the fold change was expressed as -1/RQ. The oligonucleotide primers used are detailed in Supplementary file 2.

## QUANTIFICATION AND STATISTICAL ANALYSIS

### qPCR and ddPCR analysis

qPCR data were analyzed on Quantstudio (AppliedBiosystems). RPL11 or HPRT were used as reference gene. Relative quantification (RQ) of gene expression was calculated using the 2-ΔΔCT method. ddPCR data was analyzed using the QuantaSoft 1.7.4 software (Bio-Rad). HPRT was used as a reference gene, and Neat1 RNA expression was normalized by normoxia control and expressed in %.

### Label-free quantitative proteomics analysis

For label-free relative quantification across samples, raw MS signal extraction of identified peptides was performed using Proline. The cross-assignment of MS/MS information between runs was enabled (it allows to assign peptide sequences to detected but non-identified features). Each protein intensity was based on the sum of unique peptide intensities and was normalized across all samples by the median intensity. Missing values were independently replaced for each run by its 5% quantile. After log2-transformation of the data, the values of the technical replicates were averaged for each analyzed samples. For each pairwise comparison, an unpaired two-tailed Student’s t-test was performed. Proteins were considered significantly enriched when their absolute log2-transformed fold change was higher than 1 and their p-value lower than 0.05. To eliminate false-positive hits from quantitation of low-intensity signals, two additional criteria were applied: only the proteins identified with a total number of averaged peptide spectrum match (PSM) counts>4 and quantified in a minimum of two biological replicates, before missing value replacement, for at least one of the two compared conditions were selected. Volcano plots were drawn to visualize significant protein abundance variations between the two compared conditions. They represent -log10 (p-value) according to the log2 ratio. The complete list of proteins identified and quantified in immunopurified samples and analyzed according to this statistical procedure is described in Supplementary file 7.

### Dual luciferase system

Data were analyzed on MicroWin 2000. Background noise was measured with non-transduced cell samples and removed from transduced cell sample measurement. Then LucF/LucR ratio was calculated on Excel 2007 (Microsoft Office) and mean and SD were calculated as well.

### FISH

Images were analyzed with a script for ImageJ. For each segmented nucleus, spots are segmented by detecting local maxima after applying a laplacien filter. Spot colocalization is determined by the distance between them.

### Capillary Western

Data were analyzed on compass software provided by the manufacturer.

### Statistical analysis

All statistical analyses were performed using One Way ANOVA, unpaired two-tailed student t-test, or Mann-Whitney rank comparisons test calculated on GraphPad Prism software depending on n number obtained and experiment configuration. Results are expressed as mean + standard deviation, *p<0.05, **p<0.01,***<0.001, ****<0.0001.

## DATA AND CODE AVAILABILITY

The MS proteomics data have been deposited to the ProteomeXchange Consortium via the PRIDE partner repository with the dataset identifier PXD024067.

Reviewer account details: **Username:** **Password:** vlzsdovC Lentivector plasmid sequences are available on Dryad doi:10.5061/dryad.nvx0k6dq7

All other data are presented in supplementary files.

## SUPPLEMENTAL INFORMATION

Supplemental information can be found online at https://doi.org

## ACKNOWLEDGMENTS

Our thanks go to J.J. Maoret and F. Martins from the Inserm UMR1297 I2MC GeT-TQ plateau of the GeT platform Genotoul (Toulouse), A. Lucas from the I2MC We-Met Functional Biochemistry Facility (Toulouse) and Rémy Flores-Flores from the I2MC imaging plateau. We also thank L. Colras for technical assistance.

This work was supported by Région Occitanie (Midi-Pyrénées), Association pour la Recherche sur le Cancer (ARC), Fondation Toulouse Cancer Santé and Agence Nationale de la Recherche ANR-18-CE11-0020-RIBOCARD, European funds (Fonds Européens de Développement Régional, FEDER), Toulouse Métropole, and by the French Ministry of Research with the Investissement d’Avenir Infrastructures Nationales en Biologie et Santé program (ProFI, Proteomics French Infrastructure project, ANR-10-INBS-08). A.C.G, F.H., and E.R had fellowships from the Ligue Nationale Contre le Cancer (LNCC).

## AUTHOR CONTRIBUTIONS

A.C.G., A.C.P., E.R. and F.D. conceived, and A.C.P. supervised the project. A.C.G., A.C.P., A.K.H., B.G.S., C.F., E.B., E.L., E.R., F.D., F.T., I.A. and P.V. designed the experiments. A.C.G., C.F., E.R., F.D., F.H., F.P. and J.A. performed the experiments. A.C.G., A.C.P., A.K.H., B.G.S. C.F., E.B., E.L., E.R., F.D., F.M., F.T., I.A., P.V. analyzed the results. A.C.G., A.C.P., E.R. and F.D. wrote the manuscript. A.C.G., A.C.P., A.K.H., B.G.S., C.F. E.L, E.R., F.D., F.H., F.M., F.T. and O.B.S. corrected the manuscript.

## DECLARATION OF INTEREST

The authors have no conflict of interest.

---

**Figure 1-figure supplement1.**
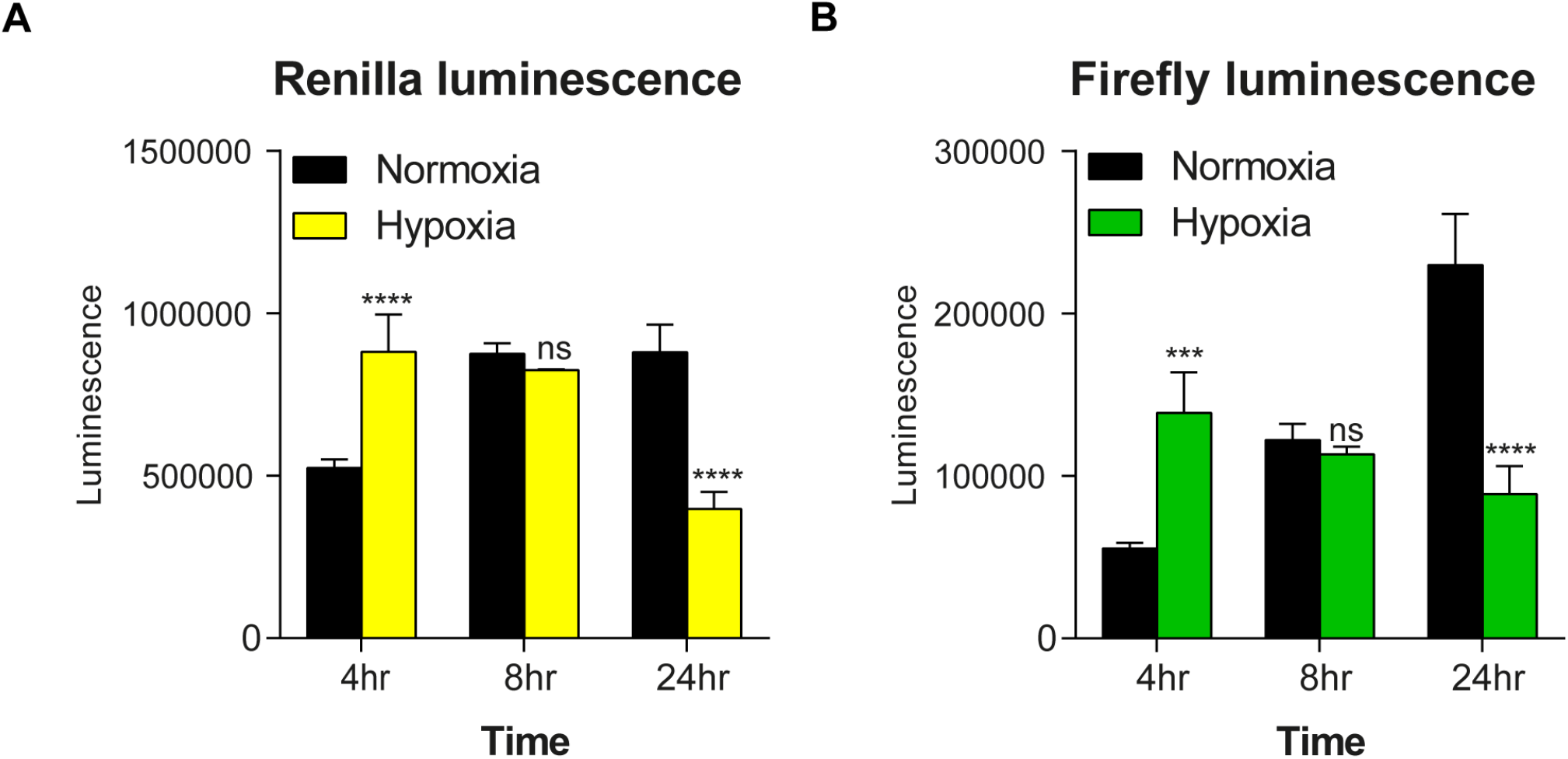
Bicistronic vector LucR and LucF expression in hypoxic HL-1 cells. Renilla (A) and firefly (B) luciferase activities were measured in HL-1 cardiomyocytes transduced by the bicistronic vector with the FGF1 IRES, after 4 hr, 8 hr or 24 hr of hypoxia or normoxia. The experiments were performed with triplicates from four distinct transduced cell samples (Supplementary file 1). The graphs show a representative triplicate experiment.

**Figure 1-figure supplement2.**
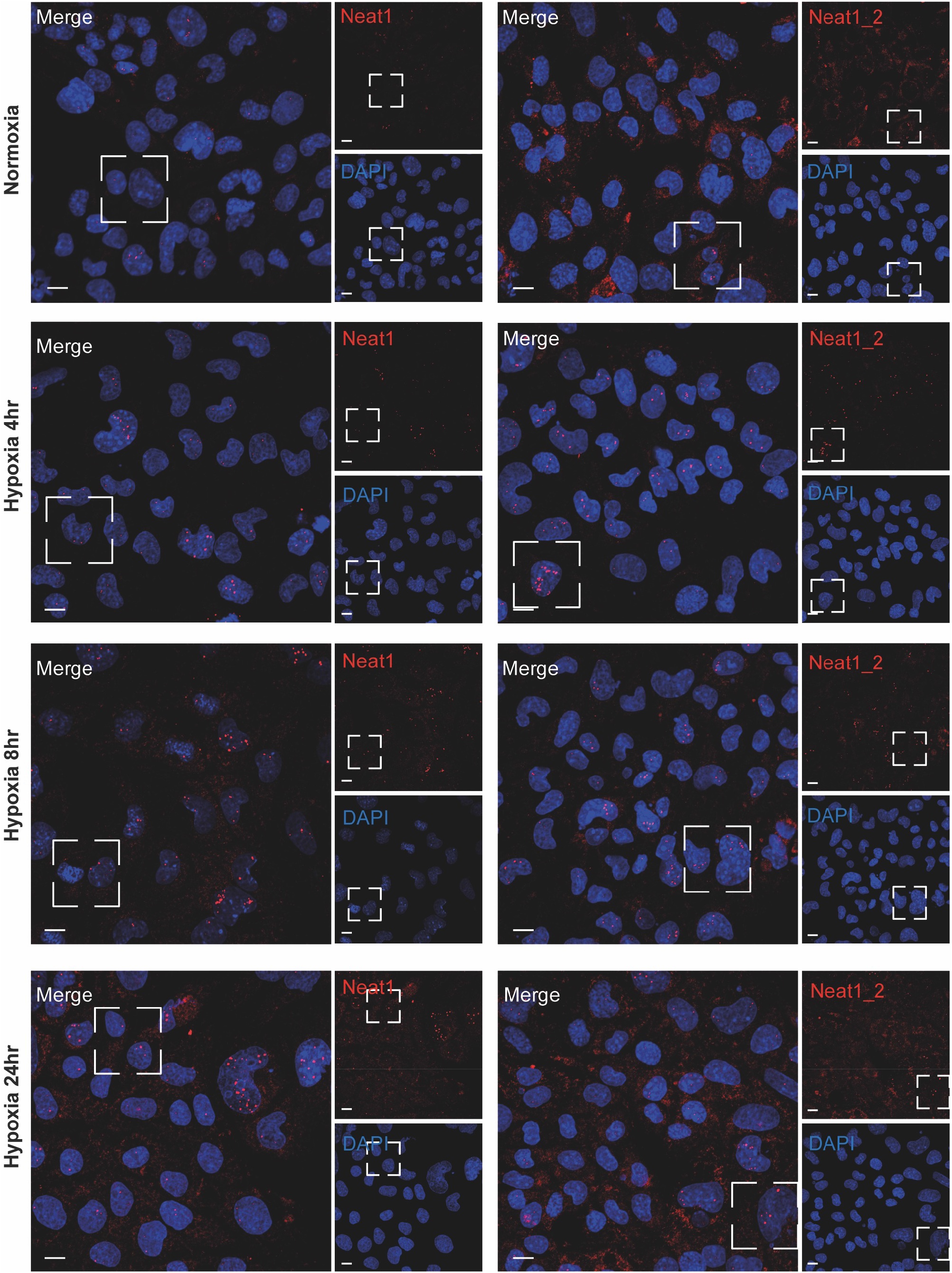
Detection of Neat1 and Neat1_2 in hypoxic HL-1 by FISH. FISH experiment with representative images of Neat1 (both isoforms)(left panels) or Neat1_2 isoform (right panels) in normoxia and hypoxia at 4 hr, 8 hr, 24 hr in HL-1 cardiomyocytes. DAPI, Neat1 cy3 or Neat1_2 cy3 labelling and merge. The open square represents magnified zones presented in Figure 1F and 1I. Scale bar: 10 μm.

**Figure 1-figure supplement3.**
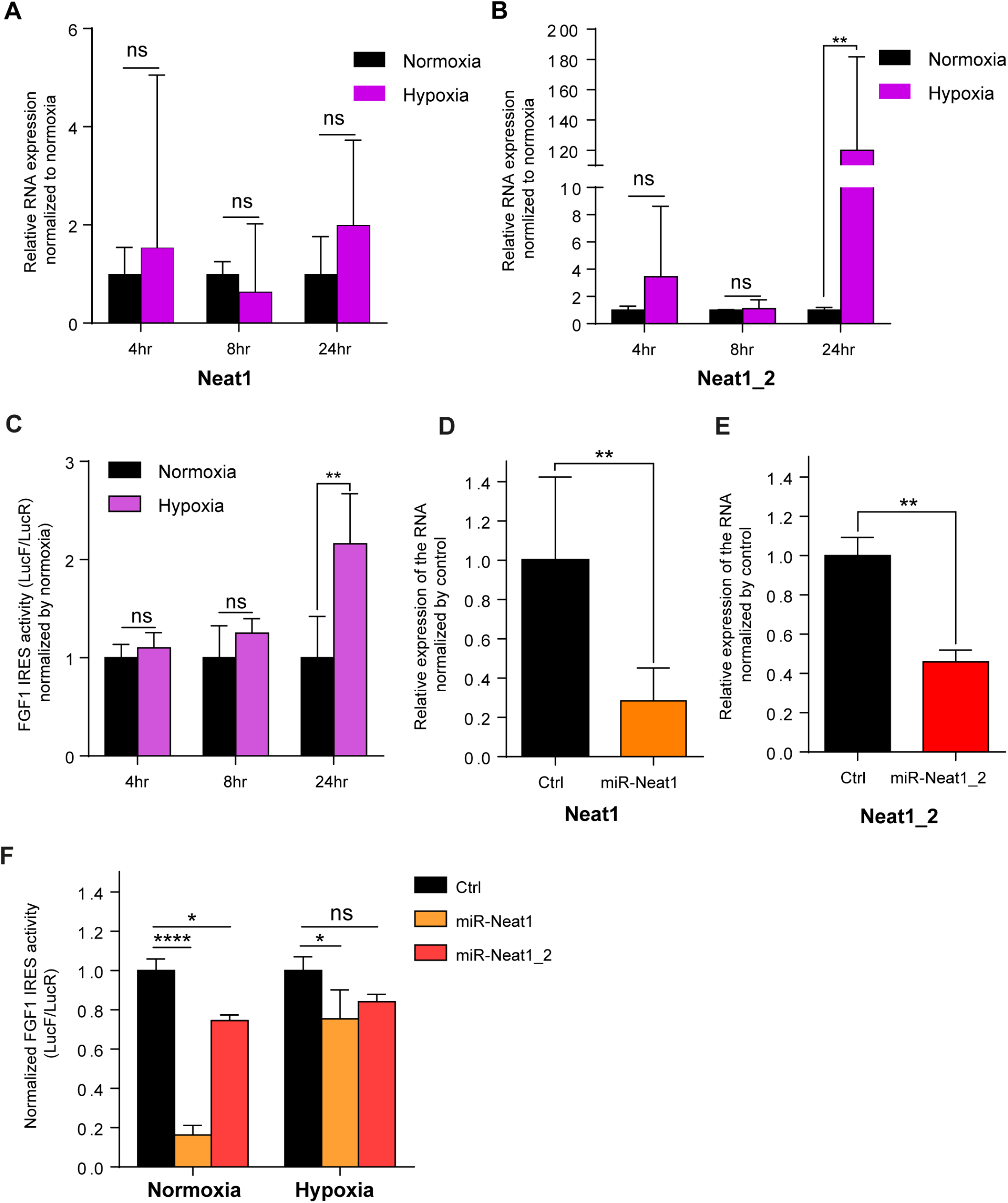
FGF1 IRES is activated by hypoxia in correlation with Neat1 induction in 67NR cells and inactivated after Neat1 knock-down. A-B) 67NR cells (mouse breast cancer) were subjected to normoxia (0 hr) or to hypoxia (1% 0_2_) during 4 hr, 8 hr and 24 hr. Neat1 (A) or Neat1_2 (B) expression was analyzed by RT qPCR (Primer sequences in Supplementary file 2). RNA expression is normalized to normoxia time point. (C) Activity of the human FGF1 IRES in 67NR cells at 4 hr, 8 hr or 24 hr of hypoxia normalized to normoxia. (D-E) Neat1 or Neat1_2 knock-down obtained by transduction of 67NR cells with lentivectors expressing an artificial miRNA targeting Neat1 (miR-Neat1, D) or Neat1_2 (miR-Neat1_2, E). To knock down Neat1_2, a pool of two lentivectors coding two different miRNAs was used (sequences in Supplementary file 2). (C) Activity of the human FGF1 IRES in 67NR cells transduced with lentivector miR-Neat1 or miR-Neat1_2 and submitted to normoxia or 24 hr of hypoxia 1% O_2_. Statistics were performed by two-way ANOVA with multiple comparison Dunnet test. Normoxia versus hypoxia for each time. *p<0.05, **p<0.005, ***p<0.0005, ****p<0.0001.

**Figure 2-figure supplement 1.**
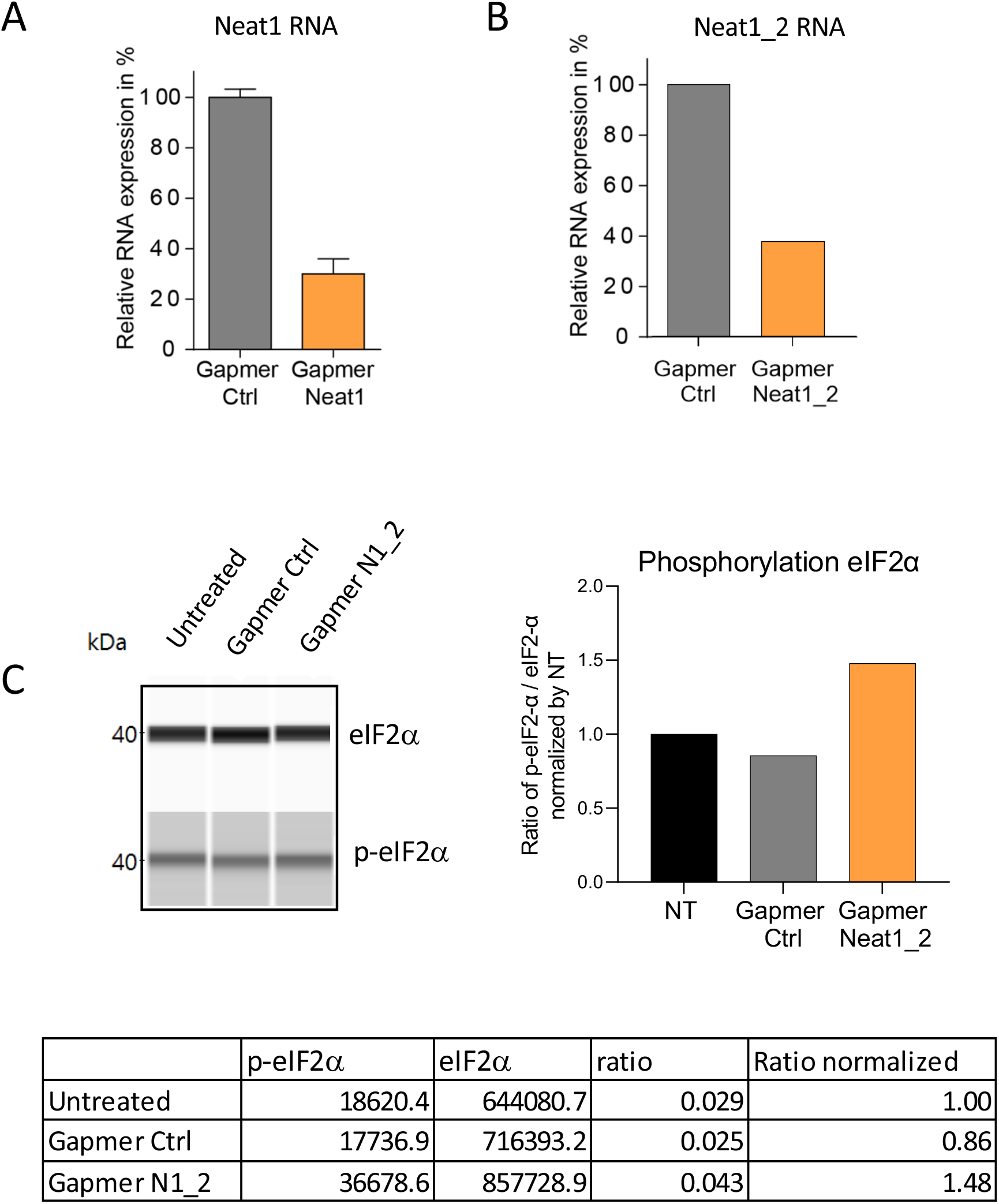
Knock-down of Neat1 and Neat1_2 in HL-1 cardiomyocytes and effect on eIF2α phosphorylation. Neat1 knock-down was performed in HL-1 cells using pooled LNA gapmers against Neat-1 (48 hr) or Neat1-2 (72 hr). Neat1 and Neat1-2 RNA expression was measured by droplet digital PCR and normalized to gapmer control (Ctrl) at 100% expression (A-B). (C) The phosphorylation of eIF2α was measured by Jess capillary Simple Western, normalized to the Jess quantification of total proteins. The ratio p-eIF2α/ eIF2α was calculated and normalized by untreated cells.

**Figure 2-figure supplement 2.**
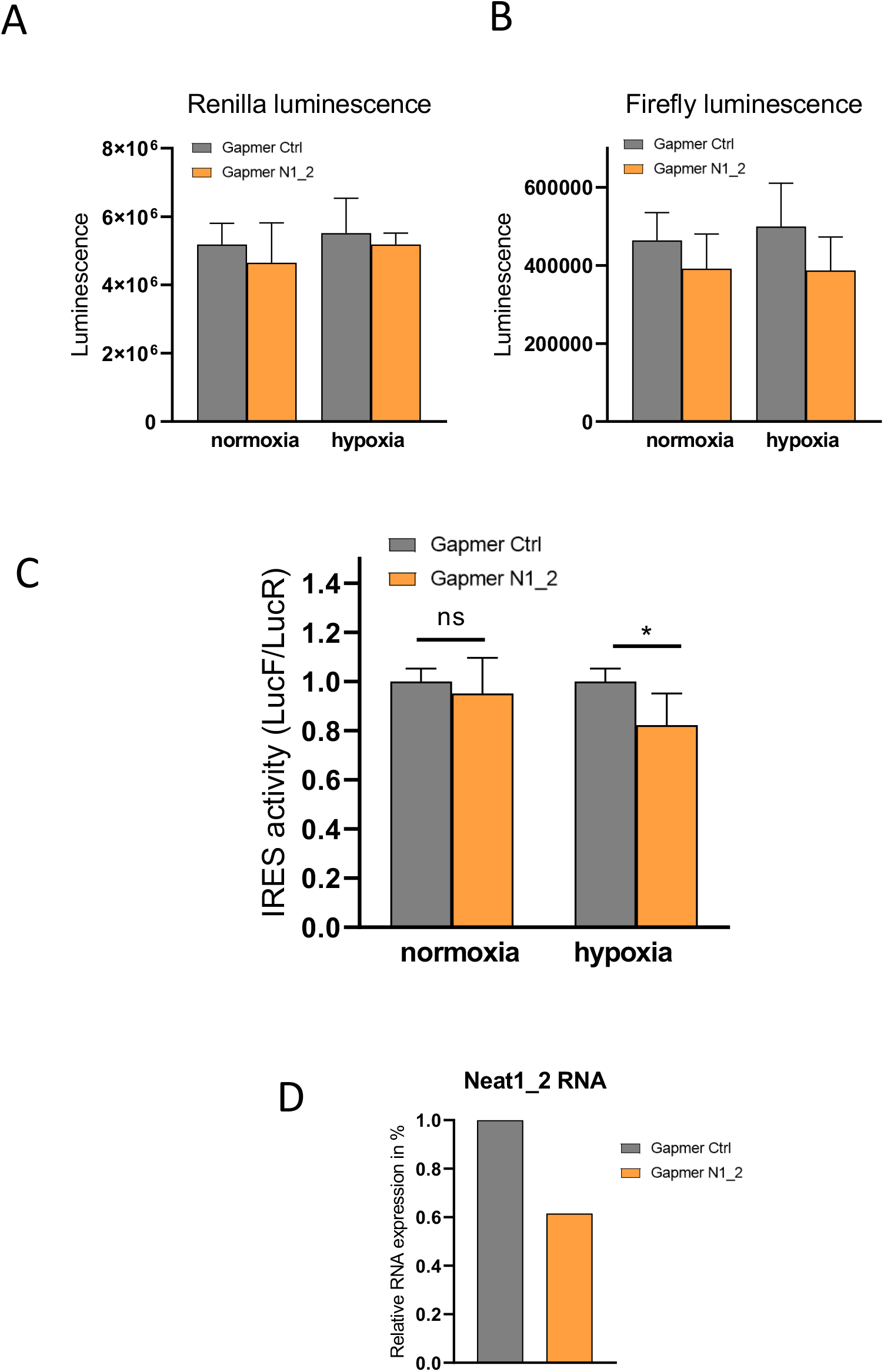
Effect of Neat 1_2 knock-down on FGF1 IRES activity. Neat1-2 knock-down was performed in HL-1 cells transduced by the bicistronic lentivector with the FGF1 IRES during normoxia or hypoxia (1% O_2_). Luciferase activities as well as the LucF/LucR ratios (defined as IRES activities) are presented. (A-B) LucR and LucF activities. (C) FGF1 IRES activities. (D) Neat1_2 RNA expression was measured by RT-qPCR and normalized to control gapmer.

**Figure 2-figure supplement 3:**
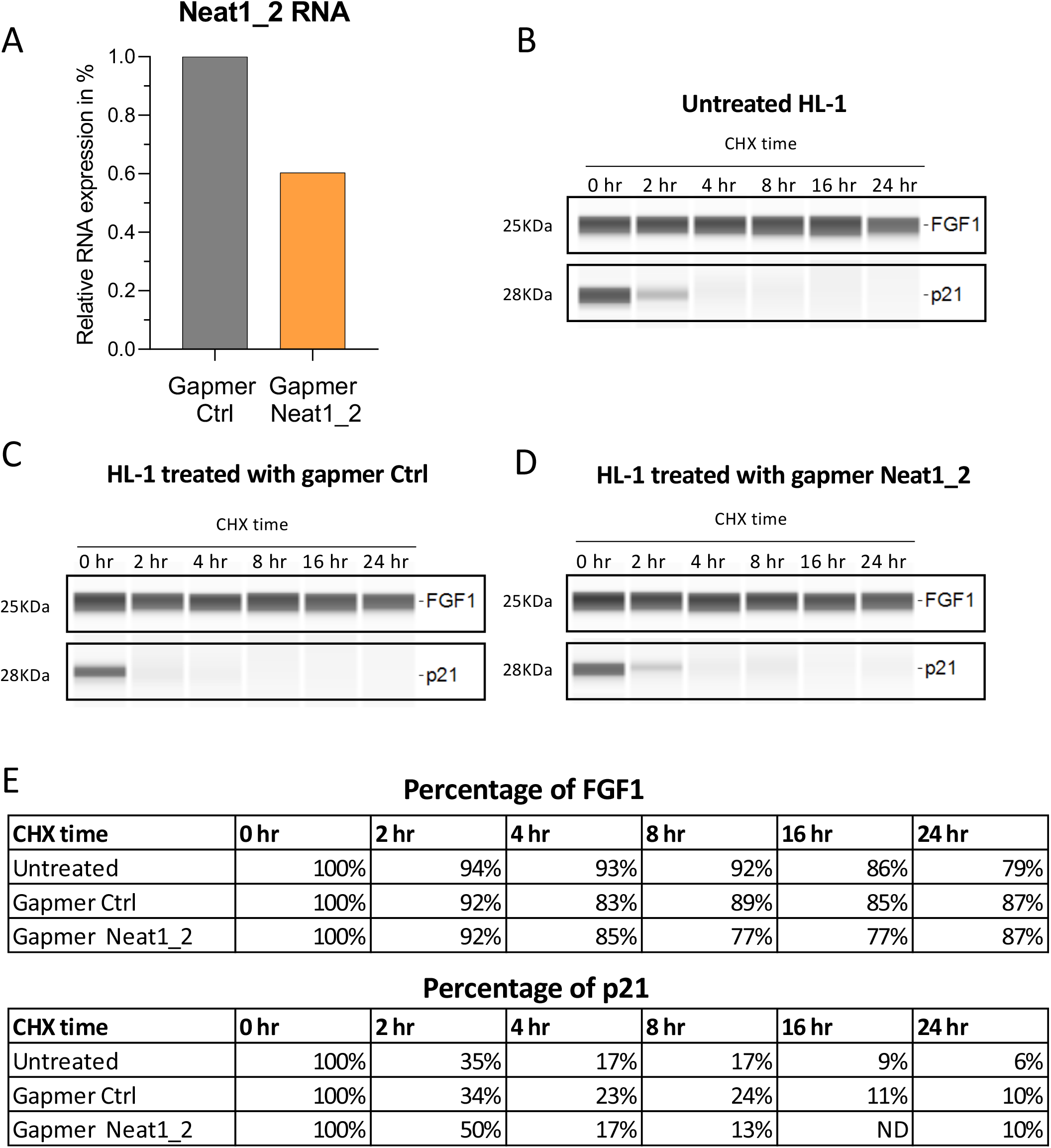
FGF1 half-life in response to Neat1_2 knock-down. (A-E) FGF1 half-life determination on HL-1 cells treated with control gapmer, Neat1_2 gapmer or untreated. The half-life determination was performed by blocking protein synthesis with cycloheximide at 10 μg/mL, with time-course points at 0 hr, 2 hr, 4 hr, 8 hr, 16 hr and 24 hr. FGF1 and p21 protein stability was measured by capillary western, with normalization to 0 hr time-course point. P21 was used as a control for its short half life. (A) Neat1_2 RNA expression was measured by RT-qPCR and normalized to control gapmer control. Capillary western showing the FGF1 and p21 proteins during the cycloheximide time course in HL-1 cells untreated (B) treated with gapmer control (C) or with gapmer Neat1_2. (E) Percentage of FGF1 and p21 normalized by 0 hr of time course point.

**Figure 4-figure supplement 1.**
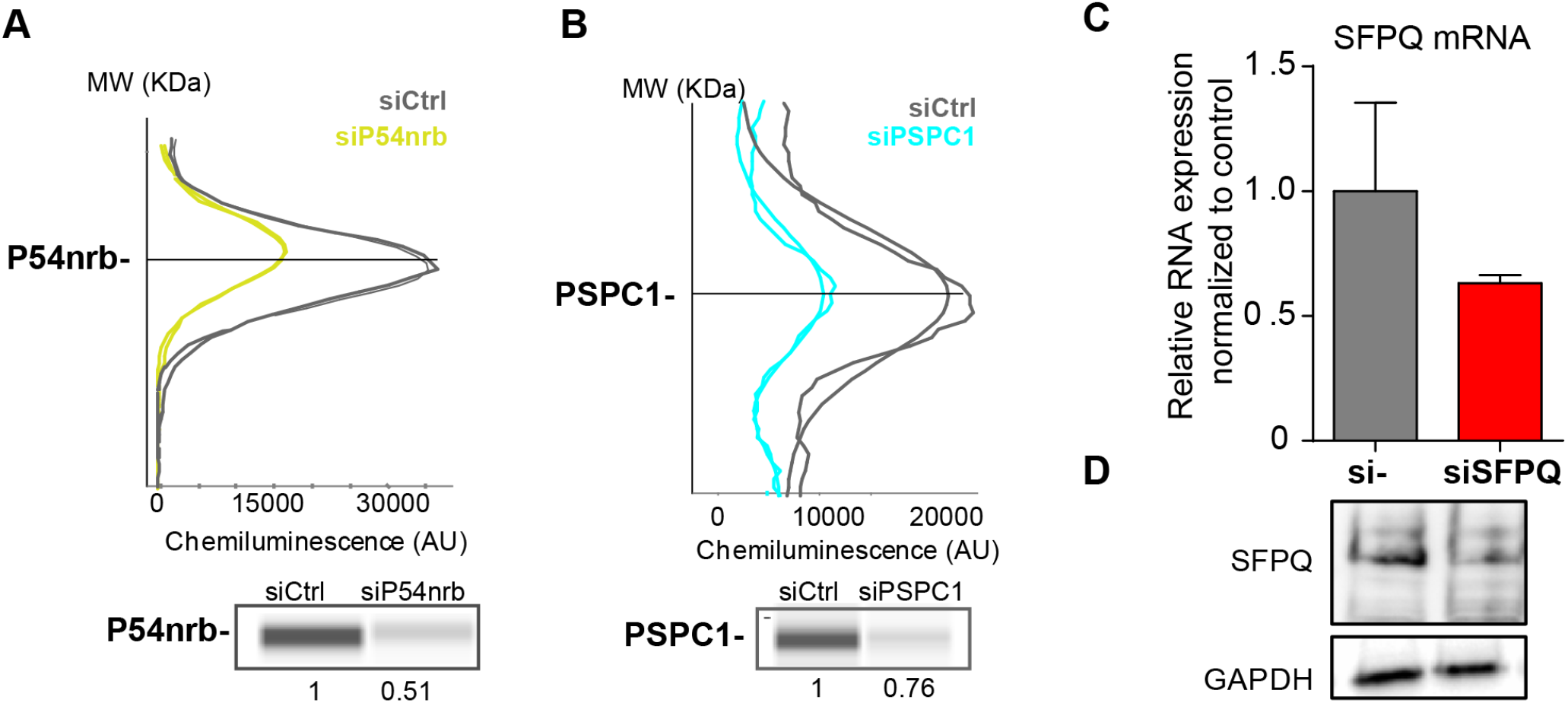
Knock-down of p54^nrb^, PCPC1 and SFPQ in HL-1 cardiomyocytes. (A-B) Capillary Simple Western detection (as described in Fig. 2) of P54^nrb^ (A) and PSPC1 (B) proteins following p54^nrb^ and PSPC1 knock-down using siRNAs. Western detection was performed 72h after siRNA treatment. (C-D) SFPQ knock-down was performed in HL-1 cells using siRNA against SFPQ. SFPQ RNA expression was measured by RT-qPCR and normalized to control siRNA (C). One representative experiment is shown with n=2 biological replicates. SFPQ protein expression was visualized by Western Blot (D). Histograms correspond to means ± standard deviation.

**Figure 4-figure supplement 2:**
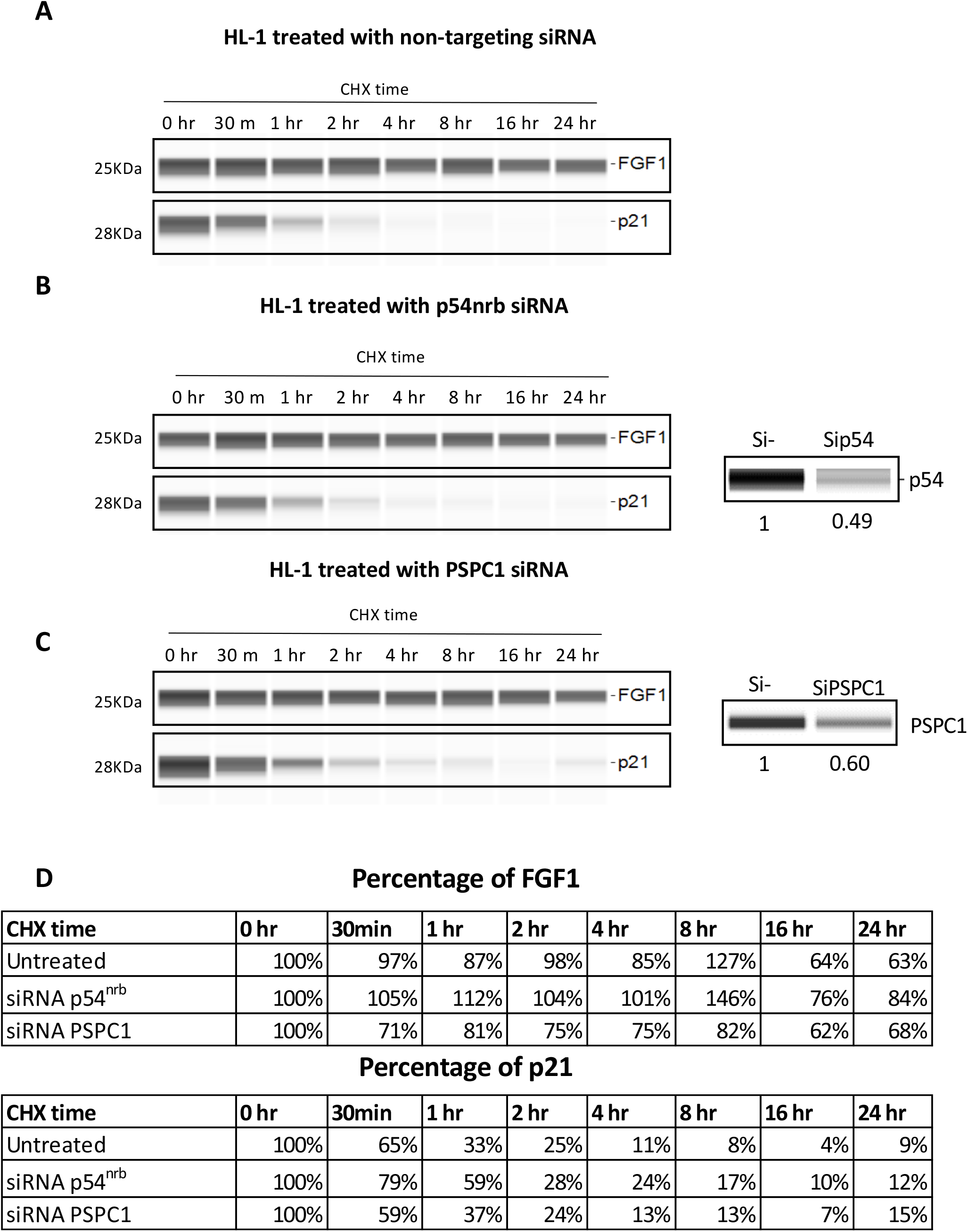
FGF1 half-life in response to p54^nrb^ or PSPC1 knock-down. (A-D) FGF1 half-life determination on HL-1 treated with p54^nrb^ siRNA, PSPC1 siRNA or control siRNA. The half life determination was performed by blocking protein synthesis with cycloheximide at 10 μg/mL, with time-course points at 0 hr, 30 min, 1 hr, 2 hr, 4 hr, 8 hr, 16 hr and 24 hr. FGF1 and p21 protein stability was measured by capillary western, with normalization to 0 hr time-course point.

**Figure 5-figure supplement 1.**
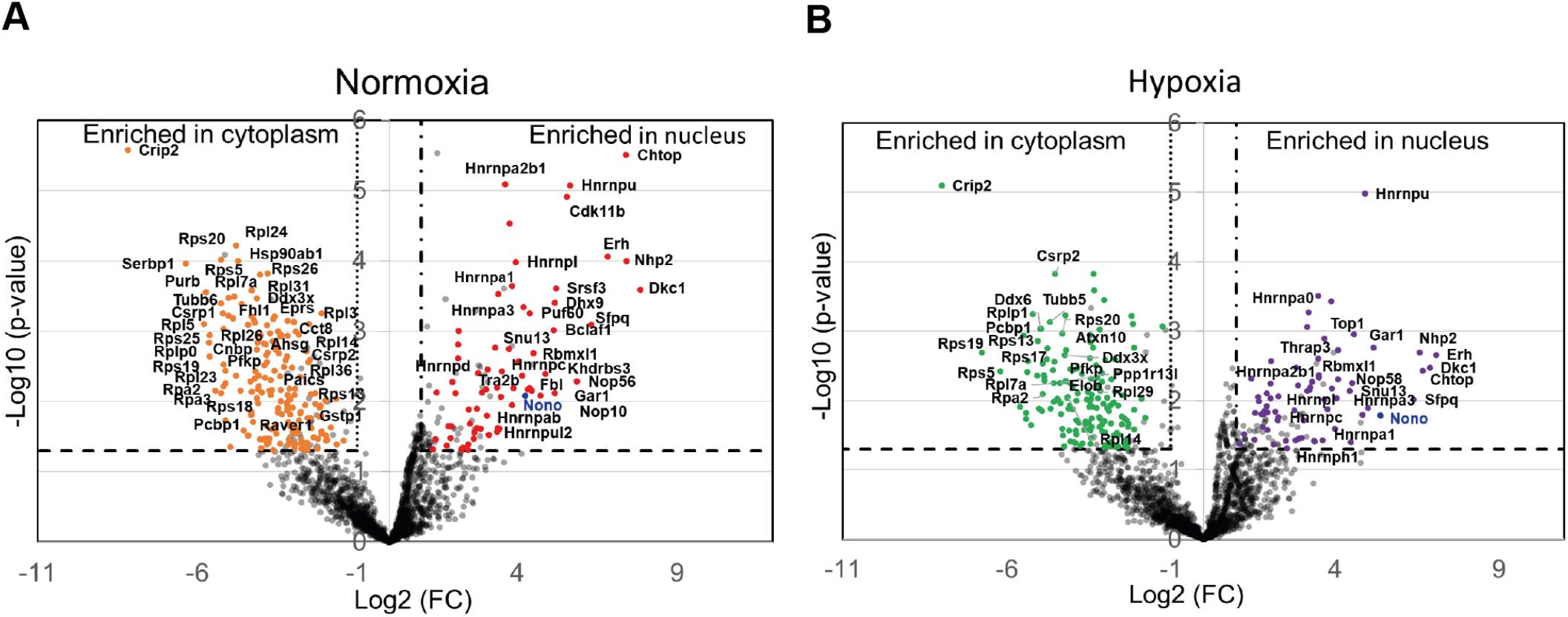
Label-free quantitative analysis of HA-P54^nrb^-bound proteins identified by mass spectrometry in different conditions. (A-B) Volcano plots showing proteins significantly enriched in the normoxia condition for nucleus (dots in red) versus cytoplasm (dots in orange) (A) or in the hypoxia condition (4 hr) for nucleus (dots in purple) versus cytoplasm (dots in green) (B). The p54^nrb^ bait (endogenous mouse Nono) is indicated in blue. An unpaired bilateral student t-test with equal variance was performed. Enrichment significance thresholds are represented by an absolute log2-transformed fold-change (FC) greater than 1 and a -log10-transformed (p-value) greater than 1.3.

**Figure 5-figure supplement 2.**
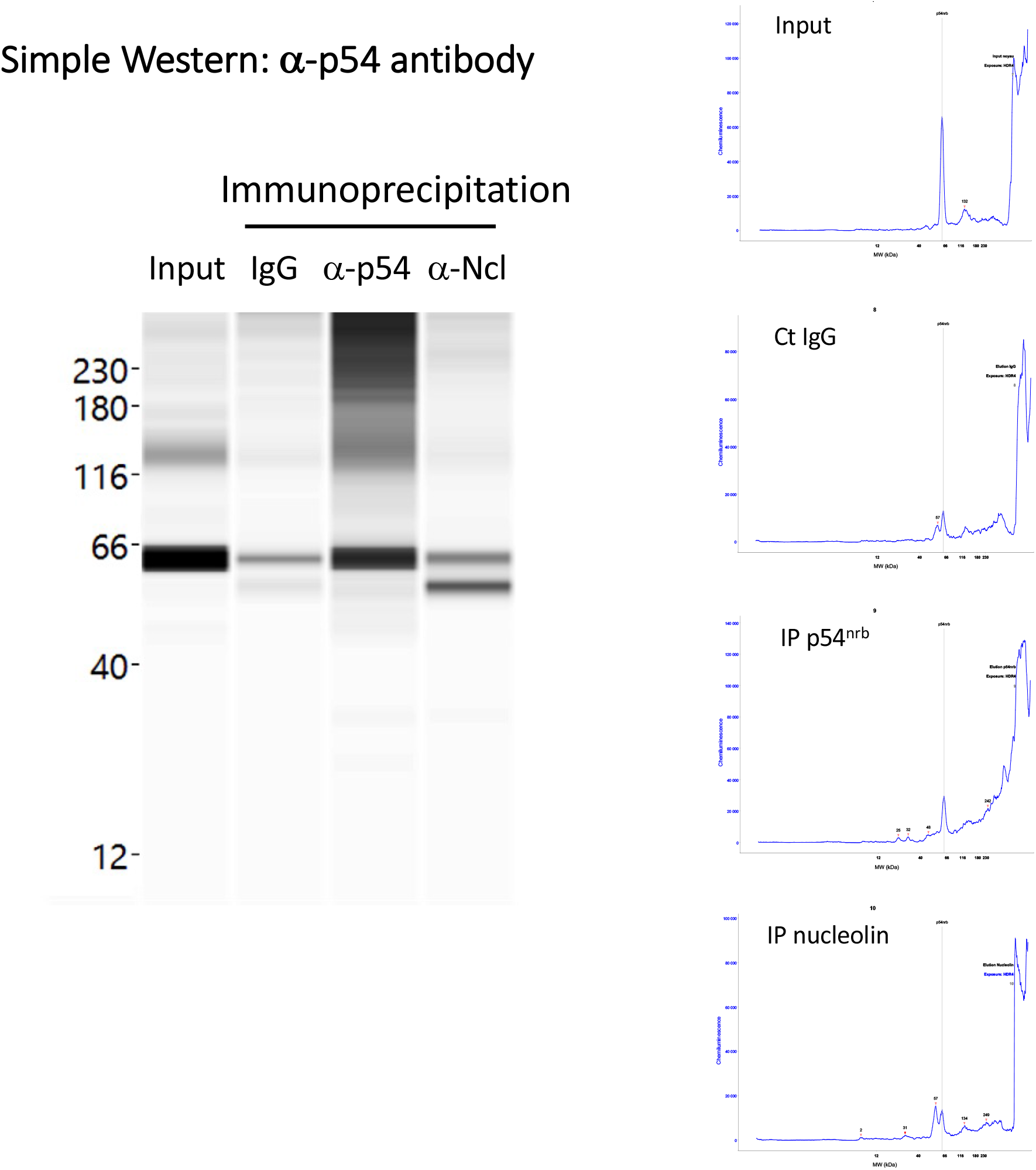
p54^nrb^ is co-immunoprecipitated by anti-nucleolin antibody. Immunoprecipitation was performed from HL-1 cell nuclear extracts, using either IgG (negative control), or antibody against p54^nrb^, or antibody against nucleolin. Capillary Simple Western (Jess) was then performed using α-p54^nrb^ antibody. Data show an enrichment of p54 using α-nucleolin antibody. Interestingly, a smaller isoform of p54 is efficiently co-immunoprecipitated, described in a previous report (Pavao et al, 2001).

**Figure 8-supplement 1.**
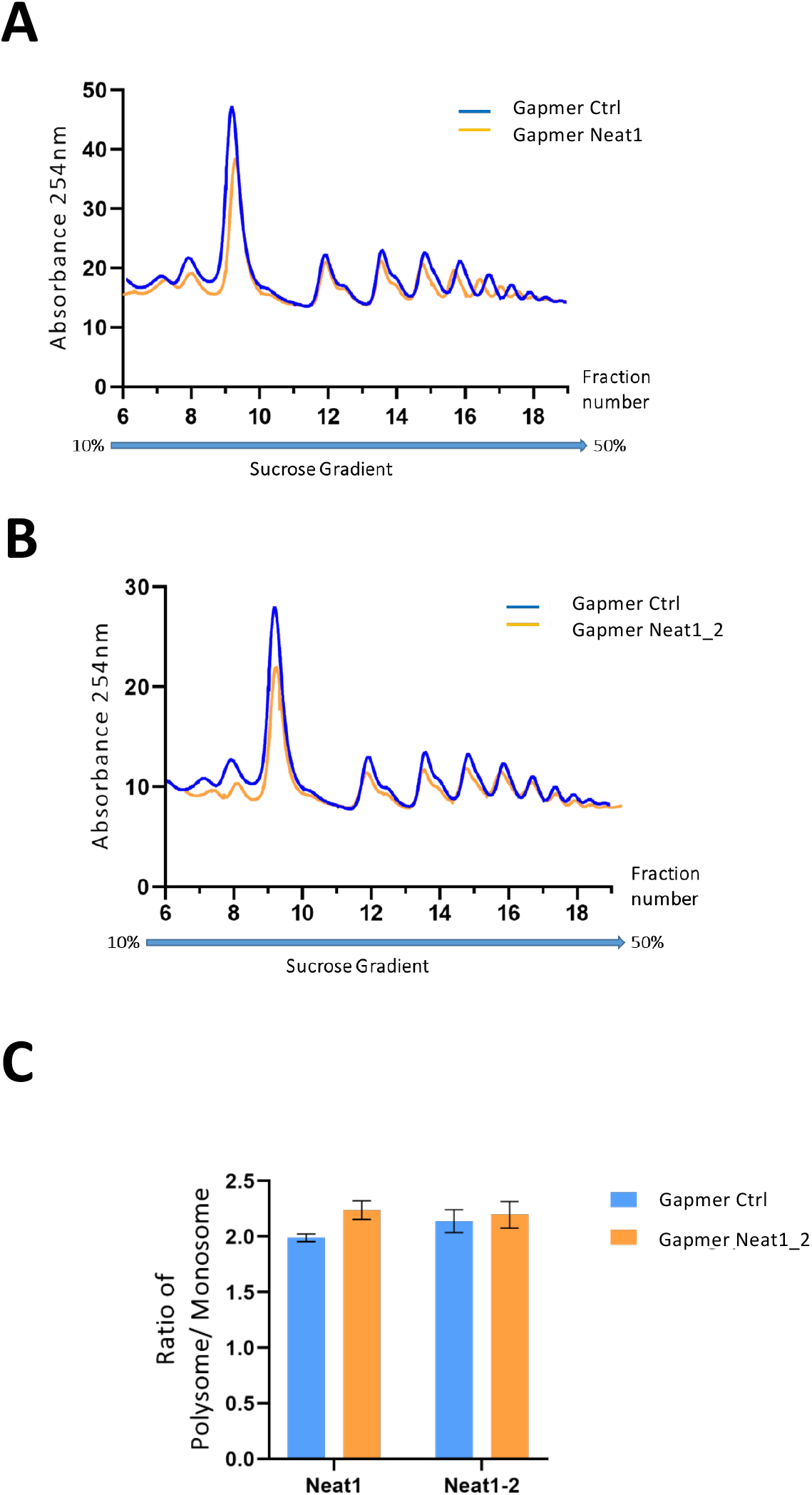
Polysome profiles of HL-1 cardiomyocytes treated by gapmer Neat1 or Neat1_2, compared to gapmer control. In order to isolate translated mRNAs, polysomes from transfected HL-1 cardiomyocytes were purified on a sucrose gradient, as described in Star Methods. Polysomal profiles of HL-1 cardiomyocytes transfected by gapmer Neat1 (A) or Neat1_2 (B) was analyzed and compared to gapmer control. The P/M ratio (polysome/monosome) (C) was determined by delimiting the 80S and polysome peaks, taking the lowest plateau values between each peak and calculating the area under the curve (AUC). Then the sum of area values of the nine polysome peaks was divided by the area of the 80S peak.

**Figure 8-figure supplement 2.**
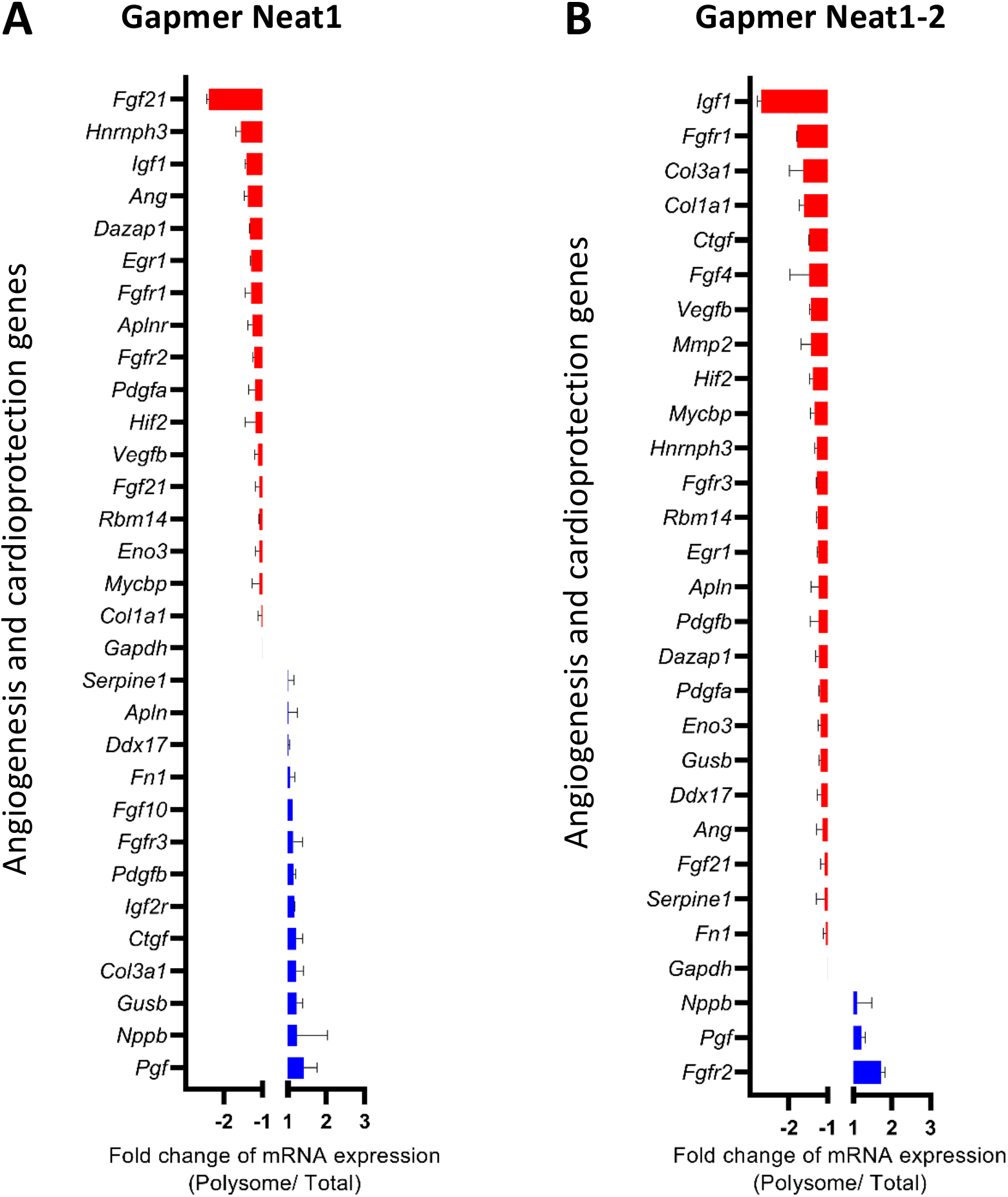
Effect of Neat-1 and Neat1_2 knock-down on translation of mRNAs coding angiogenic and cardioprotective factors. HL-1 cardiomyocytes were transfected with gapmer Neat1, Neat1-2, or control. Polysomes were purified on sucrose gradient as described in Star Methods. Polysome profile is presented in Fig. 8-suppplement 1. RNAs were purified from cytoplasmic extracts and from pooled polysomal fractions and analyzed on a Fluidigm deltagene PCR array from two biologicals replicates (cell culture dishes and cDNAs), each of them measured in three technical replicates (PCR reactions) (Supplementary file 8). Angiogenesis and cardioprotection gene mRNA levels in polysomes (polysomal RNA/ total RNA) were analyzed after knock-down of Neat1 (A) or Neat1_2 (B). Relative quantification (RQ) of mRNA levels was calculated using the 2–Δ ΔCT method with normalization to GAPDH mRNA and to HL-1 tranfected by gapmer control, and is shown as fold change of repression (red) or induction (blue).

## Notes

### Competing Interest Statement

The authors have declared no competing interest.

### Summary of Updates

The revised version contains several modifications requested by eLife reviewers. Author affiliations have been updated. Neat1 impact on IRES-dependent translation has been shown in another cell type, mouse 67NR cell line (Figure 1, suppl. 3). More importantly, smiFISH experiments have been performed and show the colocalization of an IRES-containing mRNA with lncRNA Neat1 in the paraspeckle during hypoxia in HL-1 cardiomyocytes (Fig. 3). The knock-down of Neat1_2 isoform has been performed and shows an impact on IRES-dependent translation, however less important than when knocking-down the two Neat1 isoforms, indicating that they are both involved in the translational control process. Furthermore, FGF1 half-life has been measured after gaper or siRNA treatment, showing no change in FGF1 stability. This reinforce our conclusion about FGF1 mRNA translational control (Fig. 2, Suppl 5-6; Fig. 4, Suppl 3-4).

