## Supplementary files 1-8 for "Long non-coding RNA Neat1 and paraspeckle components are translational regulators in hypoxia": V2 Supplementary file 1 FGF1 cinetique Hx.docx

|  |  |  | Renilla luciferase | | | | | Firefly luciferase | | | | |
| --- | --- | --- | --- | --- | --- | --- | --- | --- | --- | --- | --- | --- |
| **FGF1 IRES** |  | **Exp num** | **A** | **B** | **C** | **Mean** | **SD** | **A** | **B** | **C** | **Mean** | **SD** |
| 4 hr | Normoxia | 1 | 604 174 | 507 891 | 700 457 | 604 174 | 96283 | 77 310 | 91 175 | 63 445 | 77 310 | 13865 |
|  |  | 2 | 452 118 | 345 763 | 472 356 | 423 412 | 68003.37 | 52 728 | 58 279 | 56 285 | 55 764 | 2811.93 |
|  |  | 3 | 495 500 | 529 252 | 547 526 | 524 093 | 26393.94 | 59 485 | 53 665 | 53 223 | 55 458 | 3494.76 |
|  |  | 4 | 6957128 | 8160444 | 8944535 | 8 020 702 | 1001045.67 | 1154339 | 1436444 | 1532732 | 1 374 505 | 196653.61 |
|  | Hypoxia | 1 | 996 143 | 899 215 | 1 093 070 | 996 143 | 96927.5 | 248 088 | 192 876 | 303 300 | 248 088 | 55212 |
|  |  | 2 | 714 978 | 915 991 | 757 182 | 796 050 | 105993.47 | 138 626 | 171 997 | 152 394 | 154 339 | 16770.30 |
|  |  | 3 | 754 720 | 974 680 | 917 680 | 882 360 | 114154.40 | 110 552 | 157 006 | 149 316 | 138 958 | 24898.98 |
|  |  | 4 | 10450445 | 10725351 | 12210832 | 11 128 876 | 947029.52 | 1891657 | 1744526 | 2255783 | 1 963 989 | 263191.62 |
| 8 hr | Normoxia | 1 | 12 580 | 13 628 | 11 532 | 12 580 | 1048 | 3 721 | 5 041 | 2 401 | 3 721 | 1320 |
|  |  | 2 | 700 579 | 710 733 | 712 400 | 707 904 | 6398.15 | 72 622 | 72 505 | 73 578 | 72 902 | 588.63 |
|  |  | 3 | 912 286 | 860 603 | 855 579 | 876 156 | 31390.17 | 121 992 | 112 068 | 132 044 | 122 035 | 9988.06 |
|  |  | 4 | 8896288 | 8402478 | 7130206 | 8 142 991 | 911186.95 | 1772344 | 1668645 | 1416015 | 1 619 001 | 183278.36 |
|  | Hypoxia | 1 | 35 746 | 46 304 | 25 188 | 35 746 | 10558 | 10 760 | 12 896 | 8 624 | 10 760 | 2136 |
|  |  | 2 | 631 364 | 634 140 | 633 140 | 632 881 | 1405.96 | 70 529 | 72 640 | 33 059 | 58 743 | 22267.73 |
|  |  | 3 | 826 664 | 827 015 | 822 262 | 825 314 | 2648.64 | 118 486 | 109 078 | 112 198 | 113 254 | 4792.07 |
|  |  | 4 | 10592865 | 10001462 | 10350135 | 10 314 821 | 297278.83 | 2143188 | 2354330 | 2311918 | 2 269 812 | 111691.19 |
| 24 hr | Normoxia | 1 | 1 010 280 | 1 134 589 | 885 971 | 1 010 280 | 124309 | 231 071 | 198 199 | 263 943 | 231 071 | 32872 |
|  |  | 2 | 1 021 135 | 1 036 550 | 1 020 270 | 1025985 | 9159.77 | 208 454 | 220 996 | 210 670 | 213 373 | 6693.76 |
|  |  | 3 | 956 252 | 896 252 | 789 825 | 880 776 | 84285.87 | 226 971 | 199 587 | 262 787 | 229 782 | 31693.60 |
|  |  | 4 | 8896288 | 8402478 | 7130206 | 8 142 991 | 911186.95 | 1772344 | 1668645 | 1416015 | 1 619 001 | 183278.36 |
|  | Hypoxia | 1 | 662 414 | 716 741 | 608 087 | 662 414 | 54327 | 138 279 | 153 534 | 123 025 | 138 279 | 15254.5 |
|  |  | 2 | 519 324 | 514 720 | 769 988 | 601 344 | 146068.12 | 100 770 | 113 450 | 99 785 | 104 668 | 7621.07 |
|  |  | 3 | 455 239 | 386 245 | 351 256 | 397 580 | 52910.09 | 105 268 | 70 597 | 90 603 | 88 823 | 17403.92 |
|  |  | 4 | 14326454 | 13708075 | 16512683 | 14 849 071 | 1473534.06 | 1733141 | 1936317 | 2026315 | 1 898 591 | 150183.84 |

|  |  |  | Ratio LucF/LucR | | | | |  |  |  |  |  |
| --- | --- | --- | --- | --- | --- | --- | --- | --- | --- | --- | --- | --- |
| **FGF1 IRES** |  | **Exp num** | **A** | **B** | **C** | **Mean** | **SD** | **Normalization** | **Mean** | **SD** | **Mann-Whitney p-value** | **Significance** |
| 4 hr | Normoxia | 1 | 0.128 | 0.1795 | 0.0906 | 0.1327 | 0.04463597 | 1 | **1** | 0.24690202 | 0.0017 | ** |
|  |  | 2 | 0.1166 | 0.1686 | 0.1192 | 0.1348 | 0.02930051 | 1 |  |  |  |  |
|  |  | 3 | 0.1201 | 0.1014 | 0.0972 | 0.10623333 | 0.01219112 | 1 |  |  |  |  |
|  |  | 4 | 0.1659 | 0.176 | 0.1714 | 0.1711 | 0.00505668 | 1 |  |  |  |  |
|  | Hypoxia | 1 | 0.249 | 0.2145 | 0.2775 | 0.247 | 0.03154758 | 1.86134137 | **1.42104619** | 0.28052951 |  |  |
|  |  | 2 | 0.1939 | 0.1878 | 0.2013 | 0.19433333 | 0.00676042 | 1.44164194 |  |  |  |  |
|  |  | 3 | 0.1465 | 0.1611 | 0.1627 | 0.15676667 | 0.00892711 | 1.47568246 |  |  |  |  |
|  |  | 4 | 0.181 | 0.1627 | 0.1847 | 0.17613333 | 0.01177978 | 1.02941749 |  |  |  |  |
| 8 hr | Normoxia | 1 | 0.2958 | 0.3699 | 0.2082 | 0.2913 | 0.08094387 | 1 | **1** | 0.44874634 | 0.6192 | ns |
|  |  | 2 | 0.1037 | 0.102 | 0.1033 | 0.103 | 0.00088882 | 1 |  |  |  |  |
|  |  | 3 | 0.1337 | 0.1302 | 0.1543 | 0.1394 | 0.0130219 | 1 |  |  |  |  |
|  |  | 4 | 0.1992 | 0.1986 | 0.1986 | 0.1988 | 0.00034641 | 1 |  |  |  |  |
|  | Hypoxia | 1 | 0.301 | 0.2785 | 0.3424 | 0.3073 | 0.0324125 | 1.05492619 | **1.03440273** | 0.48267877 |  |  |
|  |  | 2 | 0.1117 | 0.1145 | 0.0522 | 0.0928 | 0.03518849 | 0.90097087 |  |  |  |  |
|  |  | 3 | 0.1433 | 0.1319 | 0.1365 | 0.13723333 | 0.00573527 | 0.9844572 |  |  |  |  |
|  |  | 4 | 0.2023 | 0.2354 | 0.2234 | 0.22036667 | 0.01675719 | 1.10848424 |  |  |  |  |
| 24 hr | Normoxia | 1 | 0.2287 | 0.1747 | 0.2979 | 0.23376667 | 0.06175608 | 1 | **1** | 0.20029197 | 0.1239 | ns |
|  |  | 2 | 0.2041 | 0.2132 | 0.2065 | 0.20793333 | 0.00471628 | 1 |  |  |  |  |
|  |  | 3 | 0.2374 | 0.2227 | 0.3327 | 0.26426667 | 0.05971904 | 1 |  |  |  |  |
|  |  | 4 | 0.1992 | 0.1986 | 0.1986 | 0.1988 | 0.00034641 | 1 |  |  |  |  |
|  | Hypoxia | 1 | 0.2088 | 0.2142 | 0.2023 | 0.20843333 | 0.00595847 | 0.89162983 | **0.82017463** | 0.20416592 |  |  |
|  |  | 2 | 0.194 | 0.2204 | 0.1296 | 0.18133333 | 0.04670646 | 0.87207438 |  |  |  |  |
|  |  | 3 | 0.2312 | 0.1828 | 0.2579 | 0.22396667 | 0.03806893 | 0.84750252 |  |  |  |  |
|  |  | 4 | 0.121 | 0.1413 | 0.1227 | 0.12833333 | 0.01126159 | 0.64553991 |  |  |  |  |

**Supplementary file 1. IRES activities in normoxic and hypoxic HL-1 cells after Neat1 knock-down.**

HL-1 cells were transduced with Lucky Luke bicistronic lentivector containing the IRES of FGF1. Cells were submitted to normoxia or hypoxia 1% O_2_ during 4 hr, 8 hr or 24 hr_._ Renilla and firefly luciferase activities were measured (page 1) and the IRES activities evaluated with the ratio LucF/LucR (page 2). Mann-Whitney test was performed with n=12. **p<0.01. For each hypoxia condition the mean of the LucF/LucR ratio has been calculated with 12 cell culture biological replicates, normalized to normoxia (Experiments A, B, C correspond to experiments performed at different dates, while 1, 2, 3, 4 are experiments performed in parallel at the same date, each of them being already the mean of three technical replicates (36 technical replicates in total).
