## Supplementary files 1-8 for "Long non-coding RNA Neat1 and paraspeckle components are translational regulators in hypoxia": V2 Supplementary file 2 probes primers ans siRNAs.docx

**A/ List of FISH and smiFISH probes**

| **FISH probe name** | **Probe sequence (5’ to 3’)** |
| --- | --- |
| NEAT1-A | [Cyanine3]GGCTGTGAGCCCACCCCAGGGTCCACACTCACGTCCCG |
| NEAT1-B | [Cyanine3]CCCAAGCTTGATCCATAACTCAGGTTTCACCTTGGAAGGC |
| NEAT1-C | [Cyanine3]TTTTACAAAGTGTTATCTAGTAAGATGACCCAGGGCCC |
| NEAT1_2-A | [Cyanine3]GCACCTAAGGCATGGACATGACCTCGGTTAATGCTTTCAC |
| NEAT1_2-B | [Cyanine3]GGAATAATACAGACTTTCTGACGTTGGATCTGGCAATG |

| **smiFISH FLAP name** | **FLAP probe sequence (5’ to 3’)** |
| --- | --- |
| FLAP Y-Cy5 | [Cyanine5]AATGCATGTCGACGAGGTCCGAGTGTAA[Cyanine5] |
| FLAP X-Cy3 | [Cyanine3] CACTGAGTCCAGCTCGAAACTTAGGAGG [Cyanine3] |
| **Neat1 primary probes** | **Primary probe sequence (5’ to 3’)** |
| ACG33 | AATGAGCGATCTTGCCCTGGTCACACTTACACTCGGACCTCGTCGACATGCATT |
| ACG34 | AATTCCTGTATGTGTTGTATAAGGCATGACTCTTACACTCGGACCTCGTCGACATGCATT |
| ACG35 | CTGCATCTCAAAGTAGTGCCACCTGCTTACACTCGGACCTCGTCGACATGCATT |
| ACG36 | AACTCAGGTTTCACCTTGGAAGGCCATTTTACACTCGGACCTCGTCGACATGCATT |
| ACG37 | TCAGGACACTGTCATCTGAGCAGGGCTTACACTCGGACCTCGTCGACATGCATT |
| ACG38 | CTGGTATACACAGCCAGGCCCTAAGTTTACACTCGGACCTCGTCGACATGCATT |
| ACG39 | GGAGAGACCATTCCCAGGGACCACTTTTACACTCGGACCTCGTCGACATGCATT |
| ACG40 | TACCAAGTCGGCAGAATTTGTGGCTAACTTTTACACTCGGACCTCGTCGACATGCATT |
| ACG41 | GCAGCACGCTGCGCAGACGTTAAAGTTTACACTCGGACCTCGTCGACATGCATT |
| ACG42 | CCTACACCTTACGCAATCTTCTCGAGGTTACACTCGGACCTCGTCGACATGCATT |
| ACG43 | TGGAGGCTTGTTTAGAAGATGCAGCAGTCGTTACACTCGGACCTCGTCGACATGCATT |
| ACG44 | CTCCTGAGCACAGGGCTTCCTGAGAATTACACTCGGACCTCGTCGACATGCATT |
| ACG45 | CCATCTGCTTGCCATGGTGAACACTATTACACTCGGACCTCGTCGACATGCATT |
| ACG46 | TTACCGGACGACCCAGGGGCAAGGTTTTACACTCGGACCTCGTCGACATGCATT |
| ACG47 | CCAGCGAAGCTAAGTCCACCTTCCATTTACACTCGGACCTCGTCGACATGCATT |
| ACG48 | CAACTGCTGCTCTGCCACTCAAGTTATTACACTCGGACCTCGTCGACATGCATT |
| ACG49 | TCCTGGACGTCAAGTGCCAGCAGACATTACACTCGGACCTCGTCGACATGCATT |
| ACG50 | AGGGATAGCCTGGTCTTGAAAGCTGCTTTACACTCGGACCTCGTCGACATGCATT |
| ACG51 | CAAGAGCGAGCCCTCCTTGTCACTAATTACACTCGGACCTCGTCGACATGCATT |
| ACG52 | GCGACCAACCTGCAGTTACAGCTTTCTTACACTCGGACCTCGTCGACATGCATT |
| ACG53 | AGTCTCTTTTATTTCCCAAACTGCTTTGACTTTACACTCGGACCTCGTCGACATGCATT |
| ACG54 | CAAGACCCTGTCCCCTTGTCACAATGTTACACTCGGACCTCGTCGACATGCATT |
| ACG55 | ACAACGGGAGGCGTTTACAAGCCCCTTTACACTCGGACCTCGTCGACATGCATT |
| **NEAT1_2 primary probes** | **Primary probe sequence (5’ to 3’)** |
| ACG56 | CCAGCTTCACTTCTTGGCAATCAGCTTTACACTCGGACCTCGTCGACATGCATT |
| ACG57 | CCACGGCTTCCAACTTTATGCCAATATTACACTCGGACCTCGTCGACATGCATT |
| ACG58 | CTGGCTACCTATGTCACACAGATGAGTTACACTCGGACCTCGTCGACATGCATT |
| ACG59 | GGCACTGCCTTGGCTTGGAAATGTAATTACACTCGGACCTCGTCGACATGCATT |
| ACG60 | TCCAGGTGGGAAACTGAGGGTCTGAGTTACACTCGGACCTCGTCGACATGCATT |
| ACG61 | CCATGCTGAGAAGAGGGTCCTCTCAATTACACTCGGACCTCGTCGACATGCATT |
| ACG62 | CATACTGGAGTGAGTTCCAGGACAGCTTACACTCGGACCTCGTCGACATGCATT |
| ACG63 | CACACACACTCTGAGGTTTGTGGACTTTACACTCGGACCTCGTCGACATGCATT |
| ACG64 | CAGCCTGAGAGGTACTCAGAGAGCTTTTACACTCGGACCTCGTCGACATGCATT |
| ACG65 | CTTCTAGGTATGTCCTCAACCCAAGGTTACACTCGGACCTCGTCGACATGCATT |
| ACG66 | CGTCACATGATCTCAGTGTTAGTAGCTAGTTACACTCGGACCTCGTCGACATGCATT |
| ACG67 | GTGAGATGTAGGTAAGGCTGTAGAGGACGGTTACACTCGGACCTCGTCGACATGCATT |
| ACG68 | CACTGGAGGGGTTGTTCCAATGTGACAAATTACACTCGGACCTCGTCGACATGCATT |
| ACG69 | CATGAGTTCAAAGCTGGCCAGGGCTATTACACTCGGACCTCGTCGACATGCATT |
| ACG70 | TTCTGCAGTTCAGACTCCATTCCAACTTACACTCGGACCTCGTCGACATGCATT |
| ACG71 | ACCTTCTGAGGCCTGTTCCAGTGAAATTACACTCGGACCTCGTCGACATGCATT |
| ACG72 | TGGGTCTGCTTTGAAACCATGGCCAATTACACTCGGACCTCGTCGACATGCATT |
| ACG73 | GGTTTCAAGTGGCCTCCTCAAACACGTTACACTCGGACCTCGTCGACATGCATT |
| ACG74 | CATTAGCAGAGGTGTGGGGACAGAGCTTTACACTCGGACCTCGTCGACATGCATT |
| ACG75 | TTCTTGTTAAAGGTCAGACTGGATGTCTTCATTACACTCGGACCTCGTCGACATGCATT |
| ACG76 | GTGTCATGGGACACCATAGCTGTGTATTACACTCGGACCTCGTCGACATGCATT |
| ACG77 | TCTACTGCATGCATGTGGTACACAGATTACACTCGGACCTCGTCGACATGCATT |
| ACG78 | CGGCCAGTAGGGGCTGACTAGTAAGATTACACTCGGACCTCGTCGACATGCATT |
| ACG79 | TCAGCTGTTGCTCTAGAAGGCACATATTACACTCGGACCTCGTCGACATGCATT |
| ACG80 | TGAAGTGTTGGATGTTAACTCAGCATCTTGTTTACACTCGGACCTCGTCGACATGCATT |
| ACG81 | TCAAGGCGCACGTCTGCATATGGAAGTTACACTCGGACCTCGTCGACATGCATT |
| **Bicistronic mRNA primary probes** | **Primary probe sequence (5’ to 3’)** |
| ACG87 | CTCGCTCAACGAACGATTTGATATATTTTCCCCCTCCTAAGTTTCGAGCTGGACTCAGTG |
| ACG88 | CGCGCTACTGGCTCAATATGTGGCACAACCTCCTAAGTTTCGAGCTGGACTCAGTG |
| ACG89 | TCACCTTTCTCTTTGAATGGTTCAAGATATGCCCTCCTAAGTTTCGAGCTGGACTCAGTG |
| ACG90 | TCTCATGATTTTTGATGGCAACATGGTTTCCACCTCCTAAGTTTCGAGCTGGACTCAGTG |
| ACG91 | TCAGCGTGAACTATTGCTTTGATCTTATCTTGCCTCCTAAGTTTCGAGCTGGACTCAGTG |
| ACG92 | CACTGCGGACCAGTTATCATCCGTTTCCTTCCTCCTAAGTTTCGAGCTGGACTCAGTG |
| ACG93 | ATTCAGTATTAGGAAACTTCTTGGCACCTTCACCTCCTAAGTTTCGAGCTGGACTCAGTG |
| ACG94 | ACAACGTCAGGTTTACCACCTTTTACTAACGGCCTCCTAAGTTTCGAGCTGGACTCAGTG |
| ACG95 | TTTCACGAGGCCATGATAATGTTGGACGACGACCTCCTAAGTTTCGAGCTGGACTCAGTG |
| ACG96 | TGATCAACGCAATATCTTCTTCAATATCAGGCCCTCCTAAGTTTCGAGCTGGACTCAGTG |
| ACG97 | GCTCATAGCTATAATGAAATGCCAAACAAGCACCTCCTAAGTTTCGAGCTGGACTCAGTG |
| ACG98 | GCTCATAGCTATAATGAAATGCCAAACAAGCACCTCCTAAGTTTCGAGCTGGACTCAGTG |
| ACG99 | TTTGCCCATACCAATAAGGTCTGGTATAATACCCTCCTAAGTTTCGAGCTGGACTCAGTG |
| ACG100 | TGTCGCCATAAATAAGAAGAGGCCGCGTTACCCCTCCTAAGTTTCGAGCTGGACTCAGTG |
| ACG101 | ATCAAGAACATTCATTTGTTTACATCTGGCCCCCTCCTAAGTTTCGAGCTGGACTCAGTG |
| ACG102 | TCAGGTGCATCTTCTTGCGAAAAATGAAGACCCCTCCTAAGTTTCGAGCTGGACTCAGTG |
| ACG103 | ATAGCATTGGAAAAGAATCCTGGGTCCGATTCCCTCCTAAGTTTCGAGCTGGACTCAGTG |
| ACG104 | TTCATCCCATGATTCAATCACATCTACTACACCCTCCTAAGTTTCGAGCTGGACTCAGTG |
| ACG105 | CCAATCATGGCCGACAAAAATGATCTTCTTTGCCTCCTAAGTTTCGAGCTGGACTCAGTG |
| ACG106 | GTGTTGGAGCAAGATGGATTCCAATTCAGCCTCCTAAGTTTCGAGCTGGACTCAGTG |
| ACG107 | TCAGAGACTTCAGGCGGTCAACGATGCCTCCTAAGTTTCGAGCTGGACTCAGTG |
| ACG108 | CTCCAGAATGTAGCCATCCATCCTTGCCTCCTAAGTTTCGAGCTGGACTCAGTG |
| ACG109 | TCACACACAGTTCGCCTCTTTGATTACCTCCTAAGTTTCGAGCTGGACTCAGTG |
| ACG110 | CTTGCCTGATACCTGGCAGATGGAACCCTCCTAAGTTTCGAGCTGGACTCAGTG |
| ACG111 | AATCGTATTTGTCAATCAGAGTGCTTTTGGCCTCCTAAGTTTCGAGCTGGACTCAGTG |
| ACG112 | CGCACTTTGAATCTTGTAATCCTGAAGGCCTCCTAAGTTTCGAGCTGGACTCAGTG |
| ACG113 | CATGCGAGAATCTCACGCAGGCAGTTCCTCCTAAGTTTCGAGCTGGACTCAGTG |
| ACG114 | CATACGACGATTCTGTGATTTGTATTCAGCCCTCCTAAGTTTCGAGCTGGACTCAGTG |
| ACG115 | ATATCGTTTCATAGCTTCTGCCAACCCCTCCTAAGTTTCGAGCTGGACTCAGTG |
| ACG116 | AGCAATTGTTCCAGGAACCAGGGCGTCCTCCTAAGTTTCGAGCTGGACTCAGTG |
| ACG117 | TCTCTCTGATTTTTCTTGCGTCGAGTTTCCTCCTAAGTTTCGAGCTGGACTCAGTG |
| ACG118 | CGGTAAGACCTTTCGGTACTTCGTCCCCTCCTAAGTTTCGAGCTGGACTCAGTG |
| ACG119 | GCGGTTGTTACTTGACTGGCGACGTACCTCCTAAGTTTCGAGCTGGACTCAGTG |
| ACG120 | CCACGATCTCTTTTTCCGTCATCGTCCCTCCTAAGTTTCGAGCTGGACTCAGTG |
| ACG121 | CTGCGACACCTGCGTCGAAGATGTTGCCTCCTAAGTTTCGAGCTGGACTCAGTG |
| ACG122 | GAAGTGTTCGTCTTCGTCCCAGTAAGCCCTCCTAAGTTTCGAGCTGGACTCAGTG |
| ACG123 | AATCAAGGCGTTGGTCGCTTCCGGATCCTCCTAAGTTTCGAGCTGGACTCAGTG |
| ACG124 | CAGCGTTTTCCCGGTATCCAGATCCACCTCCTAAGTTTCGAGCTGGACTCAGTG |
| ACG125 | TGCAATTGTCTTGTCCCTATCGAAGGCCTCCTAAGTTTCGAGCTGGACTCAGTG |
| ACG126 | TCATTATAAATGTCGTTCGCGGGCGCCCTCCTAAGTTTCGAGCTGGACTCAGTG |
| ACG127 | CTCTTCATAGCCTTATGCAGTTGCTCCCTCCTAAGTTTCGAGCTGGACTCAGTG |
| ACG128 | CAGCGGTTCCATCTTCCAGCGGATAGCCTCCTAAGTTTCGAGCTGGACTCAGTG |
| ACG129 | TGGCGCCGGGCCTTTCTTTATGTTTTCCTCCTAAGTTTCGAGCTGGACTCAGTG |
| ACG130 | TTTCCGCCCTTCTTGGCCTTTATGAGCCTCCTAAGTTTCGAGCTGGACTCAGTG |
| ACG131 | AAACACAACTCCTCCGCGCAACTTTTCCTCCTAAGTTTCGAGCTGGACTCAGTG |
| ACG132 | GGAGCCACCTGATAGCCTTTGTACTTCCTCCTAAGTTTCGAGCTGGACTCAGTG |
| ACG133 | TTTACATAACCGGACATAATCATAGGACCCCTCCTAAGTTTCGAGCTGGACTCAGTG |
| ACG134 | CGGTTTATCATCCCCCTCGGGTGTAACCTCCTAAGTTTCGAGCTGGACTCAGTG |

**B/ List of qPCR primers and cloning primers**

| **Primer name** | **Sequence (5’ to 3’)** |
| --- | --- |
| **qPCR/ddPCR primers** | |
| FGF1 F | TGGACACCGAAGGGCTTTTA |
| FGF1 R | GCATGCTTCTTGGAGGTGTAA |
| NEAT1_2 F | CGGCGGTAACTGTCTTCTGT |
| NEAT1_2 R | GGACGAGGGGCATAGCATAC |
| NEAT1 F | CGACTGCTGCATCTTCTAAACA |
| NEAT1 R | TTTACAACCCAACAGCTTTCCC |
| PSF/SFPQ F | TGAAAAGCTGGCCCAGAAGAA |
| PSF/SFPQ R | TGTGCCATGCTGAGCAAAAC |
| Rps2 F | GGTTCAAGGCTTTCGTCGC |
| Rps2 R | ATGGAACAGTGTGGGGCTTG |
| Nucleolin F | TTGGTGGAGCCGAAGTCAC |
| Nucleolin R | GGTTTTGCCAGCCTTTGCGA |
| hnRNPM F | GATGCCAACGATCTGAGCAAA |
| hnRNPM R | CCAAATCCTATGCCTTCCATTCC |
| **Cloning primers** | |
| HA-P54nrb F | GTCGACATGagcTACCCATACGATGTTCCTGACTATGCGGGCggcgccTATCCCTATGACGTCCCGGATTACGCAGGAATGCAGAGTAATAAAACTTTTAACTTGGAGAAGC |
| HA-P54nrb R | GGATCCACTAGTTTAGTATCGGCGAC |
| miR-Neat1-G2 Top strand | gtcgac*CCTGGAGGCTTGCTGAAGGCTGTA*TGCTGACAAGCTGACTTCAACGATGGGTTTTGGCCACTGACTGACCCATCGTTAGTCAGCTTGT*CAGGACACAAGGCCTGTTACTAGCACTCACATGGAACAAATGGCC*ggatcc |
| miR-Neat1-G2 Bottom strand | ggatccGGCCATTTGTTCCATGTGAGTGCTAGTAACAGGCCTTGTGTCCTGACAAGCTGACTAACGATGGGTCAGTCAGTGGCCAAAACCCATCGTTGAAGTCAGCTTGTCAGCATACAGCCTTCAGCAAGCCTCCAGGgtcgac |
| miR-Neat1_2-G6 Top strand | tcgac*CCTGGAGGCTTGCTGAAGGCTGTA*TGCTGTAAAGTTGCCGGCGGCACAAAGTTTTGGCCACTGACTGACTTTGTGCCCGGCAACTTTA*CAGGACACAAGGCCTGTTACTAGCACTCACATGGAACAAATGGCC* g |
| miR-Neat1_2-G6 Bottom strand | **gatcc**GGCCATTTGTTCCATGTGAGTGCTAGTAACAGGCCTTGTGTCCTGTAAAGTTGCCGGGCACAAAGTCAGTCAGTGGCCAAAACTTTGTGCCGCCGGCAACTTTACAGCATACAGCCTTCAGCAAGCCTCCAGG**g** |
| miR-Neat1_2-G7 Top strand | tcgac*CCTGGAGGCTTGCTGAAGGCTGTA*TGCTGAGAACATGATCTCACAGTACAGTTTTGGCCACTGACTGACTGTACTGTGATCATGTTCT*CAGGACACAAGGCCTGTTACTAGCACTCACATGGAACAAATGGCC* g |
| miR-Neat1_2-G7 Bottom strand | **gatcc**GGCCATTTGTTCCATGTGAGTGCTAGTAACAGGCCTTGTGTCCTGAGAACATGATCACAGTACAGTCAGTCAGTGGCCAAAACTGTACTGTGAGATCATGTTCTCAGCATACAGCCTTCAGCAAGCCTCCAGG**g** |

**C/ List of LNA gapmers used in NEAT1 knockdown assays**

| **Gapmer Name and reference** | **Sequence (5’ to 3’)** |
| --- | --- |
| NEAT1 A (LG00218175) | A*C*G*T*C*C*A*T*G*A*A*G*C*A*T*T |
| NEAT1 B (LG00218176) | A*T*C*A*G*C*C*T*T*T*A*G*A*T*T*T |
| NEAT1 C (LG00218177) | A*G*A*A*G*A*T*G*C*A*G*C*A*G*T*C |
| NEAT1 D (LG00218178) | G*G*C*A*A*A*G*C*A*C*T*C*A*T*G*A |
| NEGATIVE CONTROL A (LG00000002) | A*A*C*A*C*G*T*C*T*A*T*A*C*G*C |

**D/ List of siRNA SMARTpools (Dharmacon) used for IRES activity studies**

| **siRNA SMARTpool name** | **Target sequences (5’ to 3’)** |
| --- | --- |
| P54nrb mouse | CUAUUUGACCUGAAGAAUUU  UUUUCAUUCAUAAGGAUAA  GAAUCAUACUCCAAGGAAG  CUAUGGACCAGUUAGAUGA |
| PSPC1 mouse | GGGUUAAGAUUGUAUAAAC  UGUUAAUGUAUGUAGACUA  GUGGCAUUAUUAUUAAGUC  UCACUAUUCAUAUAUUCUG |
| SFPQ mouse | UGAUGAUUCGCCAACGUGA  UGGUGGUGGAACAAUGAAC  CCAACGAGAAGAAAGUUAC  UCAGGAGGCCAGAAAUUUC |
| Nucleolin mouse | CCAUGGAGAUCAGAUUAGU  CUCUGUUCGUGCAAGAAUA  CCAGUGAGUUCAAUUAGUA  CUAUCAAGGAUGUUUGGUU |
| Rps2 mouse | GGAUCAUCUUGUGAAAACC  GUCUUGGUGUUAAGUGCUC  CCGGUAUAGAUGACUGCUA  UUGGGGACUACAAUGGUCA |
| hnRNPM mouse | CCAGUAUCCUAAAUAAUCC  GCAAUAUCUAUGUUUAAUG  GAUUGAUGUUCGAAUUGAU  GCCUUCAUUACAAAUAUAC |

**E/ List of Fluidigm deltagene primers**

| **Target** | **Forward primer 5’ to 3’** | **Reverse primer 5’ to 3’** |
| --- | --- | --- |
| *Aanat* | GAGTTCCGCTGCCTCACA | GAAGTGCCGGATCTCATCCA |
| *Adar (Adar1)* | CGTTGTTGACGCACTTCCTA | CAGATGCCCTTGGCTGAAAA |
| *Alox15* | TGCAGCTCTGGCAAGTCA | AACAGCTTGGTCGGTCTTGTA |
| *Ang* | CCATCTGTGGAGCGAATGGAA | TTGCAAGTGGTGACCTGGAA |
| *Apaf1* | CACAGACCTTTCCATCCTTCA | CGTTTCCAAGTCCCAGAGAA |
| *Apln* | GCAGGAGGAAATTTCGCAGAC | ACTTGGCGAGCCCTTCAA |
| *Aplnr* | TTGACTGGCCTTTTGGAACC | GCAAAAGACACTGGCGTACA |
| *Bag1* | AGGCCACAGGAGTTCCACTA | TGCTGACAACGGTGTTTCCA |
| *Bax* | GCGTGGTTGCCCTCTTCTA | CTGATCAGCTCGGGCACTTTA |
| *Bcl2 (Bcl-2)* | ATGTGTGTGGAGAGCGTCAA | GATGCCGGTTCAGGTACTCA |
| *Bcl2L1 (Bcl-XL)* | AGGCAGGCGATGAGTTTGAA | GGTCCCTGGGGTTATGTGAA |
| *Bip (GRP78/Hspa5)* | TGCTGAGGCGTATTTGGGAA | TCGCTGGGCATCATTGAAGTA |
| *BNP (Nppb)* | CTAGCCAGTCTCCAGAGCAA | GTGAGGCCTTGGTCCTTCA |
| *Casp2 (Caspase-2)* | TCCAAGTCTACAGAACAAGCCAAA | TCCATCTTGCTGGTCGACAC |
| *Cdk1* | TGCCAGAGCGTTTGGAATAC | CACTTCTGGAGATCGGTACCA |
| *Col1A1* | TTCAGGGAATGCCTGGTGAA | ACCTTTGGGACCAGCATCA |
| *Col3A1* | TGCTGGAAAGAATGGGGAGAC | GGTCCAGAATCTCCCTTGTCAC |
| *Csde1* | ATTGGAACGAGCAACCAACA | TCTCTCATGGCAGCAATCAC |
| *Ctgf* | AAGCTGACCTGGAGGAAAACA | TGCAGCCAGAAAGCTCAAAC |
| *Cyr61* | CCACACCAAGGGGTTGGAA | CACAGGGTCTGCCTTCTGAC |
| *Dazap1* | TGCCCCAGACATGAGCAAA | ACTCAAGTCCTGCCCGTAAC |
| *Ddx17* | TGGCTCTTAGTGGCAGGGATA | GAACAATCGCAGGCAGCAAA |
| *Egr1* | ACAACCCTATGAGCACCTGAC | GGCTGGGATAACTCGTCTCC |
| *Elavl1* | TGGGCGAATCATCAACTCCA | GTCAAACCGGATAAAGGCAACC |
| *Eno3* | AAGCTTGGGGAGCTGTACAA | CTGGTCAAAGGGGTCCTCAA |
| *Epas1* | AAGCTTTTCGCCATGGACAC | CAAGGTCTCCAAATCCAGTTCAC |
| *Fgf1* | TGGACACCGAAGGGCTTTTA | GCATGCTTCTTGGAGGTGTAA |
| *Fgf10* | GCGGGACCAAGAATGAAGAC | GTTGCTGTTGATGGCTTTGAC |
| *Fgf2* | TCTTCCTGCGCATCCATCC | GCACACACTCCCTTGATAGACA |
| *Fgf21* | CCTGGGTGTCAAAGCCTCTA | TGCAGGCCTCAGGATCAAA |
| *Fgf23* | AACAGGAGCCATGACTCGAA | CTTGCAATTCTCTGGGCTGAA |
| *Fgf4* | TGGTGTGCACGCAGACAC | GCCACTCCGAAGATGCTCAC |
| *Fgfr1* | GACGGACAACACCAAACCAA | CGCATGCAGTTTCTTCTCCA |
| *Fgfr2* | TCAAGTGGATGGCTCCTGAA | CACATTAACACCCCGAAGGAC |
| *Fgfr3* | ACAAGGTCTCTCGCTTCCC | AAGGGGTGTGTTGGAGTTCA |
| *Fmr1 (Fmrp)* | GGGTGAGGATTGAGGCTGAAA | GGTAGGGAACTTGGTGGCATAA |
| *Fn1 (Fibronectine)* | GGAACCAGCAGAGTCCCAAA | CCTCGGTGTTGTAAGGTGGAA |
| *Fus* | GATACGGCCAGAGCAGCTA | GGGAGCTGACTGAGTTCCATA |
| *Gapdh* | CAAGGTCATCCCAGAGCTGAA | CAGATCCACGACGGACACA |
| *Gusb* | GTTTCGAGCAGCAATGGTAC | ACACCCAGCCAATAAAGTCC |
| *Hgf* | GGGTTTGGCCATGAATTTGAC | GCCTCCTTTACCAATGATGCA |
| *Hif1A* | TCGACACAGCCTCGATATGAA | TTCCGGCTCATAACCCATCA |
| *Hnrnpa1* | AAAAGGGCGAGCAGAAGGTA | GCAGCGTCAAAGAGCAGAAA |
| *Hnrnph3* | GGAAACGATGGCTTTGATGACA | AACTTGCATCACCAGCTCCA |
| *Hnrnpk* | GAAGAGGAAGGCCTGGAGAC | GCTCCATGTGTCAATTGCAGAA |
| *Hnrnpm* | GATGCCAACCATCTGAGCAAA | CCAAATCCTATGCCTTCCATTCC |
| *Hnrnpr* | TTCTGTGGCAAACAACAGAC | GAGAATAACGTCCACCAAACC |
| *Hoxa9* | TGTTTGGTGCCTCGTGGAA | GGTGGTGGTGGTGATGATACA |
| *Igf1* | CACACTGACATGCCCAAGAC | GGTCTTGTTTCCTGCACTTCC |
| *Igf1R* | TTGGGCAATGGAGTGCTGTA | CTCCCATTCATCAGGCACGTA |
| *Igf2R* | TCAACGATTCTGCTCAAGGAC | CAGGCATTGCACCACAGATA |
| *Irf2* | GTGGCTGGAGGAGCAGATAAA | GCATCCAGGGGATCTGGAAAA |
| *Lamb1* | CTGAGCTGTTGCTTGAGGAA | TTCACCATGTCTGCAGTGAC |
| *Lef1* | ACACATCCCGTCAGATGTCA | GGGTAGAAGGTGGGGATTTCA |
| *Mmp2* | CGAGGACTATGACCGGGATA | GGGCACCTTCTGAATTTCCA |
| *Mycbp* | CGCTGTGGCTGTCACTA | CCCGACTTCTCCAAGTACC |
| *Mycl* | GGCGAGCCCAAGACTCA | CCAGAGATCGCCTCTTCTCC |
| *Ncl* | CTGGAGTTGCAAGGATCCAA | AGTGGTATCCTCAGACAGACC |
| *Neat1* | GGGACTTGTGGGAGAAAGCA | GGTTCCAGGCACAATCCTCA |
| *Nfil3* | TACAGCCGCCCTTTCTTTTCC | GTTGTCCGGCACAGGGTAAA |
| *Nkrf (NRF)* | CCAAACCTTCCAAAGGCCAAA | AGGCTCAAAACGAGCTCTCA |
| *Nono* | ACGTCATGAGCACCAGGTTA | TATGCAGCTCCTCCATTCTCC |
| *Nr1D1 (rev-erb-a)* | GACCAGGTGACCCTGCTTAA | TCTGGTCCTTCACGTTGAACA |
| *p16INK4 (Cdkn2A)* | CCGACGGGCATAGCTTCA | GGGCTGAGGCCGGATTTA |
| *p27kip1 (Cdkn1B)* | CAGTGTCCAGGGATGAGGAA | TTCGGGGAACCGTCTGAAA |
| *Pdgfa* | TGTAACACCAGCAGCGTCAA | GGCTTCTTCCTGACATACTCCA |
| *Pdgfb* | CGGCCTGTGACTAGAAGTCC | TCACCGTCCGAATGGTCAC |
| *Per1* | CCTCATGCTGGCCATTCATA | CACAGAAGCGAATAGGGGAA |
| *Pfkm (Pfk1)* | CACCAGAGGACGTTTGTGTTA | ACAGGACAGAGAGGTGACAA |
| *Pgf (Plgf)* | CCAATCGGGATCCACATTTCTA | GCCTTTGTCGTCTCCAGAATA |
| *Prox1* | GCCCTCAACATGCACTACAAC | CGTGATCTGCGCAACTTCC |
| *Pspc1* | TCCCCGTGGAGCAATAAACA | ATACCCATCATTGGAGGAGGAC |
| *Rbm14* | CAGCTACGACGATCCCTACAA | GGTAATCAGAGAGCTCGGCTAA |
| *Rpl10A* | CTGCCTTCTTTCCGGTTTCC | TTCCCGTGCAGGACTTCC |
| *Rps2* | CGCGCTTCTTGGAGCACTATA | TGCACCGGCGTCATCC |
| *Rps25* | CGACAAAGCGACATACGACAA | AGAGACCACGGCTGGAGTA |
| *Rrbp1* | TTGCCCTCTCGCTCTCC | CCCAAGGTCTGAGTGTCGTA |
| *Serpine1 (PAI1)* | GGCTCAGAGCAACAAGTTCA | TGGTAGGGCAGTTCCACAA |
| *Setd7 (Set7)* | GAAAAGAATGGGCGTGGGAA | AGGGCATCGTCCACGTAATA |
| *Sfpq* | TGAAAAGCTGGCCCAGAAGAA | TGTGCCATGCTGAGCAAAAC |
| *Shmt1* | CACTCACAGAGCTAGGCTACA | ACCCTTAGAACGCAGATCCA |
| *Slc7A1 (Cat-1)* | ACAGGCATCATCTGGAGACA | ACAGGAAGTACGGGGACAAA |
| *Srebf1 (Srebp1)* | ACCCTACGAAGTGCACACAA | CACATCTGTGCCTCCTCCA |
| *Sstr2 (Sst2)* | AGCGAGTGCTCGAGGAAAA | CGTCTAGAACCAAGCTGCCTA |
| *Thbd (Thrombomodulin)* | GGCTTCGAATGCTTCTGCTA | AGATCCGAAACACGGATCCA |
| *Trp53* | CACAGCGTGGTGGTACCTTA | CCCATGCAGGAGCTATTACACA |
| *Txnip* | TTCCTGATGGACGTGTGTCA | TCAGCATGGATGGAGATGTCA |
| *Utrn (Utrophin)* | GGGGACGTGAAAGAGATCAA | CTGAGGATGGCGTTGTTCTA |
| *Vash1* | ATGCCTGGAAGCTGTGATCC | AGGTCTTGAAGCTGATGGGAA |
| *Vegfa* | CCAGCACATAGGAGAGATGAG | CTGGCTTTGTTCTGTCTTTCTT |
| *Vegfb* | GCCCCAGCCACCAGAA | GCTGGGCACTAGTTGTTTGAC |
| *Vegfc* | AGACGTTCTCTGCCAGCAA | AGGCATCGGCACATGTAGTTA |
| *Vegfd (Figf)* | TCCATTCAGACCCCAGAAGAA | GTGTTATCCCACAGCATGTCA |
| *Xiap* | CAAGTGAAGACCCTTGGGAAC | TTCTTGCCCCTTCTCATCCA |

**Supplementary file 2. Sequences of FISH and smiFISH probes, PCR primers, gapmers, siRNAs and Fluidigm probes.**

A/ FISH and smiFISH probes.

B/ qPCR primers and cloning primers.

C/ LNA gapmers used in NEAT1 knockdown assays.

D/ SiRNA SMARTpools (Dharmacon) used for IRES activity studies.

E/ Fluidigm deltagene primers. The corresponding genes are indicated in the left column.
