## Supplementary files 1-8 for "Long non-coding RNA Neat1 and paraspeckle components are translational regulators in hypoxia": V2 Supplementary file 3 Neat1 KD IRES activities.docx

|  |  |  | Renilla luciferase | | | | | Firefly luciferase | | | | |
| --- | --- | --- | --- | --- | --- | --- | --- | --- | --- | --- | --- | --- |
| **FGF1 IRES** |  | **Exp num** | **A** | **B** | **C** | **Mean** | **SD** | **A** | **B** | **C** | **Mean** | **SD** |
| Normoxia | Gapmer  Ctrl | 1 | 130797 | 98622 | 82852 | 104090 | 24436 | 33722 | 24899 | 23844 | 27488 | 5424 |
|  |  | 2 | 1324654 | 1485789 | 1482312 | 1430918 | 92044 | 173699 | 206379 | 211312 | 197130 | 20442 |
|  |  | 3 | 1268988 | 1552184 | 1543766 | 1454979 | 161128 | 148045 | 155089 | 183753 | 162296 | 18913 |
|  | Gapmer  Neat1 | 1 | 180118 | 147737 | 159475 | 162443 | 16393 | 29972 | 23539 | 23588 | 25700 | 3700 |
|  |  | 2 | 6341693 | 6755014 | 5578620 | 6225109 | 596800 | 363734 | 390867 | 323563 | 359388 | 33862 |
|  |  | 3 | 5087778 | 4266498 | 4841003 | 4731760 | 421397 | 284280 | 320062 | 273287 | 292543 | 24458 |
| Hypoxia | Gapmer  Ctrl | 1 | 116671 | 64686 | 90595 | 90651 | 25993 | 27125 | 15740 | 17859 | 20242 | 6055 |
|  |  | 2 | 1825907 | 1398828 | 1559373 | 1594702 | 215720 | 167819 | 152079 | 198757 | 172885 | 23748 |
|  |  | 3 | 1884235 | 2561907 | 1797474 | 2081205 | 418554 | 277071 | 360733 | 324190 | 320664 | 41942 |
|  | Gapmer  Neat1 | 1 | 148434 | 175604 | 250437 | 191492 | 52825 | 17478 | 20704 | 23061 | 20414 | 2803 |
|  |  | 2 | 8688776 | 8791901 | 8157013 | 8545897 | 340707 | 405878 | 412598 | 345985 | 388154 | 36673 |
|  |  | 3 | 7787636 | 6838627 | 8432833 | 7686365 | 801913 | 492167 | 545342 | 511643 | 516384 | 26903 |

|  |  |  | Ratio LucF/LucR | | | | |  |  |  |  |  | |
| --- | --- | --- | --- | --- | --- | --- | --- | --- | --- | --- | --- | --- | --- |
|  |  | **Exp num** | **A** | **B** | **C** | **Mean** | **SD** | **Normalization** | **Mean** | **SD** | **Mann-Whitney p-value** | | **Significance** |
| Normoxia | Gapmer  Ctrl | 1 | 0.258 | 0.252 | 0.288 | 0.266 | 0.019 | 1.000 | 1.000 | 0.421 | *0.0469* | | * |
|  |  | 2 | 0.131 | 0.139 | 0.143 | 0.138 | 0.006 | 1.000 |  |  |  |  |  |
|  |  | 3 | 0.117 | 0.100 | 0.119 | 0.112 | 0.010 | 1.000 |  |  |  |  |  |
|  | Gapmer  Neat1 | 1 | 0.166 | 0.159 | 0.148 | 0.158 | 0.009 | 0.593 | 0.539 | 0.288 |  |  |  |
|  |  | 2 | 0.057 | 0.058 | 0.058 | 0.058 | 0.000 | 0.420 |  |  |  |  |  |
|  |  | 3 | 0.056 | 0.075 | 0.056 | 0.062 | 0.011 | 0.558 |  |  |  |  |  |
| Hypoxia | Gapmer  Ctrl | 1 | 0.232 | 0.243 | 0.197 | 0.224 | 0.024 | 1.000 | 1.000 | 0.327 | *0.0006* | | *** |
|  |  | 2 | 0.092 | 0.109 | 0.127 | 0.109 | 0.018 | 1.000 |  |  |  |  |  |
|  |  | 3 | 0.147 | 0.141 | 0.180 | 0.156 | 0.021 | 1.000 |  |  |  |  |  |
|  | Gapmer  Neat1 | 1 | 0.118 | 0.118 | 0.092 | 0.109 | 0.015 | 0.487 | 0.454 | 0.181 |  |  |  |
|  |  | 2 | 0.047 | 0.047 | 0.042 | 0.045 | 0.003 | 0.415 |  |  |  |  |  |
|  |  | 3 | 0.063 | 0.080 | 0.061 | 0.068 | 0.010 | 0.435 |  |  |  |  |  |

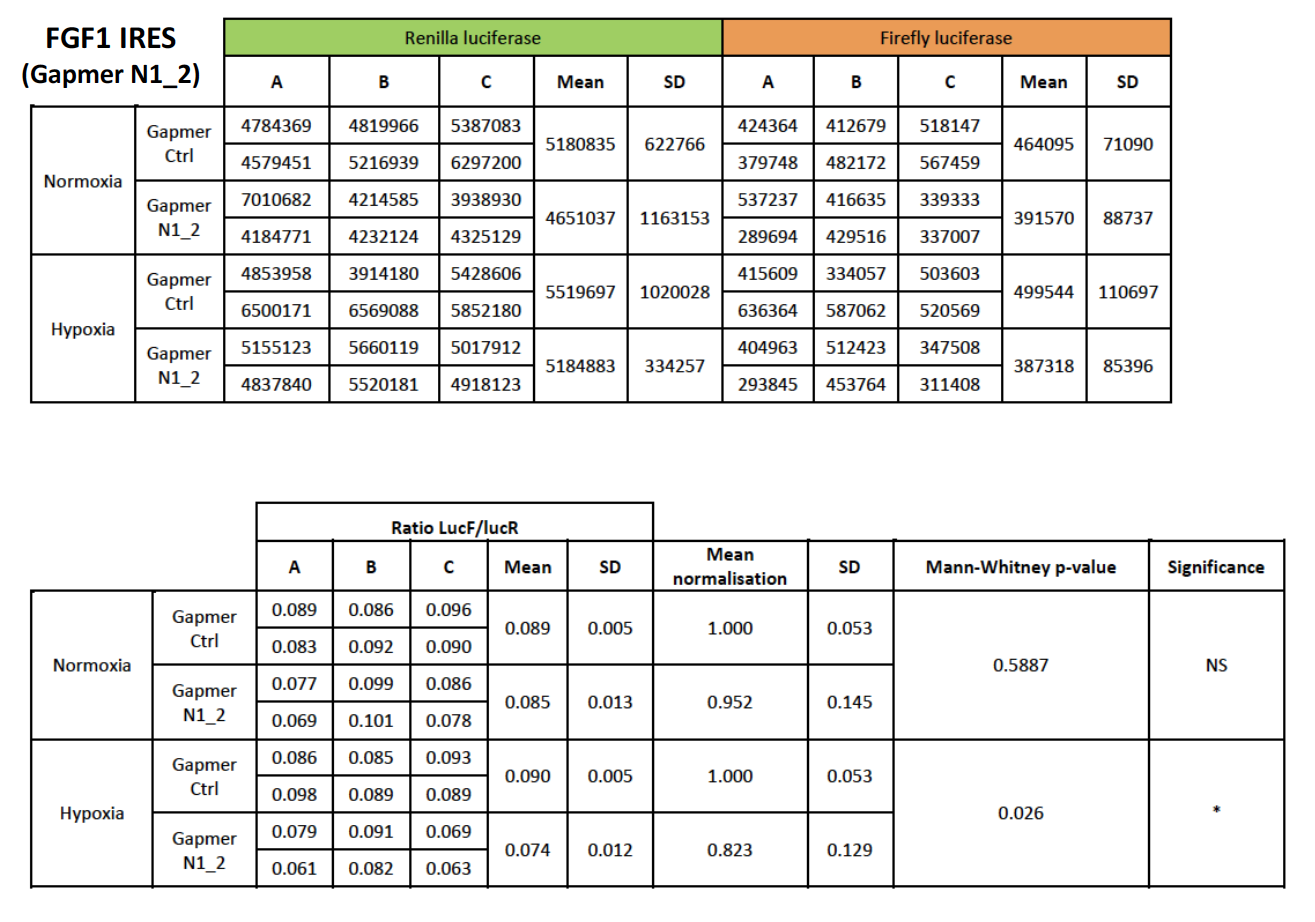

|  |  |  | Renilla luciferase | | | | | Firefly luciferase | | | | |
| --- | --- | --- | --- | --- | --- | --- | --- | --- | --- | --- | --- | --- |
| **FGF2 IRES** |  | **Exp num** | **A** | **B** | **C** | **Mean** | **SD** | **A** | **B** | **C** | **Mean** | **SD** |
| Normoxia | Gapmer  Ctrl | 1 | 56290 | 26712 | 40631 | 41211 | 14798 | 17104 | 7094 | 9014 | 11071 | 5313 |
|  |  | 2 | 32727 | 23049 | 30834 | 28870 | 5129 | 6980 | 6533 | 7575 | 7030 | 523 |
|  |  | 3 | 43246 | 33170 | 73107 | 49841 | 20769 | 4905 | 4152 | 5865 | 4974 | 858 |
|  | Gapmer  Neat1 | 1 | 113254 | 699484 | 191238 | 334659 | 318345 | 10903 | 54564 | 17640 | 27702 | 23505 |
|  |  | 2 | 42957 | 48949 | 36294 | 42733 | 6330 | 5758 | 6556 | 5688 | 6001 | 482 |
|  |  | 3 | 67023 | 86168 | 116483 | 89891 | 24939 | 4890 | 4386 | 6964 | 5413 | 1366 |
| Hypoxia | Gapmer  Ctrl | 1 | 183592 | 223789 | 222880 | 210087 | 22949 | 32155 | 35425 | 42292 | 36624 | 5174 |
|  |  | 2 | 35379 | 12778 | 30119 | 26092 | 11827 | 6056 | 2279 | 5383 | 4573 | 2014 |
|  |  | 3 | 77903 | 83791 | 97087 | 86261 | 9828 | 8422 | 7184 | 1087 | 5564 | 3926 |
|  | Gapmer  Neat1 | 1 | 546229 | 603279 | 160000 | 436503 | 241151 | 49241 | 45909 | 12000 | 35717 | 20606 |
|  |  | 2 | 40391 | 57630 | 46616 | 48212 | 8730 | 3766 | 5063 | 4136 | 4322 | 668 |
|  |  | 3 | 196606 | 161857 | 181246 | 179903 | 17413 | 10066 | 7553 | 7996 | 8538 | 1341 |

|  |  |  | Ratio LucF/LucR | | | | |  |  |  |  |  | |
| --- | --- | --- | --- | --- | --- | --- | --- | --- | --- | --- | --- | --- | --- |
|  |  | **Exp num** | **A** | **B** | **C** | **Mean** | **SD** | **Normalization** | **Mean** | **SD** | **Mann-Whitney p-value** | | **Significance** |
| Normoxia | Gapmer  Ctrl | 1 | 0.304 | 0.266 | 0.222 | 0.264 | 0.041 | 1.000 | 1.000 | 0.391 | *0.0071* | | ** |
|  |  | 2 | 0.213 | 0.283 | 0.246 | 0.247 | 0.035 | 1.000 |  |  |  |  |  |
|  |  | 3 | 0.113 | 0.125 | 0.080 | 0.106 | 0.023 | 1.000 |  |  |  |  |  |
|  | Gapmer  Neat1 | 1 | 0.096 | 0.078 | 0.092 | 0.089 | 0.010 | 0.337 | 0.472 | 0.178 |  |  |  |
|  |  | 2 | 0.134 | 0.134 | 0.157 | 0.142 | 0.013 | 0.572 |  |  |  |  |  |
|  |  | 3 | 0.073 | 0.051 | 0.060 | 0.061 | 0.011 | 0.576 |  |  |  |  |  |
| Hypoxia | Gapmer  Ctrl | 1 | 0.175 | 0.158 | 0.190 | 0.174 | 0.016 | 1.000 | 1.000 | 0.428 | *0.0142* | | * |
|  |  | 2 | 0.171 | 0.178 | 0.179 | 0.176 | 0.004 | 1.000 |  |  |  |  |  |
|  |  | 3 | 0.108 | 0.086 | 0.011 | 0.097 | 0.016 | 1.000 |  |  |  |  |  |
|  | Gapmer  Neat1 | 1 | 0.090 | 0.076 | 0.075 | 0.080 | 0.008 | 0.461 | 0.520 | 0.143 |  |  |  |
|  |  | 2 | 0.093 | 0.088 | 0.089 | 0.090 | 0.003 | 0.511 |  |  |  |  |  |
|  |  | 3 | 0.051 | 0.047 | 0.044 | 0.047 | 0.004 | 0.488 |  |  |  |  |  |

|  |  |  | Renilla luciferase | | | | | Firefly luciferase | | | | |
| --- | --- | --- | --- | --- | --- | --- | --- | --- | --- | --- | --- | --- |
| **VEGFAa IRES** |  | **Exp num** | **A** | **B** | **C** | **Mean** | **SD** | **A** | **B** | **C** | **Mean** | **SD** |
| Normoxia | Gapmer  Ctrl | 1 | 228280 | 264370 | 254890 | 249180 | 18710 | 392 | 426 | 428 | 415 | 21 |
|  |  | 2 | 345732 | 223452 | 336037 | 340884 | 67973 | 1188 | -78 | 995 | 1091 | 682 |
|  |  | 3 | 59329 | 46352 | 58549 | 54743 | 7278 | 443 | 414 | 509 | 455 | 49 |
|  | Gapmer  Neat1 | 1 | 850495 | 892309 | 696255 | 813020 | 103260 | 1961 | 2433 | 1916 | 2103 | 286 |
|  |  | 2 | 888941 | 1182727 | 977300 | 1016323 | 150730 | 2060 | 3309 | 2433 | 2601 | 641 |
|  |  | 3 | 137134 | 125505 | 129468 | 130702 | 5912 | 708 | 570 | 519 | 599 | 98 |
| Hypoxia | Gapmer  Ctrl | 1 | 454068 | 420879 | 488434 | 454460 | 33779 | 3371 | 3252 | 3749 | 3457 | 259 |
|  |  | 2 | 311252 | 476952 | 336648 | 374951 | 89244 | 4018 | 4420 | 4405 | 4281 | 228 |
|  |  | 3 | 125829 | 106517 | 117675 | 116674 | 9695 | 796 | 769 | 896 | 820 | 67 |
|  | Gapmer  Neat1 | 1 | 1755725 | 1786388 | 1571152 | 1704422 | 116429 | 6368 | 5851 | 4420 | 5546 | 1009 |
|  |  | 2 | 1518298 | 885182 | 1006695 | 1136725 | 335990 | 7792 | 5082 | 5605 | 6160 | 1438 |
|  |  | 3 | 239161 | 234022 | 237766 | 236983 | 2657 | 1243 | 1011 | 919 | 1058 | 167 |

|  |  |  | Ratio LucF/LucR | | | | |  |  |  |  |  | |
| --- | --- | --- | --- | --- | --- | --- | --- | --- | --- | --- | --- | --- | --- |
|  |  | **Exp num** | **A** | **B** | **C** | **Mean** | **SD** | **Normalization** | **Mean** | **SD** | **Mann-Whitney p-value** | | **Significance** |
| Normoxia | Gapmer  Ctrl | 1 | 0.002 | 0.002 | 0.002 | 0.002 | 0.000 | 1.000 | 1.000 | 0.858 | *> 0.9999* | | ns |
|  |  | 2 | 0.003 | 0.000 | 0.003 | 0.003 | 0.002 | 1.000 |  |  |  |  |  |
|  |  | 3 | 0.007 | 0.009 | 0.009 | 0.008 | 0.001 | 1.000 |  |  |  |  |  |
|  | Gapmer  Neat1 | 1 | 0.002 | 0.003 | 0.003 | 0.003 | 0.000 | 1.554 | 0.805 | 0.264 |  |  |  |
|  |  | 2 | 0.002 | 0.003 | 0.002 | 0.003 | 0.000 | 0.793 |  |  |  |  |  |
|  |  | 3 | 0.005 | 0.005 | 0.004 | 0.005 | 0.001 | 0.547 |  |  |  |  |  |
| Hypoxia | Gapmer  Ctrl | 1 | 0.007 | 0.008 | 0.008 | 0.008 | 0.000 | 1.000 | 1.000 | 0.283 | *0.0001* | | *** |
|  |  | 2 | 0.013 | 0.009 | 0.013 | 0.012 | 0.002 | 1.000 |  |  |  |  |  |
|  |  | 3 | 0.006 | 0.007 | 0.008 | 0.007 | 0.001 | 1.000 |  |  |  |  |  |
|  | Gapmer  Neat1 | 1 | 0.004 | 0.003 | 0.003 | 0.003 | 0.000 | 0.426 | 0.499 | 0.121 |  |  |  |
|  |  | 2 | 0.005 | 0.006 | 0.006 | 0.005 | 0.000 | 0.466 |  |  |  |  |  |
|  |  | 3 | 0.005 | 0.004 | 0.004 | 0.004 | 0.001 | 0.632 |  |  |  |  |  |

|  | |  | Renilla luciferase | | | | | Firefly luciferase | | | | |
| --- | --- | --- | --- | --- | --- | --- | --- | --- | --- | --- | --- | --- |
| **VEGFAb IRES** |  | **Exp num** | **A** | **B** | **C** | **Mean** | **SD** | **A** | **B** | **C** | **Mean** | **SD** |
| Normoxia | Gapmer  Ctrl | 1 | 592130 | 588445 | 584899 | 588491 | 3616 | 89024 | 83789 | 81707 | 84840 | 3770 |
|  |  | 2 | 1154940 | 754768 | 976328 | 962012 | 200470 | 160743 | 101525 | 127989 | 130085 | 29665 |
|  |  | 3 | 1393867 | 283626 | 301647 | 659714 | 635860 | 251094 | 80519 | 74070 | 135228 | 100395 |
|  | Gapmer  Neat1 | 1 | 2164817 | 2214347 | 2242314 | 2207159 | 39245 | 192780 | 214978 | 207359 | 205039 | 11279 |
|  |  | 2 | 3195306 | 3809065 | 3545090 | 3516487 | 307878 | 246312 | 305725 | 315591 | 289209 | 37476 |
|  |  | 3 | 4299224 | 3987335 | 4317327 | 4201296 | 185516 | 508352 | 405949 | 464525 | 459608 | 51378 |
| Hypoxia | Gapmer  Ctrl | 1 | 1001414 | 800377 | 1089184 | 963658 | 148059 | 121784 | 102844 | 139794 | 121474 | 18477 |
|  |  | 2 | 1667877 | 1751303 | 1499475 | 1639552 | 128281 | 276036 | 273662 | 246016 | 265238 | 16689 |
|  |  | 3 | 2113072 | 1572673 | 1587910 | 1757885 | 307695 | 365933 | 451063 | 420135 | 412377 | 43092 |
|  | Gapmer  Neat1 | 1 | 3813545 | 4028730 | 4229566 | 4023947 | 208052 | 311678 | 333270 | 311099 | 318682 | 12637 |
|  |  | 2 | 6718828 | 7159760 | 8651154 | 7509914 | 1012633 | 555220 | 578402 | 631064 | 588229 | 38865 |
|  |  | 3 | 5436854 | 5530708 | 6676394 | 5881319 | 690153 | 520942 | 573694 | 689309 | 594648 | 86117 |

|  |  |  | Ratio LucF/LucR | | | | |  |  | |  |  |
| --- | --- | --- | --- | --- | --- | --- | --- | --- | --- | --- | --- | --- |
|  |  | **Exp num** | **A** | **B** | **C** | **Mean** | **SD** | **Normalization** | **Mean** | **SD** | **Mann-Whitney p-value** | **Significance** |
| Normoxia | Gapmer  Ctrl | 1 | 0.1503 | 0.1424 | 0.1397 | 0.1441 | 0.005538 | 1.000 | 1.000 | 0.322 | *< 0.0001* | **** |
|  |  | 2 | 0.1392 | 0.1345 | 0.1311 | 0.1349 | 0.004059 | 1.000 |  |  |  |  |
|  |  | 3 | 0.1801 | 0.2839 | 0.2456 | 0.2365 | 0.05246 | 1.000 |  |  |  |  |
|  | Gapmer  Neat1 | 1 | 0.0891 | 0.0971 | 0.0925 | 0.0929 | 0.004031 | 0.644 | 0.551 | 0.076 |  |  |
|  |  | 2 | 0.0771 | 0.0803 | 0.089 | 0.0821 | 0.006182 | 0.609 |  |  |  |  |
|  |  | 3 | 0.1182 | 0.1018 | 0.1076 | 0.1092 | 0.008335 | 0.462 |  |  |  |  |
| Hypoxia | Gapmer  Ctrl | 1 | 0.1216 | 0.1285 | 0.1283 | 0.1262 | 0.003932 | 1.000 | 1.000 | 0.337 | *< 0.0001* | **** |
|  |  | 2 | 0.1655 | 0.1563 | 0.1641 | 0.1619 | 0.004972 | 1.000 |  |  |  |  |
|  |  | 3 | 0.1732 | 0.2868 | 0.2646 | 0.2415 | 0.060226 | 1.000 |  |  |  |  |
|  | Gapmer  Neat1 | 1 | 0.0817 | 0.0827 | 0.0736 | 0.0793 | 0.005032 | 0.629 | 0.489 | 0.066 |  |  |
|  |  | 2 | 0.0826 | 0.0808 | 0.0729 | 0.0788 | 0.005145 | 0.487 |  |  |  |  |
|  |  | 3 | 0.0958 | 0.1037 | 0.1032 | 0.1009 | 0.004435 | 0.418 |  |  |  |  |

|  |  |  | Renilla luciferase | | | | | Firefly luciferase | | | | |
| --- | --- | --- | --- | --- | --- | --- | --- | --- | --- | --- | --- | --- |
| **VEGFC IRES** |  | **Exp num** | **A** | **B** | **C** | **Mean** | **SD** | **A** | **B** | **C** | **Mean** | **SD** |
| Normoxia | Gapmer  Ctrl | 1 | 268948 | 215669 | 235986 | 240201 | 26889 | 20507 | 17652 | 22646 | 20268 | 2506 |
|  |  | 2 | 415401 | 314699 | 323438 | 351180 | 55789 | 77553 | 62353 | 68839 | 69581 | 7627 |
|  |  | 3 | 351021 | 1255659 | 996065 | 867582 | 465804 | 97373 | 242267 | 222849 | 187496 | 78651 |
|  | Gapmer  Neat1 | 1 | 744706 | 826236 | 741497 | 770813 | 48025 | 22755 | 23693 | 20765 | 22404 | 1495 |
|  |  | 2 | 1187605 | 1719953 | 1807553 | 1571704 | 335510 | 49411 | 71173 | 66705 | 62430 | 11494 |
|  |  | 3 | 1306031 | 1317445 | 1313896 | 1312458 | 5841 | 67750 | 61091 | 71268 | 66703 | 5169 |
| Hypoxia | Gapmer  Ctrl | 1 | 451023 | 464658 | 465074 | 460251 | 7995 | 23155 | 22050 | 24515 | 23240 | 1235 |
|  |  | 2 | 535899 | 538350 | 512109 | 528786 | 14494 | 94316 | 93879 | 102964 | 97053 | 5124 |
|  |  | 3 | 542368 | 596298 | 481042 | 539903 | 57668 | 149790 | 126538 | 153082 | 143137 | 14469 |
|  | Gapmer  Neat1 | 1 | 1414694 | 1558330 | 1365966 | 1446330 | 100008 | 30827 | 31648 | 24899 | 29125 | 3682 |
|  |  | 2 | 2542154 | 2180157 | 2613132 | 2445148 | 232217 | 111051 | 121182 | 118657 | 116964 | 5274 |
|  |  | 3 | 2405631 | 2088828 | 2353655 | 2282704 | 169902 | 107081 | 93526 | 94431 | 98346 | 7579 |

|  |  |  | Ratio LucF/LucR | | | | |  |  |  |  |  | |
| --- | --- | --- | --- | --- | --- | --- | --- | --- | --- | --- | --- | --- | --- |
|  |  | **Exp num** | **A** | **B** | **C** | **Mean** | **SD** | **Normalization** | **Mean** | **SD** | **Mann-Whitney p-value** | | **Significance** |
| Normoxia | Gapmer  Ctrl | 1 | 0.0762 | 0.0818 | 0.096 | 0.0847 | 0.01016 | 1.000 | 1.000 | 0.411 | *< 0.0001* | | **** |
|  |  | 2 | 0.1867 | 0.1981 | 0.2128 | 0.1992 | 0.013104 | 1.000 |  |  |  |  |  |
|  |  | 3 | 0.2774 | 0.1929 | 0.2237 | 0.2314 | 0.042743 | 1.000 |  |  |  |  |  |
|  | Gapmer  Neat1 | 1 | 0.0306 | 0.0287 | 0.028 | 0.0291 | 0.001323 | 0.343 | 0.233 | 0.057 |  |  |  |
|  |  | 2 | 0.0416 | 0.0414 | 0.0369 | 0.04 | 0.002652 | 0.201 |  |  |  |  |  |
|  |  | 3 | 0.0519 | 0.0464 | 0.0542 | 0.0508 | 0.004038 | 0.220 |  |  |  |  |  |
| Hypoxia | Gapmer  Ctrl | 1 | 0.0513 | 0.0475 | 0.0527 | 0.0505 | 0.002728 | 1.000 | 1.000 | 0.592 | *0.0003* | | *** |
|  |  | 2 | 0.176 | 0.1744 | 0.2011 | 0.1838 | 0.014958 | 1.000 |  |  |  |  |  |
|  |  | 3 | 0.2762 | 0.2122 | 0.3182 | 0.2689 | 0.053388 | 1.000 |  |  |  |  |  |
|  | Gapmer  Neat1 | 1 | 0.0218 | 0.0203 | 0.0182 | 0.0201 | 0.00179 | 0.398 | 0.222 | 0.080 |  |  |  |
|  |  | 2 | 0.0437 | 0.0556 | 0.0454 | 0.0482 | 0.006431 | 0.262 |  |  |  |  |  |
|  |  | 3 | 0.0445 | 0.0448 | 0.0401 | 0.0431 | 0.002614 | 0.160 |  |  |  |  |  |

|  | |  | Renilla luciferase | | | | | Firefly luciferase | | | | |
| --- | --- | --- | --- | --- | --- | --- | --- | --- | --- | --- | --- | --- |
| **VEGFD IRES** |  | **Exp num** | **A** | **B** | **C** | **Mean** | **SD** | **A** | **B** | **C** | **Mean** | **SD** |
| Normoxia | Gapmer  Ctrl | 1 | 186598 | 174458 | 194967 | 185341 | 10312 | 14000 | 12899 | 15863 | 14254 | 1498 |
|  |  | 2 | 236094 | 257184 | 307585 | 266954 | 36733 | 18079 | 16903 | 20655 | 18545 | 1919 |
|  |  | 3 | 238009 | 272627 | 201183 | 237273 | 35728 | 24834 | 23845 | 23129 | 23936 | 856 |
|  | Gapmer  Neat1 | 1 | 850116 | 863359 | 900946 | 871473 | 26369 | 53614 | 54124 | 46571 | 51436 | 4221 |
|  |  | 2 | 1699249 | 1517611 | 1361981 | 1526280 | 168801 | 83001 | 65774 | 61480 | 70085 | 11390 |
|  |  | 3 | 1227420 | 1258580 | 1355633 | 1280544 | 66869 | 67516 | 75980 | 85947 | 76481 | 9226 |
| Hypoxia | Gapmer  Ctrl | 1 | 320104 | 331836 | 371059 | 341000 | 26685 | 21527 | 20031 | 23687 | 21748 | 1838 |
|  |  | 2 | 370828 | 422876 | 450235 | 414646 | 40338 | 30107 | 32468 | 37560 | 33378 | 3809 |
|  |  | 3 | 550918 | 435861 | 281152 | 422644 | 135368 | 53330 | 39855 | 31894 | 41693 | 10836 |
|  | Gapmer  Neat1 | 1 | 1582942 | 1749642 | 1606739 | 1646441 | 90163 | 80420 | 99481 | 84808 | 88237 | 9982 |
|  |  | 2 | 2540713 | 2734360 | 3042934 | 2772669 | 253292 | 125626 | 136340 | 114288 | 125418 | 11027 |
|  |  | 3 | 2337486 | 1957657 | 1984333 | 2093158 | 212014 | 135295 | 129351 | 121929 | 128858 | 6697 |

|  |  |  | Ratio LucF/LucR | | | | |  |  |  |  |  | |
| --- | --- | --- | --- | --- | --- | --- | --- | --- | --- | --- | --- | --- | --- |
|  |  | **Exp num** | **A** | **B** | **C** | **Mean** | **SD** | **Normalization** | **Mean** | **SD** | **Mann-Whitney p-value** | | **Significance** |
| Normoxia | Gapmer  Ctrl | 1 | 0.075 | 0.0739 | 0.0814 | 0.0768 | 0.004009 | 1.000 | 1.000 | 0.202 | *< 0.0001* | | **** |
|  |  | 2 | 0.0766 | 0.0657 | 0.0672 | 0.0698 | 0.005896 | 1.000 |  |  |  |  |  |
|  |  | 3 | 0.1043 | 0.0875 | 0.115 | 0.1023 | 0.013868 | 1.000 |  |  |  |  |  |
|  | Gapmer  Neat1 | 1 | 0.0631 | 0.0627 | 0.0517 | 0.0591 | 0.006461 | 0.770 | 0.661 | 0.096 |  |  |  |
|  |  | 2 | 0.0488 | 0.0433 | 0.0451 | 0.0458 | 0.002807 | 0.656 |  |  |  |  |  |
|  |  | 3 | 0.055 | 0.0604 | 0.0634 | 0.0596 | 0.004251 | 0.583 |  |  |  |  |  |
| Hypoxia | Gapmer  Ctrl | 1 | 0.0672 | 0.0604 | 0.0638 | 0.0638 | 0.003442 | 1.000 | 1.000 | 0.210 | *0.0003* | | *** |
|  |  | 2 | 0.0812 | 0.0768 | 0.0834 | 0.0805 | 0.00338 | 1.000 |  |  |  |  |  |
|  |  | 3 | 0.0968 | 0.0914 | 0.1134 | 0.1006 | 0.011471 | 1.000 |  |  |  |  |  |
|  | Gapmer  Neat1 | 1 | 0.0508 | 0.0569 | 0.0528 | 0.0535 | 0.003087 | 0.838 | 0.657 | 0.101 |  |  |  |
|  |  | 2 | 0.0494 | 0.0499 | 0.0376 | 0.0456 | 0.006986 | 0.567 |  |  |  |  |  |
|  |  | 3 | 0.0579 | 0.0661 | 0.0614 | 0.0618 | 0.004108 | 0.615 |  |  |  |  |  |

|  |  |  | Renilla luciferase | | | | | Firefly luciferase | | | | |
| --- | --- | --- | --- | --- | --- | --- | --- | --- | --- | --- | --- | --- |
| **IGF1R IRES** |  | **Exp num** | **A** | **B** | **C** | **Mean** | **SD** | **A** | **B** | **C** | **Mean** | **SD** |
| Normoxia | Gapmer  Ctrl | 1 | 243181 | 214165 | 256708 | 238018 | 21736 | 64800 | 50375 | 74714 | 63296 | 12239 |
|  |  | 2 | 289104 | 236279 | 320432 | 281938 | 42532 | 57102 | 48588 | 64902 | 56864 | 8160 |
|  |  | 3 | 468014 | 341725 | 354953 | 388231 | 69410 | 89849 | 69873 | 68938 | 76220 | 11812 |
|  | Gapmer  Neat1 | 1 | 641540 | 541068 | 476621 | 553076 | 83113 | 90421 | 84962 | 64587 | 79990 | 13616 |
|  |  | 2 | 934259 | 1001588 | 982343 | 972730 | 34678 | 76744 | 86636 | 86743 | 83375 | 5742 |
|  |  | 3 | 909959 | 1202096 | 1057982 | 1056679 | 146073 | 87047 | 120227 | 113487 | 106920 | 17538 |
| Hypoxia | Gapmer  Ctrl | 1 | 150223 | 167451 | 189584 | 169086 | 19731 | 48001 | 55519 | 72037 | 58519 | 12295 |
|  |  | 2 | 567189 | 531510 | 569829 | 556176 | 21402 | 112568 | 108808 | 121830 | 114402 | 6702 |
|  |  | 3 | 213578 | 283108 | 594185 | 363623 | 202676 | 58103 | 88067 | 132981 | 93050 | 37687 |
|  | Gapmer  Neat1 | 1 | 422741 | 384291 | 361023 | 389351 | 31169 | 74275 | 73243 | 63546 | 70355 | 5919 |
|  |  | 2 | 1828828 | 1843042 | 1902069 | 1857980 | 38838 | 121169 | 137047 | 132974 | 130397 | 8247 |
|  |  | 3 | 1403043 | 1439067 | 800965 | 1214358 | 358462 | 148105 | 154836 | 97166 | 133369 | 31533 |

|  |  |  | Ratio LucF/LucR | | | | |  |  |  |  |  | |
| --- | --- | --- | --- | --- | --- | --- | --- | --- | --- | --- | --- | --- | --- |
|  |  | **Exp num** | **A** | **B** | **C** | **Mean** | **SD** | **Normalization** | **Mean** | **SD** | **Mann-Whitney p-value** | | **Significance** |
| Normoxia | Gapmer  Ctrl | 1 | 0.266 | 0.235 | 0.291 | 0.264 | 0.028 | 1.000 | 1.000 | 0.161 | *< 0.0001* | | **** |
|  |  | 2 | 0.198 | 0.206 | 0.203 | 0.202 | 0.004 | 1.000 |  |  |  |  |  |
|  |  | 3 | 0.192 | 0.204 | 0.194 | 0.197 | 0.007 | 1.000 |  |  |  |  |  |
|  | Gapmer  Neat1 | 1 | 0.141 | 0.157 | 0.136 | 0.144 | 0.011 | 0.547 | 0.499 | 0.123 |  |  |  |
|  |  | 2 | 0.082 | 0.086 | 0.088 | 0.086 | 0.003 | 0.424 |  |  |  |  |  |
|  |  | 3 | 0.096 | 0.100 | 0.107 | 0.101 | 0.006 | 0.513 |  |  |  |  |  |
| Hypoxia | Gapmer  Ctrl | 1 | 0.320 | 0.332 | 0.380 | 0.344 | 0.032 | 1.000 | 1.000 | 0.241 | *< 0.0001* | | **** |
|  |  | 2 | 0.198 | 0.205 | 0.214 | 0.206 | 0.008 | 1.000 |  |  |  |  |  |
|  |  | 3 | 0.272 | 0.311 | 0.224 | 0.269 | 0.044 | 1.000 |  |  |  |  |  |
|  | Gapmer  Neat1 | 1 | 0.176 | 0.191 | 0.176 | 0.181 | 0.009 | 0.526 | 0.443 | 0.179 |  |  |  |
|  |  | 2 | 0.066 | 0.074 | 0.070 | 0.070 | 0.004 | 0.341 |  |  |  |  |  |
|  |  | 3 | 0.106 | 0.108 | 0.121 | 0.111 | 0.009 | 0.414 |  |  |  |  |  |

|  |  |  |  | Renilla luciferase | | | | | Firefly luciferase | | | | |
| --- | --- | --- | --- | --- | --- | --- | --- | --- | --- | --- | --- | --- | --- |
| **c-myc IRES** |  | **Exp num** | | **A** | **B** | **C** | **Mean** | **SD** | **A** | **B** | **C** | **Mean** | **SD** |
| Normoxia | Gapmer  Ctrl | 1 | | 409115.667 | 409858.667 | 477078.667 | 432017.667 | 39026 | 56600 | 57218 | 70155 | 61324.4444 | 7654 |
|  |  | 2 | | 537995.333 | 865640 | 785543.667 | 729726.333 | 170805 | 129074 | 188805 | 190860 | 169579.778 | 35094 |
|  |  | 3 | | 817358.417 | 884814.75 | 692293.083 | 798155.417 | 97687 | 386170 | 358606 | 369298 | 371358 | 13897 |
|  | Gapmer  Neat1 | 1 | | 815950.667 | 1782212.67 | 1451628 | 1349930.44 | 491093 | 107182 | 141253 | 114757 | 121064 | 17889 |
|  |  | 2 | | 4228764.33 | 3975787.67 | 3648284.33 | 3950945.44 | 291036 | 312037 | 298030 | 266821 | 292296 | 23147 |
|  |  | 3 | | 3337134.42 | 3277986.42 | 3093559.08 | 3236226.64 | 127044 | 467337 | 469577 | 434836 | 457250 | 19444 |
| Hypoxia | Gapmer  Ctrl | 1 | | 848388.333 | 733310.667 | 820763 | 800820.667 | 60075 | 87044 | 81354 | 91270 | 86555.8889 | 4976 |
|  |  | 2 | | 1022245 | 1179393.67 | 1261896 | 1154511.56 | 121748 | 256743 | 327423 | 339370 | 307845.444 | 44658 |
|  |  | 3 | | 1261477.5 | 931395.5 | 910791.833 | 1034554.94 | 196791 | 676240 | 500283 | 605173 | 593899 | 88519 |
|  | Gapmer  Neat1 | 1 | | 3163184.67 | 3416411.67 | 2949987.67 | 3176528 | 233498 | 214005 | 209213 | 176425 | 199881.111 | 20454 |
|  |  | 2 | | 6882853 | 6708709.33 | 6411856.67 | 6667806.33 | 238147 | 562750 | 553645 | 461767 | 526054.111 | 55860 |
|  |  | 3 | | 4854553.83 | 4079910.83 | 4678820.17 | 4537761.61 | 406129 | 834137 | 638674 | 715474 | 729428 | 98476 |

|  |  | |  | | Ratio LucF/LucR | | | | |  |  |  |  |  | | |
| --- | --- | --- | --- | --- | --- | --- | --- | --- | --- | --- | --- | --- | --- | --- | --- | --- |
|  |  |  | **Exp num** | | **A** | **B** | **C** | **Mean** | **SD** | **Normalization** | **Mean** | **SD** | **Mann-Whitney p-value** | | **Significance** | |
| Normoxia | Gapmer  Ctrl | | 1 | | 0.1383 | 0.1396 | 0.1471 | 0.1417 | 0.004705 | 1.000 | 1.000 | 0.534 | *0.0009* | | *** | |
|  |  |  | 2 | | 0.2399 | 0.2181 | 0.243 | 0.2337 | 0.013556 | 1.000 |  |  |  |  |  |  |
|  |  |  | 3 | | 0.4725 | 0.4053 | 0.5334 | 0.4704 | 0.064101 | 1.000 |  |  |  |  |  |  |
|  | Gapmer  Neat1 | | 1 | | 0.1314 | 0.0793 | 0.0791 | 0.0966 | 0.03014 | 0.682 | 0.369 | 0.118 |  |  |  |  |
|  |  |  | 2 | | 0.0738 | 0.075 | 0.0731 | 0.074 | 0.000925 | 0.317 |  |  |  |  |  |  |
|  |  |  | 3 | | 0.14 | 0.1433 | 0.1406 | 0.1413 | 0.001723 | 0.300 |  |  |  |  |  |  |
| Hypoxia | Gapmer  Ctrl | | 1 | | 0.1026 | 0.1109 | 0.1112 | 0.1082 | 0.004893 | 1.000 | 1.000 | 0.664 | *0.0036* | | ** | |
|  |  |  | 2 | | 0.2512 | 0.2776 | 0.2689 | 0.2659 | 0.01349 | 1.000 |  |  |  |  |  |  |
|  |  |  | 3 | | 0.5361 | 0.5371 | 0.6644 | 0.5792 | 0.073814 | 1.000 |  |  |  |  |  |  |
|  | Gapmer  Neat1 | | 1 | | 0.0677 | 0.0612 | 0.0598 | 0.0629 | 0.00418 | 0.581 | 0.317 | 0.144 |  |  |  |  |
|  |  |  | 2 | | 0.0818 | 0.0825 | 0.072 | 0.0788 | 0.005859 | 0.296 |  |  |  |  |  |  |
|  |  |  | 3 | | 0.1718 | 0.1565 | 0.1529 | 0.1604 | 0.010035 | 0.277 |  |  |  |  |  |  |
|  | |  | |  | | Renilla luciferase | | | | | Firefly luciferase | | | | | |
| **EMCV IRES** | |  | | **Exp num** | | **A** | **B** | **C** | **Mean** | **SD** | **A** | **B** | **C** | **Mean** | | **SD** |
| Normoxia | | Gapmer  Ctrl | | 1 | | 231360 | 200119 | 195599 | 209026 | 19473 | 184926 | 161799 | 151222 | 165982 | | 17237 |
|  |  |  |  | 2 | | 1236254 | 1034251 | 1190294 | 1153600 | 105883 | 802079 | 639691 | 733674 | 725148 | | 81529 |
|  |  |  |  | 3 | | 1605131 | 1009872 | 1248146 | 1287716 | 299596 | 978404 | 644626 | 759438 | 794156 | | 169576 |
|  |  | Gapmer  Neat1 | | 1 | | 262296 | 274310 | 227852 | 254819 | 24115 | 212785 | 205023 | 182198 | 200002 | | 15900 |
|  |  |  |  | 2 | | 4541077 | 4722169 | 3821501 | 4361582 | 476408 | 2388347 | 2531111 | 2288426 | 2402628 | | 121971 |
|  |  |  |  | 3 | | 3736975 | 4412734 | 4672802 | 4274170 | 483056 | 2141250 | 2525467 | 2756413 | 2474377 | | 310748 |
| Hypoxia | | Gapmer  Ctrl | | 1 | | 139570 | 102655 | 101666 | 114630 | 21604 | 134824 | 88039 | 89602 | 104155 | | 26571 |
|  |  |  |  | 2 | | 1557383 | 1688310 | 1647310 | 1631001 | 66969 | 1176925 | 1138768 | 1159710 | 1158467 | | 19109 |
|  |  |  |  | 3 | | 1469437 | 1513608 | 1782542 | 1588529 | 169465 | 1497706 | 1194433 | 1533246 | 1408462 | | 186204 |
|  |  | Gapmer  Neat1 | | 1 | | 159440 | 130826 | 155274 | 148513 | 15459 | 154391 | 116021 | 107166 | 125859 | | 25103 |
|  |  |  |  | 2 | | 9243748 | 9423188 | 9243336 | 9303424 | 103719 | 4831155 | 5317539 | 5263458 | 5137384 | | 266577 |
|  |  |  |  | 3 | | 4821342 | 4654942 | 3747799 | 4408028 | 577796 | 3512787 | 3241094 | 3165797 | 3306559 | | 182523 |

|  |  |  | Ratio LucF/LucR | | | | |  |  |  |  |  | |
| --- | --- | --- | --- | --- | --- | --- | --- | --- | --- | --- | --- | --- | --- |
|  |  | **Exp num** | **A** | **B** | **C** | **Mean** | **SD** | **Normalization** | **Mean** | **SD** | **Mann-Whitney p-value** | | **Significance** |
| Normoxia | Gapmer  Ctrl | 1 | 0.799 | 0.809 | 0.773 | 0.794 | 0.018 | 1.000 | 1.000 | 0.127 | *0.1348* | | ns |
|  |  | 2 | 0.649 | 0.619 | 0.616 | 0.628 | 0.018 | 1.000 |  |  |  |  |  |
|  |  | 3 | 0.610 | 0.638 | 0.608 | 0.619 | 0.017 | 1.000 |  |  |  |  |  |
|  | Gapmer  Neat1 | 1 | 0.811 | 0.747 | 0.800 | 0.786 | 0.034 | 0.990 | 0.940 | 0.167 |  |  |  |
|  |  | 2 | 0.526 | 0.536 | 0.599 | 0.554 | 0.039 | 0.882 |  |  |  |  |  |
|  |  | 3 | 0.573 | 0.572 | 0.590 | 0.578 | 0.010 | 0.935 |  |  |  |  |  |
| Hypoxia | Gapmer  Ctrl | 1 | 0.966 | 0.858 | 0.881 | 0.902 | 0.057 | 1.000 | 1.000 | 0.138 | *0.1348* | | ns |
|  |  | 2 | 0.756 | 0.675 | 0.704 | 0.711 | 0.041 | 1.000 |  |  |  |  |  |
|  |  | 3 | 1.019 | 0.789 | 0.860 | 0.890 | 0.118 | 1.000 |  |  |  |  |  |
|  | Gapmer  Neat1 | 1 | 0.968 | 0.887 | 0.690 | 0.848 | 0.143 | 0.941 | 0.862 | 0.186 |  |  |  |
|  |  | 2 | 0.523 | 0.564 | 0.569 | 0.552 | 0.026 | 0.776 |  |  |  |  |  |
|  |  | 3 | 0.729 | 0.696 | 0.845 | 0.757 | 0.078 | 0.850 |  |  |  |  |  |

**Supplementary file 3. IRES activities in normoxic and hypoxic HL-1 cells after Neat1 knock-down.**

HL-1 cells were transduced with Lucky Luke bicistronic lentivectors containing the IRES of FGF1, FGF2, VEGFA (a or b), VEGFC, VEGFD, IGF1R, c-myc or EMCV. Cells were then treated with a pool of gapmers Neat1, or control during normoxia or hypoxia 1% O_2._ For the lentivector with FGF1 IRES, cells were also treated with the gapmer Neat1_2 (N1_2, page 2). Renilla and firefly luciferase activities were measured (upper panel) and the IRES activities evaluated with the ratio LucF/LucR (lower panel). Mann-Whitney test was performed with n=9. *p<0.05, **p<0.01, ***<0.001, ****p<0.0001. For each IRES the mean has been calculated with nine cell culture biological replicates (Experiments A, B, C correspond to experiments performed at different dates, while 1, 2, 3 are experiments performed in parallel at the same date, each of them being already the mean of three technical replicates (27 technical replicates in total).
