## Supplementary files 1-8 for "Long non-coding RNA Neat1 and paraspeckle components are translational regulators in hypoxia": V2 Supplementary file 4 SFPQ rps2 hM Ncl KD.docx

|  |  |  | Renilla luciferase | | | | | Firefly luciferase | | | | |
| --- | --- | --- | --- | --- | --- | --- | --- | --- | --- | --- | --- | --- |
| **A/Sfpq knock-down**  **(FGF1 IRES)** |  | **Exp num** | **A** | **B** | **C** | **Mean** | **SD** | **A** | **B** | **C** | **Mean** | **SD** |
| Normoxia | siCtrl | 1 | 2673188 | 3784317 | 3075954 | 3177819 | 459300 | 455288 | 826990 | 612860 | 631712 | 152331 |
|  |  | 2 | 4321498 | 6005077 | 5319145 | 5215240 | 691234 | 908764 | 1480053 | 1477175 | 1288664 | 268633 |
|  |  | 3 | 1955203 | 1917089 | 2081085 | 1984459 | 70074 | 338768 | 304406 | 369720 | 337631 | 26676 |
|  | siSfpq | 1 | 4277464 | 4832185 | 4054960 | 4388203 | 326820 | 1062037 | 1257385 | 1057070 | 1125497 | 93281 |
|  |  | 2 | 4686601 | 6609368 | 6721005 | 6005658 | 933827 | 1029120 | 1623127 | 1821880 | 1491375 | 336785 |
|  |  | 3 | 2985174 | 2493406 | 2992549 | 2823709 | 233579 | 468485 | 388556 | 538616 | 465219 | 61305 |
| Hypoxia | siCtrl | 1 | 4241597 | 3454129 | 3728290 | 3808005 | 326387 | 761014 | 716463 | 654339 | 710605 | 43747 |
|  |  | 2 | 6004439 | 5725646 | 6186192 | 5972092 | 189403 | 1768671 | 1788136 | 1744898 | 1767235 | 17681 |
|  |  | 3 | 2151385 | 2008858 | 1959317 | 2039853 | 81417 | 322854 | 298111 | 329513 | 316826 | 13510 |
|  | siSfpq | 1 | 4901651 | 4468562 | 5339205 | 4903139 | 355440 | 1219052 | 1149188 | 1346529 | 1238256 | 81701 |
|  |  | 2 | 7044581 | 6792348 | 6677279 | 6838069 | 153396 | 2304701 | 1968534 | 2252785 | 2175340 | 147762 |
|  |  | 3 | 3350779 | 3020378 | 2829277 | 3066811 | 215419 | 481217 | 394264 | 435170 | 436883 | 35519 |

|  |  |  | Ratio LucF/LucR | | | | |  |  |  |  |  |
| --- | --- | --- | --- | --- | --- | --- | --- | --- | --- | --- | --- | --- |
|  |  | **Exp num** | **A** | **B** | **C** | **Mean** | **SD** | **Normalisation** | **Mean** | **SD** | **Mann-Whitney p-value** | **Significance** |
| Normoxia | siCtrl | 1 | 0.170 | 0.219 | 0.199 | 0.194 | 0.020 | 1.000 | 1.000 | 0.193 | *0.3993* | ns |
|  |  | 2 | 0.210 | 0.246 | 0.278 | 0.245 | 0.034 | 1.000 |  |  |  |  |
|  |  | 3 | 0.173 | 0.159 | 0.178 | 0.170 | 0.010 | 1.000 |  |  |  |  |
|  | siSfpq | 1 | 0.248 | 0.260 | 0.261 | 0.256 | 0.006 | 1.319 | 1.091 | 0.226 |  |  |
|  |  | 2 | 0.220 | 0.246 | 0.271 | 0.245 | 0.026 | 1.002 |  |  |  |  |
|  |  | 3 | 0.157 | 0.156 | 0.180 | 0.164 | 0.014 | 0.967 |  |  |  |  |
| Hypoxia | siCtrl | 1 | 0.179 | 0.207 | 0.176 | 0.193 | 0.020 | 1.000 | 1.000 | 0.306 | *0.7137* | ns |
|  |  | 2 | 0.295 | 0.312 | 0.282 | 0.296 | 0.015 | 1.000 |  |  |  |  |
|  |  | 3 | 0.150 | 0.148 | 0.168 | 0.156 | 0.011 | 1.000 |  |  |  |  |
|  | siSfpq | 1 | 0.249 | 0.257 | 0.252 | 0.253 | 0.003 | 1.306 | 1.116 | 0.366 |  |  |
|  |  | 2 | 0.327 | 0.290 | 0.337 | 0.318 | 0.025 | 1.074 |  |  |  |  |
|  |  | 3 | 0.144 | 0.131 | 0.154 | 0.143 | 0.012 | 0.917 |  |  |  |  |

|  |  |  | Renilla luciferase | | | | | Firefly luciferase | | | | |
| --- | --- | --- | --- | --- | --- | --- | --- | --- | --- | --- | --- | --- |
| **B/Rps2 knock-down**  **(FGF1 IRES)** |  | **Exp num** | **A** | **B** | **C** | **Mean** | **SD** | **A** | **B** | **C** | **Mean** | **SD** |
| Normoxia | siCtrl | 1 | 70175 | 101667 | 79068 | 83637 | 16236 | 32496 | 38405 | 28998 | 33300 | 4755 |
|  |  | 2 | 480646 | 282659 | 449570 | 404292 | 106477 | 150414.583 | 102400.583 | 132897.917 | 128571.028 | 24297.6856 |
|  |  | 3 | 352222 | 388895 | 368737 | 369951 | 18367 | 126831 | 120091 | 104519 | 117147 | 11444 |
|  | siRps2 | 1 | 86346 | 72676 | 72110 | 77044 | 8061 | 29502 | 25309 | 23922 | 26245 | 2905 |
|  |  | 2 | 372877 | 248168 | 482623 | 367889 | 117307 | 106876.917 | 75175.5833 | 122715.583 | 101589.361 | 24207.0565 |
|  |  | 3 | 315917 | 304772 | 236089 | 285593 | 43232 | 77833 | 76598 | 53539 | 69323 | 13684 |
| Hypoxia | siCtrl | 1 | 108205 | 109413 | 118115 | 111911 | 5406 | 50653 | 46424 | 44762 | 47280 | 3037 |
|  |  | 2 | 1200321.13 | 675913.125 | 1069716.79 | 981983.681 | 272990.406 | 301970.917 | 212601.75 | 252865.917 | 255812.861 | 44757.4055 |
|  |  | 3 | 619170 | 654180 | 613280 | 628877 | 22110 | 191284 | 180584 | 172469 | 181446 | 9437 |
|  | siRps2 | 1 | 116363 | 104800 | 128883 | 116682 | 12045 | 46670 | 35470 | 40340 | 40826 | 5616 |
|  |  | 2 | 919414.625 | 544829.792 | 1039654.46 | 834632.958 | 258077.141 | 226374.25 | 182160.583 | 212224.917 | 206919.917 | 22579.1795 |
|  |  | 3 | 531687 | 477308 | 541537 | 516844 | 34592 | 145562 | 107598 | 114449 | 122537 | 20233 |

|  |  |  | Ratio LucF/LucR | | | | |  |  |  |  |  |
| --- | --- | --- | --- | --- | --- | --- | --- | --- | --- | --- | --- | --- |
|  |  | **Exp num** | **A** | **B** | **C** | **Mean** | **SD** | **Normalisation** | **Mean** | **SD** | **Mann-Whitney p-value** | **Significance** |
| Normoxia | siCtrl | 1 | 0.46307601 | 0.37775121 | 0.36674288 | 0.40252337 | 0.0527282 | 1.000 | 1.000 | 0.159 | 0.0244 | * |
|  |  | 2 | 0.31294268 | 0.36227588 | 0.29561109 | 0.32360989 | 0.03458888 | 1.000 |  |  |  |  |
|  |  | 3 | 0.36008992 | 0.30879981 | 0.28345172 | 0.31744715 | 0.03904403 | 1.000 |  |  |  |  |
|  | siRps2 | 1 | 0.3416753 | 0.34824991 | 0.33174882 | 0.34055801 | 0.00830709 | 0.846 | 0.827 | 0.131 |  |  |
|  |  | 2 | 0.28662797 | 0.30292199 | 0.25426793 | 0.28127263 | 0.02476518 | 0.869 |  |  |  |  |
|  |  | 3 | 0.24637236 | 0.25132931 | 0.2267749 | 0.24149219 | 0.01298429 | 0.761 |  |  |  |  |
| Hypoxia | siCtrl | 1 | 0.4681233 | 0.42429711 | 0.37897318 | 0.42379786 | 0.04457715 | 1.000 | 1.000 | 0.245 | 0.2547 | ns |
|  |  | 2 | 0.25157511 | 0.31454005 | 0.23638585 | 0.26750034 | 0.04143947 | 1.000 |  |  |  |  |
|  |  | 3 | 0.30893655 | 0.27604721 | 0.28122452 | 0.28873609 | 0.01768459 | 1.000 |  |  |  |  |
|  | siRps2 | 1 | 0.40106926 | 0.33845174 | 0.31299501 | 0.35083867 | 0.04532489 | 0.828 | 0.867 | 0.207 |  |  |
|  |  | 2 | 0.24621563 | 0.33434402 | 0.20413024 | 0.2615633 | 0.06644976 | 0.978 |  |  |  |  |
|  |  | 3 | 0.27377428 | 0.22542705 | 0.21134192 | 0.23684775 | 0.0327456 | 0.820 |  |  |  |  |

|  |  |  | Renilla luciferase | | | | | Firefly luciferase | | | | |
| --- | --- | --- | --- | --- | --- | --- | --- | --- | --- | --- | --- | --- |
| **C/hnRNPM knock-down**  **(FGF1 IRES)** |  | **Exp num** | **A** | **B** | **C** | **Mean** | **SD** | **A** | **B** | **C** | **Mean** | **SD** |
| Normoxia | siCtrl | 1 | 70175 | 101667 | 79068 | 83637 | 16236 | 32496 | 38405 | 28998 | 33300 | 4755 |
|  |  | 2 | 480646 | 282659 | 449570 | 404292 | 106477 | 150414.583 | 102400.583 | 132897.917 | 128571.028 | 24297.6856 |
|  |  | 3 | 352222 | 388895 | 368737 | 369951 | 18367 | 126831 | 120091 | 104519 | 117147 | 11444 |
|  | sihnRNPM | 1 | 73070 | 86184 | 94096 | 84450 | 10620 | 31149 | 31919 | 39363 | 34144 | 4536 |
|  |  | 2 | 463775.458 | 444702.125 | 669255.458 | 525911.014 | 124505.704 | 160032.917 | 111057.583 | 169284.917 | 146791.806 | 31290.5873 |
|  |  | 3 | 435784 | 383796 | 326717 | 382099 | 54553 | 116837 | 91766 | 91569 | 100057 | 14532 |
| Hypoxia | siCtrl | 1 | 108205 | 109413 | 118115 | 111911 | 5406 | 50653 | 46424 | 44762 | 47280 | 3037 |
|  |  | 2 | 1200321.13 | 675913.125 | 1069716.79 | 981983.681 | 272990.406 | 301970.917 | 212601.75 | 252865.917 | 255812.861 | 44757.4055 |
|  |  | 3 | 619170 | 654180 | 613280 | 628877 | 22110 | 191284 | 180584 | 172469 | 181446 | 9437 |
|  | sihnRNPM | 1 | 149997 | 102166 | 111967 | 121377 | 25266 | 65583 | 48006 | 55825 | 56471 | 8806 |
|  |  | 2 | 736219.125 | 989554.458 | 989554.458 | 905109.347 | 146263.223 | 329601.25 | 237066.917 | 313103.583 | 293257.25 | 49356.4426 |
|  |  | 3 | 727443 | 691434 | 691434 | 703437 | 20790 | 197670 | 177901 | 152871 | 176147 | 22451 |

|  |  |  | Ratio LucF/LucR | | | | |  |  |  |  |  |
| --- | --- | --- | --- | --- | --- | --- | --- | --- | --- | --- | --- | --- |
|  |  | **Exp num** | **A** | **B** | **C** | **Mean** | **SD** | **Normalisation** | **Mean** | **SD** | **Mann-Whitney p-value** | **Significance** |
| Normoxia | siCtrl | 1 | 0.46307601 | 0.37775121 | 0.36674288 | 0.40252337 | 0.0527282 | 1.000 | 1.000 | 0.159 | 0.2547 | ns |
|  |  | 2 | 0.31294268 | 0.36227588 | 0.29561109 | 0.32360989 | 0.03458888 | 1.000 |  |  |  |  |
|  |  | 3 | 0.36008992 | 0.30879981 | 0.28345172 | 0.31744715 | 0.03904403 | 1.000 |  |  |  |  |
|  | sihnRNPM | 1 | 0.42629377 | 0.37035442 | 0.41832908 | 0.40499242 | 0.03026058 | 1.006 | 0.910 | 0.214 |  |  |
|  |  | 2 | 0.34506551 | 0.24973477 | 0.25294514 | 0.28258181 | 0.05413628 | 0.873 |  |  |  |  |
|  |  | 3 | 0.2681077 | 0.23910067 | 0.28027021 | 0.26249286 | 0.0211513 | 0.827 |  |  |  |  |
| Hypoxia | siCtrl | 1 | 0.4681233 | 0.42429711 | 0.37897318 | 0.42379786 | 0.04457715 | 1.000 | 1.000 | 0.245 | 0.7137 | ns |
|  |  | 2 | 0.25157511 | 0.31454005 | 0.23638585 | 0.26750034 | 0.04143947 | 1.000 |  |  |  |  |
|  |  | 3 | 0.30893655 | 0.27604721 | 0.28122452 | 0.28873609 | 0.01768459 | 1.000 |  |  |  |  |
|  | sihnRNPM | 1 | 0.43722842 | 0.46988011 | 0.49858391 | 0.46856415 | 0.03069891 | 1.106 | 1.075 | 0.339 |  |  |
|  |  | 2 | 0.4476945 | 0.23956935 | 0.31640864 | 0.3345575 | 0.10524284 | 1.251 |  |  |  |  |
|  |  | 3 | 0.27173224 | 0.25729349 | 0.22109288 | 0.25003954 | 0.02608737 | 0.866 |  |  |  |  |

|  |  |  | Renilla luciferase | | | | | Firefly luciferase | | | | |
| --- | --- | --- | --- | --- | --- | --- | --- | --- | --- | --- | --- | --- |
| **D/Nucleolin knock-down**  **(FGF1 IRES)** |  | **Exp num** | **A** | **B** | **C** | **Mean** | **SD** | **A** | **B** | **C** | **Mean** | **SD** |
| Normoxia | siCtrl | 1 | 70175 | 101667 | 79068 | 83637 | 16236 | 32496 | 38405 | 28998 | 33300 | 4755 |
|  |  | 2 | 480646 | 282659 | 449570 | 404292 | 106477 | 150414.583 | 102400.583 | 132897.917 | 128571.028 | 24297.6856 |
|  |  | 3 | 352222 | 388895 | 368737 | 369951 | 18367 | 126831 | 120091 | 104519 | 117147 | 11444 |
|  | siNcl | 1 | 140 655 | 93 009 | 113 335 | 115 666 | 23909 | 36314 | 36674 | 31297 | 34762 | 3006 |
|  |  | 2 | 689994.458 | 996580.458 | 930164.792 | 872246.569 | 161290.527 | 159799.25 | 180704.25 | 200234.917 | 180246.139 | 20221.7255 |
|  |  | 3 | 528 673 | 559 187 | 454 446 | 514102 | 53869 | 136788 | 112437 | 111030 | 120085 | 14483 |
| Hypoxia | siCtrl | 1 | 108205 | 109413 | 118115 | 111911 | 5406 | 50653 | 46424 | 44762 | 47280 | 3037 |
|  |  | 2 | 1200321.13 | 675913.125 | 1069716.79 | 981983.681 | 272990.406 | 301970.917 | 212601.75 | 252865.917 | 255812.861 | 44757.4055 |
|  |  | 3 | 619170 | 654180 | 613280 | 628877 | 22110 | 191284 | 180584 | 172469 | 181446 | 9437 |
|  | siNcl | 1 | 148 450 | 157 125 | 158 518 | 154 698 | 5455 | 56578 | 58848 | 52861 | 56096 | 3023 |
|  |  | 2 |  | 1894315.79 | 2234800.46 | 2064558.13 | 240759.017 |  | 319677.917 | 412429.25 | 366053.583 | 65585.0968 |
|  |  | 3 | 825 673 | 826 588 | 771 221 | 807827 | 31706 | 212040 | 205754 | 208187 | 208660 | 3169 |

|  |  |  | Ratio LucF/LucR | | | | |  |  |  |  |  |
| --- | --- | --- | --- | --- | --- | --- | --- | --- | --- | --- | --- | --- |
|  |  | **Exp num** | **A** | **B** | **C** | **Mean** | **SD** | **Normalisation** | **Mean** | **SD** | **Mann-Whitney p-value** | **Significance** |
| Normoxia | siCtrl | 1 | 0.46307601 | 0.37775121 | 0.36674288 | 0.40252337 | 0.0527282 | 1.000 | 1.000 | 0.159 | 0.0028 | ** |
|  |  | 2 | 0.31294268 | 0.36227588 | 0.29561109 | 0.32360989 | 0.03458888 | 1.000 |  |  |  |  |
|  |  | 3 | 0.36008992 | 0.30879981 | 0.28345172 | 0.31744715 | 0.03904403 | 1.000 |  |  |  |  |
|  | siNcl | 1 | 0.25817998 | 0.39430687 | 0.27614486 | 0.3095439 | 0.07395441 | 0.769 | 0.722 | 0.177 |  |  |
|  |  | 2 | 0.23159498 | 0.1813243 | 0.21526822 | 0.20939583 | 0.02564467 | 0.647 |  |  |  |  |
|  |  | 3 | 0.25873928 | 0.20107203 | 0.2443192 | 0.23471017 | 0.03001046 | 0.739 |  |  |  |  |
| Hypoxia | siCtrl | 1 | 0.4681233 | 0.42429711 | 0.37897318 | 0.42379786 | 0.04457715 | 1.000 | 1.000 | 0.245 | 0.2716 | ns |
|  |  | 2 | 0.25157511 | 0.31454005 | 0.23638585 | 0.26750034 | 0.04143947 | 1.000 |  |  |  |  |
|  |  | 3 | 0.30893655 | 0.27604721 | 0.28122452 | 0.28873609 | 0.01768459 | 1.000 |  |  |  |  |
|  | siNcl | 1 | 0.38112243 | 0.37453169 | 0.33346768 | 0.3630406 | 0.02582204 | 0.857 | 0.849 | 0.246 |  |  |
|  |  | 2 |  | 0.1687564 | 0.18454858 | 0.17665249 | 0.01116676 | 0.660 |  |  |  |  |
|  |  | 3 | 0.25680835 | 0.24892012 | 0.26994497 | 0.25855781 | 0.01062104 | 0.895 |  |  |  |  |

**Supplementary file 4. FGF1 IRES activity in normoxic and hypoxic HL-1 cells after Sfpq, Rps2, hnRNPM knock-down.**

HL-1 cells were transduced with a Lucky Luke bicistronic lentivector containing the IRES of FGF1. Cells were then treated with siSfpq (A), siRps2 (B), sihnRNPM (C), siNucleolin (siNcl, D) or siControl (siCtrl) smartpools during normoxia or hypoxia 1% O_2._ Renilla and firefly luciferase activities were measured and the IRES activities evaluated with the ratio LucF/LucR. Mann-Whitney test was performed with n=9 (n=12 for FGF1 IRES). *p<0.05, **p<0.01, ***<0.001, ****p<0.0001. For each siRNA the mean has been calculated with nine cell culture biological replicates (Experiments A, B, C correspond to experiments performed at different dates, while 1, 2, 3 are experiments performed in parallel at the same date, each of them being already the mean of three technical replicates (27 technical replicates in total).
