## Supplementary files 1-8 for "Long non-coding RNA Neat1 and paraspeckle components are translational regulators in hypoxia": V2 Supplementary file 5 P54nrb KD.docx

|  |  |  | Renilla luciferase | | | | | Firefly luciferase | | | | |
| --- | --- | --- | --- | --- | --- | --- | --- | --- | --- | --- | --- | --- |
| **FGF1** |  | **Exp num** | **A** | **B** | **C** | **Mean** | **SD** | **A** | **B** | **C** | **Mean** | **SD** |
| normoxia | siCtrl | 1 | 15614583 | 13834477 | 14212422 | 14553828 | 937877 | 4112521 | 3337660 | 3785300 | 3745160 | 388987 |
|  |  | 2 | 6211969 | 5719040 | 5980393 | 5970467 | 246615 | 1921913 | 1714365 | 1979669 | 1871982 | 139522 |
|  |  | 3 | 1590009 | 1381692 | 1381001 | 1450901 | 120472 | 251834 | 194834 | 238849 | 228506 | 29875 |
|  |  | 4 | 79768 | 104037 | 89183 | 90996 | 9990 | 10219 | 19335 | 13189 | 14248 | 3796 |
|  | siP54nrb | 1 | 22178973 | 20147753 | 24411755 | 22246160 | 2132795 | 3686041 | 3111936 | 4004601 | 3600859 | 452388 |
|  |  | 2 | 2829392 | 2480111 | 3381753 | 2897086 | 454617 | 477464 | 426225 | 582440 | 495376 | 79633 |
|  |  | 3 | 1431898 | 2111776 | 1687224 | 1743633 | 343431 | 188751 | 304274 | 257561 | 250196 | 58112 |
|  |  | 4 | 102804 | 151024 | 134259 | 129362 | 24480 | 8357 | 20008 | 15857 | 14741 | 5905 |
| Hypoxia | siCtrl | 1 | 12867685 | 12005389 | 13333771 | 12735615 | 673967 | 4412100 | 4267292 | 4713037 | 4464143 | 227384 |
|  |  | 2 | 6348250 | 5853231 | 5412593 | 5871358 | 468092 | 2064972 | 1756583 | 1711633 | 1844396 | 192342 |
|  |  | 3 | 1121156 | 1093631 | 1022224 | 1079004 | 51062 | 287035 | 264567 | 162669 | 238090 | 66276 |
|  |  | 4 | 138993 | 155606 | 162284 | 152294 | 9792 | 14170 | 24385 | 20632 | 19729 | 4219 |
|  | siP54nrb | 1 | 20105090 | 25274081 | 24284875 | 23221349 | 2743708 | 3842071 | 5123829 | 5006211 | 4657370 | 708515 |
|  |  | 2 | 6618266 | 5251099 | 5113389 | 5660918 | 831942 | 1054684 | 761848 | 686201 | 834244 | 194617 |
|  |  | 3 | 1197812 | 1245065 | 1131357 | 1191411 | 57123 | 198596 | 323032 | 239965 | 253864 | 63372 |
|  |  | 4 | 227563 | 130725 | 169940 | 176076 | 39772 | 24628 | 15987 | 22092 | 20902 | 3626 |

|  |  |  | Ratio LucF/LucR | | | | |  |  | |  | |  | |  |  |
| --- | --- | --- | --- | --- | --- | --- | --- | --- | --- | --- | --- | --- | --- | --- | --- | --- |
| **FGF1 IRES** |  | **Exp num** | **A** | **B** | **C** | **Mean** | **SD** | **Normalisation** | | **Mean** | | **SD** | | **Mann-Whitney p-value** | | **Significance** |
| normoxia | siCtrl | 1 | 0.263 | 0.241 | 0.266 | 0.257 | 0.014 | 1.000 | | 1.000 | | 0.330 | | *0.0138* | | * |
|  |  | 2 | 0.309 | 0.300 | 0.331 | 0.313 | 0.016 | 1.000 | |  |  |  |  |  |  |  |
|  |  | 3 | 0.158 | 0.141 | 0.173 | 0.157 | 0.016 | 1.000 | |  |  |  |  |  |  |  |
|  |  | 4 | 0.128 | 0.186 | 0.148 | 0.154 | 0.027 | 1.000 | |  |  |  |  |  |  |  |
|  | siP54nrb | 1 | 0.166 | 0.154 | 0.164 | 0.162 | 0.006 | 0.629 | | 0.665 | | 0.123 | |  |  |  |
|  |  | 2 | 0.169 | 0.172 | 0.172 | 0.171 | 0.002 | 0.545 | |  |  |  |  |  |  |  |
|  |  | 3 | 0.132 | 0.144 | 0.153 | 0.143 | 0.010 | 0.907 | |  |  |  |  |  |  |  |
|  |  | 4 | 0.081 | 0.132 | 0.118 | 0.111 | 0.010 | 0.719 | |  |  |  |  |  |  |  |
| Hypoxia | siCtrl | 1 | 0.343 | 0.355 | 0.353 | 0.351 | 0.007 | 1.000 | | 1.000 | | 0.371 | | *0.0466* | | * |
|  |  | 2 | 0.325 | 0.300 | 0.316 | 0.314 | 0.013 | 1.000 | |  |  |  |  |  |  |  |
|  |  | 3 | 0.256 | 0.242 | 0.159 | 0.219 | 0.052 | 1.000 | |  |  |  |  |  |  |  |
|  |  | 4 | 0.102 | 0.157 | 0.127 | 0.129 | 0.021 | 1.000 | |  |  |  |  |  |  |  |
|  | siP54nrb | 1 | 0.191 | 0.203 | 0.206 | 0.200 | 0.008 | 0.570 | | 0.671 | | 0.178 | |  |  |  |
|  |  | 2 | 0.159 | 0.145 | 0.134 | 0.146 | 0.013 | 0.466 | |  |  |  |  |  |  |  |
|  |  | 3 | 0.166 | 0.259 | 0.212 | 0.212 | 0.047 | 0.970 | |  |  |  |  |  |  |  |
|  |  | 4 | 0.108 | 0.122 | 0.130 | 0.120 | 0.005 | 0.934 | |  |  |  |  |  |  |  |

|  |  |  | Renilla luciferase | | | | | Firefly luciferase | | | | |
| --- | --- | --- | --- | --- | --- | --- | --- | --- | --- | --- | --- | --- |
| **FGF2 IRES** |  | **Exp num** | **A** | **B** | **C** | **Mean** | **SD** | **A** | **B** | **C** | **Mean** | **SD** |
| normoxia | siCtrl | 1 | 338160 | 297348 | 356513 | 330674 | 24727 | 40284 | 42583 | 47910 | 43592 | 3194 |
|  |  | 2 | 190962 | 193130 | 172918 | 185670 | 11097 | 37869 | 28018 | 26650 | 30845 | 6120 |
|  |  | 3 | 192661 | 139882 | 163766 | 165437 | 26429 | 49933 | 47191 | 45737 | 47620 | 2131 |
|  | siP54nrb | 1 | 428321 | 376736 | 381434 | 395497 | 28523 | 22548 | 41972 | 25361 | 29960 | 10497 |
|  |  | 2 | 350806 | 326400 | 298994 | 325400 | 25920 | 46979 | 41071 | 44185 | 44078 | 2955 |
|  |  | 3 | 204274 | 159073 | 175330 | 179559 | 22896 | 51209 | 33872 | 48282 | 44454 | 9281 |
| Hypoxia | siCtrl | 1 | 647702 | 452787 | 497364 | 532618 | 83387 | 61659 | 61503 | 65386 | 62849 | 1795 |
|  |  | 2 | 126638 | 99956 | 87564 | 104719 | 19968 | 18408 | 11584 | 13110 | 14367 | 3581 |
|  |  | 3 | 105830 | 128727 | 123293 | 119283 | 11964 | 28238 | 45322 | 36300 | 36620 | 8546 |
|  | siP54nrb | 1 | 461605 | 534207 | 496440 | 497417 | 29648 | 38628 | 53472 | 50079 | 47393 | 6350 |
|  |  | 2 | 177421 | 155411 | 147051 | 159961 | 15688 | 17991 | 16396 | 15643 | 16677 | 1199 |
|  |  | 3 | 189859 | 106099 | 172669 | 156209 | 44239 | 39168 | 29026 | 44814 | 37669 | 8000 |

|  |  |  | Ratio LucF/LucR | | | | |  |  |  |  |  | |
| --- | --- | --- | --- | --- | --- | --- | --- | --- | --- | --- | --- | --- | --- |
|  |  | **Exp num** | **A** | **B** | **C** | **Mean** | **SD** | **Normalisation** | **Mean** | **SD** | **Mann-Whitney p-value** | | **Significance** |
| normoxia | siCtrl | 1 | 0.119 | 0.143 | 0.134 | 0.132 | 0.012 | 1.000 | 1.000 | 0.393 | *0.1983* | | ns |
|  |  | 2 | 0.198 | 0.145 | 0.154 | 0.166 | 0.028 | 1.000 |  |  |  |  |  |
|  |  | 3 | 0.259 | 0.337 | 0.279 | 0.292 | 0.041 | 1.000 |  |  |  |  |  |
|  | siP54nrb | 1 | 0.053 | 0.111 | 0.066 | 0.077 | 0.031 | 0.581 | 0.778 | 0.396 |  |  |  |
|  |  | 2 | 0.134 | 0.126 | 0.148 | 0.136 | 0.011 | 0.819 |  |  |  |  |  |
|  |  | 3 | 0.251 | 0.213 | 0.275 | 0.246 | 0.031 | 0.844 |  |  |  |  |  |
| hypoxia | siCtrl | 1 | 0.095 | 0.136 | 0.131 | 0.121 | 0.022 | 1.000 | 1.000 | 0.490 | *0.1547* | | ns |
|  |  | 2 | 0.145 | 0.116 | 0.150 | 0.137 | 0.018 | 1.000 |  |  |  |  |  |
|  |  | 3 | 0.267 | 0.352 | 0.294 | 0.304 | 0.043 | 1.000 |  |  |  |  |  |
|  | siP54nrb | 1 | 0.084 | 0.100 | 0.101 | 0.095 | 0.010 | 0.785 | 0.793 | 0.404 |  |  |  |
|  |  | 2 | 0.101 | 0.105 | 0.106 | 0.104 | 0.003 | 0.762 |  |  |  |  |  |
|  |  | 3 | 0.206 | 0.274 | 0.260 | 0.246 | 0.035 | 0.810 |  |  |  |  |  |

|  |  |  | Renilla luciferase | | | | | Firefly luciferase | | | | |
| --- | --- | --- | --- | --- | --- | --- | --- | --- | --- | --- | --- | --- |
| **VEGFAa IRES** |  | **Exp num** | **A** | **B** | **C** | **Mean** | **SD** | **A** | **B** | **C** | **Mean** | **SD** |
| normoxia | siCtrl | 1 | 478052 | 536681 | 460040 | 491591 | 40074 | 6091 | 8319 | 6881 | 7097 | 1130 |
|  |  | 2 | 686015 | 662255 | 669803 | 672691 | 9913 | 3870 | 4191 | 5255 | 4439 | 592 |
|  |  | 3 | 234523 | 254948 | 311971 | 267147 | 40139 | 2790 | 3237 | 4956 | 3661 | 1143 |
|  | siP54nrb | 1 | 151211 | 192635 | 218217 | 187355 | 33814 | 1630 | 1993 | 1954 | 1859 | 199 |
|  |  | 2 | 806710 | 705533 | 883525 | 798589 | 89273 | 1701 | 3698 | 3235 | 2878 | 1045 |
|  |  | 3 | 364632 | 394869 | 414659 | 391387 | 25194 | 4480 | 5186 | 5791 | 5152 | 656 |
| Hypoxia | siCtrl | 1 | 603187 | 551004 | 622577 | 592256 | 37017 | 8658 | 8380 | 8676 | 8571 | 166 |
|  |  | 2 | 1393047 | 1550540 | 1357285 | 1433624 | 83951 | 11436 | 15150 | 15655 | 14080 | 1881 |
|  |  | 3 | 241300 | 357533 | 405013 | 334615 | 84228 | 7744 | 9020 | 9048 | 8604 | 745 |
|  | siP54nrb | 1 | 289694 | 172233 | 213909 | 225279 | 59550 | 1883 | 1497 | 1775 | 1718 | 199 |
|  |  | 2 | 759932 | 1804316 | 2188354 | 1584201 | 603565 | 4715 | 21908 | 24607 | 23258 | 1350 |
|  |  | 3 | 364632 | 394869 | 414659 | 391387 | 25194 | 4480 | 5186 | 5791 | 5152 | 656 |

|  |  |  | Ratio LucF/LucR | | | | |  |  |  |  |  |
| --- | --- | --- | --- | --- | --- | --- | --- | --- | --- | --- | --- | --- |
|  |  | **Exp num** | **A** | **B** | **C** | **Mean** | **SD** | **Normalisation** | **Mean** | **SD** | **Mann-Whitney p-value** | **Significance** |
| normoxia | siCtrl | 1 | 0.013 | 0.016 | 0.015 | 0.014 | 0.001 | 1.000 | 1.000 | 0.343 | *0.1394* | ns |
|  |  | 2 | 0.006 | 0.006 | 0.008 | 0.007 | 0.001 | 1.000 |  |  |  |  |
|  |  | 3 | 0.012 | 0.013 | 0.016 | 0.013 | 0.002 | 1.000 |  |  |  |  |
|  | siP54nrb | 1 | 0.011 | 0.010 | 0.009 | 0.010 | 0.001 | 0.696 | 0.778 | 0.374 |  |  |
|  |  | 2 | 0.002 | 0.005 | 0.004 | 0.004 | 0.002 | 0.556 |  |  |  |  |
|  |  | 3 | 0.012 | 0.013 | 0.014 | 0.013 | 0.001 | 0.973 |  |  |  |  |
| hypoxia | siCtrl | 1 | 0.014 | 0.015 | 0.014 | 0.014 | 0.001 | 1.000 | 1.000 | 0.467 | *0.0244* | * |
|  |  | 2 | 0.008 | 0.010 | 0.012 | 0.010 | 0.002 | 1.000 |  |  |  |  |
|  |  | 3 | 0.032 | 0.025 | 0.022 | 0.027 | 0.005 | 1.000 |  |  |  |  |
|  | siP54nrb | 1 | 0.006 | 0.009 | 0.008 | 0.008 | 0.001 | 0.540 | 0.606 | 0.171 |  |  |
|  |  | 2 | 0.006 | 0.012 | 0.011 | 0.010 | 0.003 | 1.003 |  |  |  |  |
|  |  | 3 | 0.012 | 0.013 | 0.014 | 0.013 | 0.001 | 0.494 |  |  |  |  |

|  |  |  | Renilla luciferase | | | | | Firefly luciferase | | | | |
| --- | --- | --- | --- | --- | --- | --- | --- | --- | --- | --- | --- | --- |
| **VEGFAb IRES** |  | **Exp num** | **A** | **B** | **C** | **Mean** | **SD** | **A** | **B** | **C** | **Mean** | **SD** |
| normoxia | siCtrl | 1 | 2212227 | 2025224 | 2444311 | 2227254 | 171421 | 257293 | 244474 | 310896 | 270888 | 28770 |
|  |  | 2 | 1683303 | 2038361 | 1994936 | 1905533 | 193678 | 197466 | 248062 | 267940 | 237823 | 36336 |
|  |  | 3 | 353069 | 379878 | 485489 | 406145 | 70009 | 45733 | 42142 | 54059 | 47311 | 6113 |
|  | siP54nrb | 1 | 2207682 | 2357474 | 2524435 | 2363197 | 158454 | 204259 | 208483 | 241137 | 217960 | 20183 |
|  |  | 2 | 2664413 | 2662585 | 2817225 | 2714741 | 1293 | 315825 | 385479 | 389540 | 363614 | 49253 |
|  |  | 3 | 480449 | 481870 | 421856 | 461392 | 34246 | 46543 | 57011 | 44830 | 49461 | 6594 |
| Hypoxia | siCtrl | 1 | 3468976 | 4647906 | 6219613 | 5433760 | 1173373 | 309378 | 604844 | 915426 | 760135 | 219614 |
|  |  | 2 | 1184270 | 1246274 | 1144477 | 1191674 | 51300 | 105816 | 111858 | 124528 | 114067 | 9550 |
|  |  | 3 | 350981 | 476613 | 454337 | 427310 | 67035 | 90388 | 93717 | 79327 | 87811 | 7533 |
|  | siP54nrb | 1 | 1663099 | 4715961 | 5314781 | 5015371 | 299410 | 241012 | 644542 | 785788 | 715165 | 70623 |
|  |  | 2 | 1942998 | 1487664 | 4176614 | 1715331 | 321970 | 178076 | 130815 | 539089 | 154446 | 33418 |
|  |  | 3 | 386024 | 404162 | 360362 | 383516 | 22007 | 54030 | 60713 | 64724 | 59822 | 5402 |

|  |  |  | Ratio LucF/LucR | | | | |  |  |  |  |  | |
| --- | --- | --- | --- | --- | --- | --- | --- | --- | --- | --- | --- | --- | --- |
|  |  | **Exp num** | **A** | **B** | **C** | **Mean** | **SD** | **Normalisation** | **Mean** | **SD** | **Mann-Whitney p-value** | | **Significance** |
| normoxia | siCtrl | 1 | 0.116 | 0.121 | 0.127 | 0.121 | 0.005 | 1.000 | 1.000 | 0.067 | *0.2136* | | ns |
|  |  | 2 | 0.117 | 0.122 | 0.134 | 0.124 | 0.009 | 1.000 |  |  |  |  |  |
|  |  | 3 | 0.130 | 0.111 | 0.111 | 0.117 | 0.011 | 1.000 |  |  |  |  |  |
|  | siP54nrb | 1 | 0.093 | 0.088 | 0.096 | 0.092 | 0.004 | 0.759 | 0.918 | 0.168 |  |  |  |
|  |  | 2 | 0.119 | 0.145 | 0.138 | 0.134 | 0.019 | 1.076 |  |  |  |  |  |
|  |  | 3 | 0.097 | 0.118 | 0.106 | 0.107 | 0.011 | 0.914 |  |  |  |  |  |
| hypoxia | siCtrl | 1 | 0.089 | 0.130 | 0.147 | 0.139 | 0.030 | 1.000 | 1.000 | 0.407 | *0.9797* | | ns |
|  |  | 2 | 0.089 | 0.090 | 0.109 | 0.096 | 0.011 | 1.000 |  |  |  |  |  |
|  |  | 3 | 0.258 | 0.197 | 0.175 | 0.210 | 0.043 | 1.000 |  |  |  |  |  |
|  | siP54nrb | 1 | 0.145 | 0.137 | 0.148 | 0.142 | 0.008 | 1.026 | 0.941 | 0.202 |  |  |  |
|  |  | 2 | 0.092 | 0.088 | 0.129 | 0.090 | 0.003 | 0.936 |  |  |  |  |  |
|  |  | 3 | 0.140 | 0.150 | 0.180 | 0.157 | 0.021 | 0.747 |  |  |  |  |  |

|  |  |  | Renilla luciferase | | | | | Firefly luciferase | | | | |
| --- | --- | --- | --- | --- | --- | --- | --- | --- | --- | --- | --- | --- |
| **VEGFC IRES** |  | **Exp num** | **A** | **B** | **C** | **Mean** | **SD** | **A** | **B** | **C** | **Mean** | **SD** |
| normoxia | siCtrl | 1 | 1652281 | 1747731 | 1495968 | 1631994 | 103778 | 73436 | 99773 | 102461 | 91890 | 13095 |
|  |  | 2 | 1750605 | 1126430 | 1746470 | 1541168 | 441358 | 165177 | 104638 | 156405 | 142073 | 42808 |
|  |  | 3 | 448409 | 423742 | 553919 | 475357 | 69146 | 38149 | 39741 | 55704 | 44531 | 9709 |
|  | siP54nrb | 1 | 2650823 | 1576011 | 1731777 | 1986204 | 580823 | 94097 | 69410 | 57191 | 73566 | 18801 |
|  |  | 2 | 2245024 | 2448145 | 2157646 | 2283605 | 149043 | 145454 | 171896 | 155238 | 157529 | 13369 |
|  |  | 3 | 258334 | 310510 | 290358 | 286401 | 26312 | 20846 | 18792 | 20565 | 20068 | 1113 |
| Hypoxia | siCtrl | 1 | 3052125 | 3781966 | 3407369 | 3413820 | 297991 | 103804 | 161246 | 146015 | 137022 | 24297 |
|  |  | 2 | 2172556 | 2224870 | 2838755 | 2198713 | 36992 | 132193 | 132703 | 163425 | 132448 | 361 |
|  |  | 3 | 548838 | 425660 | 473070 | 482523 | 62131 | 48174 | 42953 | 55521 | 48883 | 6314 |
|  | siP54nrb | 1 | 3260370 | 3638044 | 3628119 | 3508844 | 175744 | 108546 | 150891 | 163872 | 141103 | 23623 |
|  |  | 2 | 3765712 | 3336395 | 4380633 | 3827580 | 524861 | 186120 | 185420 | 226711 | 199417 | 23640 |
|  |  | 3 | 538893 | 609759 | 580373 | 576342 | 35604 | 39084 | 46202 | 42424 | 42570 | 3561 |

|  |  |  | Ratio LucF/LucR | | | | |  |  |  |  |  | |
| --- | --- | --- | --- | --- | --- | --- | --- | --- | --- | --- | --- | --- | --- |
|  |  | **Exp num** | **A** | **B** | **C** | **Mean** | **SD** | **Normalisation** | **Mean** | **SD** | **Mann-Whitney p-value** | | **Significance** |
| normoxia | siCtrl | 1 | 0.044 | 0.057 | 0.068 | 0.057 | 0.012 | 1.000 | 1.000 | 0.241 | *0.0333* | | * |
|  |  | 2 | 0.094 | 0.093 | 0.090 | 0.092 | 0.001 | 1.000 |  |  |  |  |  |
|  |  | 3 | 0.085 | 0.094 | 0.101 | 0.093 | 0.008 | 1.000 |  |  |  |  |  |
|  | siP54nrb | 1 | 0.035 | 0.044 | 0.033 | 0.038 | 0.006 | 0.662 | 0.732 | 0.214 |  |  |  |
|  |  | 2 | 0.065 | 0.070 | 0.072 | 0.069 | 0.004 | 0.748 |  |  |  |  |  |
|  |  | 3 | 0.081 | 0.061 | 0.071 | 0.071 | 0.010 | 0.759 |  |  |  |  |  |
| hypoxia | siCtrl | 1 | 0.034 | 0.043 | 0.043 | 0.040 | 0.005 | 1.000 | 1.000 | 0.427 | *0.423* | | ns |
|  |  | 2 | 0.061 | 0.060 | 0.058 | 0.059 | 0.001 | 1.000 |  |  |  |  |  |
|  |  | 3 | 0.088 | 0.101 | 0.117 | 0.102 | 0.015 | 1.000 |  |  |  |  |  |
|  | siP54nrb | 1 | 0.033 | 0.041 | 0.045 | 0.040 | 0.006 | 1.004 | 0.825 | 0.227 |  |  |  |
|  |  | 2 | 0.049 | 0.056 | 0.052 | 0.052 | 0.003 | 0.880 |  |  |  |  |  |
|  |  | 3 | 0.073 | 0.076 | 0.073 | 0.074 | 0.002 | 0.723 |  |  |  |  |  |

|  |  |  | Renilla luciferase | | | | | Firefly luciferase | | | | |
| --- | --- | --- | --- | --- | --- | --- | --- | --- | --- | --- | --- | --- |
| **VEGFD IRES** |  | **Exp num** | **A** | **B** | **C** | **Mean** | **SD** | **A** | **B** | **C** | **Mean** | **SD** |
| normoxia | siCtrl | 1 | 365990 | 427317 | 365164 | 386157 | 29106 | 20158 | 31306 | 30864 | 27443 | 5154 |
|  |  | 2 | 275585 | 220937 | 243324 | 246615 | 27472 | 20951 | 24166 | 16467 | 20528 | 3867 |
|  |  | 3 | 217847 | 327906 | 285000 | 276918 | 55473 | 13957 | 27377 | 23716 | 21683 | 6937 |
|  | siP54nrb | 1 | 493065 | 461440 | 496916 | 483807 | 19466 | 31715 | 25746 | 32021 | 29828 | 3538 |
|  |  | 2 | 567823 | 549217 | 372141 | 496394 | 108008 | 33285 | 37862 | 23280 | 31476 | 7457 |
|  |  | 3 | 592115 | 670999 | 704955 | 656023 | 57892 | 44366 | 59232 | 60346 | 54648 | 8922 |
| Hypoxia | siCtrl | 1 | 440436 | 602979 | 487986 | 510467 | 68236 | 48804 | 59199 | 48550 | 52184 | 4961 |
|  |  | 2 | 430900 | 671630 | 616169 | 572899 | 126063 | 36827 | 54089 | 43938 | 44951 | 8675 |
|  |  | 3 | 249034 | 353678 | 287905 | 296872 | 52895 | 29619 | 31439 | 27641 | 29566 | 1899 |
|  | siP54nrb | 1 | 758794 | 743577 | 1011622 | 837998 | 122928 | 60275 | 76085 | 101933 | 79431 | 17171 |
|  |  | 2 | 830670 | 773593 | 571751 | 725338 | 136037 | 56451 | 58640 | 37000 | 50697 | 11912 |
|  |  | 3 | 390927 | 304315 | 223146 | 306129 | 83905 | 26926 | 26335 | 18850 | 24037 | 4502 |

|  |  |  | Ratio LucF/LucR | | | | |  |  |  |  |  | |
| --- | --- | --- | --- | --- | --- | --- | --- | --- | --- | --- | --- | --- | --- |
|  |  | **Exp num** | **A** | **B** | **C** | **Mean** | **SD** | **Normalisation** | **Mean** | **SD** | **Mann-Whitney p-value** | | **Significance** |
| normoxia | siCtrl | 1 | 0.055 | 0.073 | 0.085 | 0.071 | 0.015 | 1.000 | 1.000 | 0.201 | *0.3272* | | ns |
|  |  | 2 | 0.076 | 0.109 | 0.068 | 0.084 | 0.022 | 1.000 |  |  |  |  |  |
|  |  | 3 | 0.064 | 0.083 | 0.083 | 0.077 | 0.011 | 1.000 |  |  |  |  |  |
|  | siP54nrb | 1 | 0.064 | 0.056 | 0.064 | 0.062 | 0.005 | 0.867 | 0.895 | 0.148 |  |  |  |
|  |  | 2 | 0.059 | 0.069 | 0.063 | 0.063 | 0.005 | 0.751 |  |  |  |  |  |
|  |  | 3 | 0.075 | 0.088 | 0.086 | 0.083 | 0.007 | 1.078 |  |  |  |  |  |
| hypoxia | siCtrl | 1 | 0.111 | 0.098 | 0.099 | 0.103 | 0.007 | 1.000 | 1.000 | 0.157 | *0.0934* | | ns |
|  |  | 2 | 0.085 | 0.081 | 0.071 | 0.079 | 0.007 | 1.000 |  |  |  |  |  |
|  |  | 3 | 0.119 | 0.089 | 0.096 | 0.101 | 0.016 | 1.000 |  |  |  |  |  |
|  | siP54nrb | 1 | 0.079 | 0.102 | 0.101 | 0.094 | 0.013 | 0.916 | 0.860 | 0.145 |  |  |  |
|  |  | 2 | 0.068 | 0.076 | 0.065 | 0.069 | 0.006 | 0.878 |  |  |  |  |  |
|  |  | 3 | 0.069 | 0.087 | 0.084 | 0.080 | 0.010 | 0.790 |  |  |  |  |  |

|  |  |  | Renilla luciferase | | | | | Firefly luciferase | | | | |
| --- | --- | --- | --- | --- | --- | --- | --- | --- | --- | --- | --- | --- |
| **IGF1R IRES** |  | **Exp num** | **A** | **B** | **C** | **Mean** | **SD** | **A** | **B** | **C** | **Mean** | **SD** |
| normoxia | siCtrl | 1 | 568295 | 624464 | 574870 | 589210 | 25073 | 88840 | 130924 | 115377 | 111714 | 17375 |
|  |  | 2 | 255284 | 263849 | 293168 | 270767 | 19867 | 103285 | 90152 | 119297 | 104245 | 14596 |
|  |  | 3 | 254097 | 202990 | 248049 | 235045 | 27925 | 76689 | 80603 | 85770 | 81021 | 4555 |
|  | siP54nrb | 1 | 763746 | 750413 | 563812 | 757079 | 111782 | 76734 | 105925 | 19969 | 91329 | 20641 |
|  |  | 2 | 319438 | 405317 | 373312 | 366022 | 43401 | 96134 | 121000 | 114481 | 110538 | 12893 |
|  |  | 3 | 252281 | 328205 | 261075 | 280521 | 41530 | 104078 | 132182 | 105496 | 113919 | 15832 |
| Hypoxia | siCtrl | 1 | 1674693 | 1508464 | 1456958 | 1546705 | 92912 | 193429 | 238652 | 242544 | 224875 | 22292 |
|  |  | 2 | 210319 | 236396 | 346932 | 264549 | 72527 | 98731 | 90687 | 136145 | 108521 | 24259 |
|  |  | 3 | 294943 | 209447 | 245953 | 250114 | 42899 | 156272 | 138708 | 126516 | 140499 | 14959 |
|  | siP54nrb | 1 | 1821099 | 1509328 | 1813745 | 1714724 | 145268 | 199152 | 219058 | 287110 | 235107 | 37659 |
|  |  | 2 | 302340 | 322463 | 359354 | 328052 | 28915 | 131203 | 131468 | 133843 | 132171 | 1454 |
|  |  | 3 | 340401 | 361295 | 421935 | 374544 | 42351 | 188827 | 171762 | 198048 | 186212 | 13336 |

|  |  |  | Ratio LucF/LucR | | | | |  |  |  |  |  | |
| --- | --- | --- | --- | --- | --- | --- | --- | --- | --- | --- | --- | --- | --- |
|  |  | **Exp num** | **A** | **B** | **C** | **Mean** | **SD** | **Normalisation** | **Mean** | **SD** | **Mann-Whitney p-value** | | **Significance** |
| normoxia | siCtrl | 1 | 0.156 | 0.210 | 0.201 | 0.189 | 0.029 | 1.000 | 1.000 | 0.313 | *0.5346* | | ns |
|  |  | 2 | 0.405 | 0.342 | 0.407 | 0.384 | 0.037 | 1.000 |  |  |  |  |  |
|  |  | 3 | 0.302 | 0.397 | 0.346 | 0.348 | 0.048 | 1.000 |  |  |  |  |  |
|  | siP54nrb | 1 | 0.100 | 0.141 | 0.035 | 0.121 | 0.029 | 0.640 | 0.869 | 0.459 |  |  |  |
|  |  | 2 | 0.301 | 0.299 | 0.307 | 0.302 | 0.004 | 0.786 |  |  |  |  |  |
|  |  | 3 | 0.413 | 0.403 | 0.404 | 0.406 | 0.005 | 1.167 |  |  |  |  |  |
| hypoxia | siCtrl | 1 | 0.116 | 0.158 | 0.166 | 0.147 | 0.027 | 1.000 | 1.000 | 0.508 | *0.6502* | | ns |
|  |  | 2 | 0.469 | 0.384 | 0.392 | 0.415 | 0.047 | 1.000 |  |  |  |  |  |
|  |  | 3 | 0.530 | 0.662 | 0.514 | 0.569 | 0.081 | 1.000 |  |  |  |  |  |
|  | siP54nrb | 1 | 0.109 | 0.145 | 0.158 | 0.138 | 0.025 | 0.938 | 0.922 | 0.439 |  |  |  |
|  |  | 2 | 0.434 | 0.408 | 0.372 | 0.405 | 0.031 | 0.975 |  |  |  |  |  |
|  |  | 3 | 0.555 | 0.475 | 0.469 | 0.500 | 0.048 | 0.879 |  |  |  |  |  |

|  |  |  | Renilla luciferase | | | | | Firefly luciferase | | | | |
| --- | --- | --- | --- | --- | --- | --- | --- | --- | --- | --- | --- | --- |
| **c-myc IRES** |  | **Exp num** | **A** | **B** | **C** | **Mean** | **SD** | **A** | **B** | **C** | **Mean** | **SD** |
| normoxia | siCtrl | 1 | 2337917 | 2493989 | 2024414 | 2285440 | 195261 | 240239 | 402619 | 259913 | 300923 | 72357 |
|  |  | 2 | 936981 | 912821 | 923416 | 924406 | 12111 | 353453 | 426897 | 451667 | 410672 | 51078 |
|  |  | 3 | 996324 | 852009 | 1203390 | 1017241 | 176622 | 202668 | 208450 | 280358 | 230492 | 43282 |
|  | siP54nrb | 1 | 2233970 | 2621631 | 1839095 | 2231565 | 391274 | 73105 | 120223 | 122124 | 105151 | 27769 |
|  |  | 2 | 1221473 | 1402859 | 1186112 | 1270148 | 116283 | 364957 | 379515 | 379515 | 374662 | 8405 |
|  |  | 3 | 1449138 | 1489153 | 1240142 | 1392811 | 133721 | 303849 | 264306 | 264306 | 277487 | 22830 |
| Hypoxia | siCtrl | 1 | 4682318 | 4074549 | 3763596 | 4173488 | 381536 | 522595 | 516285 | 499798 | 512893 | 9611 |
|  |  | 2 | 802990 | 789755 | 806209 | 799651 | 8720 | 392506 | 481930 | 399124 | 424520 | 49828 |
|  |  | 3 | 1079348 | 1019947 | 1437006 | 1178767 | 225605 | 243035 | 311946 | 408904 | 321295 | 83329 |
|  | siP54nrb | 1 | 4646883 | 6584091 | 5894884 | 6239487 | 844550 | 470519 | 991993 | 947313 | 969653 | 22340 |
|  |  | 2 | 774836 | 734712 | 594396 | 701314 | 94743 | 334975 | 410691 | 410691 | 385452 | 43715 |
|  |  | 3 | 1368840 | 729774 | 968602 | 1022406 | 322912 | 386054 | 283090 | 283090 | 317411 | 59446 |

|  |  |  | Ratio LucF/LucR | | | | |  |  |  |  |  | |
| --- | --- | --- | --- | --- | --- | --- | --- | --- | --- | --- | --- | --- | --- |
|  |  | **Exp num** | **A** | **B** | **C** | **Mean** | **SD** | **normalisation** | **Mean** | **SD** | **Mann-Whitney p-value** | | **significance** |
| normoxia | siCtrl | 1 | 0.103 | 0.161 | 0.128 | 0.131 | 0.029 | 1.000 | 1.000 | 0.536 | *0.3347* | | ns |
|  |  | 2 | 0.377 | 0.468 | 0.489 | 0.445 | 0.059 | 1.000 |  |  |  |  |  |
|  |  | 3 | 0.203 | 0.245 | 0.233 | 0.227 | 0.021 | 1.000 |  |  |  |  |  |
|  | siP54nrb | 1 | 0.033 | 0.046 | 0.066 | 0.048 | 0.017 | 0.369 | 0.679 | 0.410 |  |  |  |
|  |  | 2 | 0.299 | 0.271 | 0.320 | 0.296 | 0.025 | 0.667 |  |  |  |  |  |
|  |  | 3 | 0.210 | 0.177 | 0.213 | 0.200 | 0.020 | 0.881 |  |  |  |  |  |
| hypoxia | siCtrl | 1 | 0.112 | 0.127 | 0.133 | 0.124 | 0.011 | 1.000 | 1.000 | 0.593 | *0.7775* | | ns |
|  |  | 2 | 0.489 | 0.610 | 0.495 | 0.531 | 0.068 | 1.000 |  |  |  |  |  |
|  |  | 3 | 0.225 | 0.306 | 0.285 | 0.272 | 0.042 | 1.000 |  |  |  |  |  |
|  | siP54nrb | 1 | 0.101 | 0.151 | 0.161 | 0.156 | 0.007 | 1.258 | 1.099 | 0.640 |  |  |  |
|  |  | 2 | 0.432 | 0.559 | 0.691 | 0.561 | 0.129 | 1.055 |  |  |  |  |  |
|  |  | 3 | 0.282 | 0.388 | 0.292 | 0.321 | 0.058 | 1.180 |  |  |  |  |  |

|  |  |  | Renilla luciferase | | | | | Firefly luciferase | | | | |
| --- | --- | --- | --- | --- | --- | --- | --- | --- | --- | --- | --- | --- |
| **EMCV IRES** |  | **Exp num** | **A** | **B** | **C** | **Mean** | **SD** | **A** | **B** | **C** | **Mean** | **SD** |
| normoxia | siCtrl | 1 | 6365162 | 8322840 | 7640560 | 7442854 | 993701 | 6103347 | 9113681 | 8490981 | 7902670 | 1589060 |
|  |  | 2 | 3307064 | 2859152 | 2994865 | 3053694 | 229678 | 4239469 | 4033743 | 4582152 | 4285121 | 277040 |
|  |  | 3 | 2337917 | 3146033 | 2611255 | 2698402 | 335617 | 1603385 | 2929991 | 2033089 | 2188822 | 552666 |
|  | siP54nrb | 1 | 7970781 | 10596055 | 7928241 | 8831692 | 1528131 | 11475614 | 14992302 | 9480997 | 11982971 | 2790462 |
|  |  | 2 | 1258884 | 1484865 | 1279441 | 1341063 | 124959 | 1393602 | 2201468 | 2095182 | 1896751 | 438968 |
|  |  | 3 | 2108239 | 1965979 | 1451461 | 1841893 | 345525 | 1398909 | 965448 | 698436 | 1020931 | 353517 |
| Hypoxia | siCtrl | 1 | 7931570 | 7957225 | 6740129 | 7542975 | 695403 | 12387597 | 12394766 | 9494868 | 11425744 | 1672191 |
|  |  | 2 | 3204089 | 2728506 | 2221130 | 2717908 | 491566 | 5358325 | 4472124 | 3540859 | 4457103 | 908826 |
|  |  | 3 | 4682318 | 7943966 | 6605200 | 6410495 | 1338661 | 6266686 | 7709654 | 6349910 | 6775417 | 661478 |
|  | siP54nrb | 1 | 9797962 | 9225136 | 9956752 | 9659950 | 384839 | 17275460 | 16007194 | 16719339 | 16667331 | 635730 |
|  |  | 2 | 2133610 | 2023162 | 2087983 | 2081585 | 55501 | 2636753 | 2775357 | 2968316 | 2793475 | 166523 |
|  |  | 3 | 4646883 | 9684636 | 7328583 | 7220034 | 2058086 | 5094831 | 8216051 | 7029763 | 6780215 | 1286393 |

|  |  |  | Ratio LucF/LucR | | | | |  |  |  |  |  | |
| --- | --- | --- | --- | --- | --- | --- | --- | --- | --- | --- | --- | --- | --- |
|  |  | **Exp num** | **A** | **B** | **C** | **Mean** | **SD** | **Normalisation** | **Mean** | **SD** | **Mann-Whitney p-value** | | **Significance** |
| normoxia | siCtrl | 1 | 0.959 | 1.095 | 1.111 | 1.055 | 0.084 | 1.000 | 1.000 | 0.259 | *0.7775* | | *ns* |
|  |  | 2 | 1.282 | 1.411 | 1.530 | 1.408 | 0.124 | 1.000 |  |  |  |  |  |
|  |  | 3 | 0.686 | 0.931 | 0.779 | 0.799 | 0.124 | 1.000 |  |  |  |  |  |
|  | siP54nrb | 1 | 1.440 | 1.415 | 1.196 | 1.350 | 0.134 | 1.280 | 1.013 | 0.412 |  |  |  |
|  |  | 2 | 1.107 | 1.483 | 1.638 | 1.409 | 0.273 | 1.001 |  |  |  |  |  |
|  |  | 3 | 0.664 | 0.491 | 0.481 | 0.545 | 0.103 | 0.683 |  |  |  |  |  |
| hypoxia | siCtrl | 1 | 1.562 | 1.558 | 1.409 | 1.509 | 0.087 | 1.000 | 1.000 | 0.194 | *0.8427* | | *ns* |
|  |  | 2 | 1.672 | 1.639 | 1.594 | 1.635 | 0.039 | 1.000 |  |  |  |  |  |
|  |  | 3 | 1.338 | 0.971 | 0.961 | 1.090 | 0.215 | 1.000 |  |  |  |  |  |
|  | siP54nrb | 1 | 1.763 | 1.735 | 1.679 | 1.726 | 0.043 | 1.143 | 0.953 | 0.240 |  |  |  |
|  |  | 2 | 1.236 | 1.372 | 1.422 | 1.343 | 0.096 | 0.821 |  |  |  |  |  |
|  |  | 3 | 1.096 | 0.848 | 0.959 | 0.968 | 0.124 | 0.888 |  |  |  |  |  |

**Supplementary file 5. IRES activities in normoxic and hypoxic HL-1 cells after p54^nrb^ knock-down.**

HL-1 cells were transduced with Lucky Luke bicistronic lentivectors containing the IRES of FGF1, FGF2, VEGFA (a or b), VEGFC, VEGFD, IGF1R, c-myc or EMCV. Cells were then treated with siP54^nrb^ or siControl (siCtrl) smartpool during normoxia or hypoxia 1% O_2._ Renilla and firefly luciferase activities were measured (upper panel, or page 1 for FGF1 IRES) and the IRES activities evaluated with the ratio LucF/LucR (lower panel, or page 2 for FGF1 IRES). Mann-Whitney test was performed with n=9 (n=12 for FGF1 IRES). *p<0.05, **p<0.01, ***<0.001, ****p<0.0001. For each IRES the mean has been calculated with nine cell culture biological replicates (Experiments A, B, C correspond to experiments performed at different dates, while 1, 2, 3 are experiments performed in parallel at the same date, each of them being already the mean of three technical replicates (27 technical replicates in total).
