## Supplementary files 1-8 for "Long non-coding RNA Neat1 and paraspeckle components are translational regulators in hypoxia": V2 Supplementary file 6 PSPC1 KD.docx

|  |  |  | | Renilla luciferase | | | | | | Firefly luciferase | | | | |
| --- | --- | --- | --- | --- | --- | --- | --- | --- | --- | --- | --- | --- | --- | --- |
| **FGF1 IRES** |  | **Exp num** | **A** | | **B** | **C** | **Mean** | **SD** | **A** | | **B** | **C** | **Mean** | **SD** |
| normoxia | siCtrl | 1 | 4321498 | | 6005077 | 5319145 | 5215240 | 691234 | 908764 | | 1480053 | 1477175 | 1288664 | 268633 |
|  |  | 2 | 1955203 | | 1917089 | 2081085 | 1984459 | 70074 | 338768 | | 304406 | 369720 | 337631 | 26676 |
|  |  | 3 | 1590009 | | 1381692 | 1381001 | 1450901 | 120472 | 251834 | | 194834 | 238849 | 228506 | 29875 |
|  |  | 4 | 659990 | | 678158 | 540834 | 626327 | 74595 | 140428 | | 158968 | 119763 | 139720 | 19612 |
|  | siPSPC1 | 1 | 4082147 | | 7774218 | 6274884 | 6043749 | 1516117 | 980547 | | 1378223 | 1184000 | 1180923 | 162365 |
|  |  | 2 | 3311503 | | 2283125 | 2768769 | 2787799 | 420049 | 392942 | | 307052 | 331271 | 343755 | 36159 |
|  |  | 3 | 1955592 | | 1606440 | 1859696 | 1807243 | 180389 | 279558 | | 220270 | 240182 | 246670 | 30172 |
|  |  | 4 | 861047 | | 966693 | 868097 | 898612 | 59065 | 160531 | | 157534 | 158640 | 158902 | 1516 |
| Hypoxia | siCtrl | 1 | 6004439 | | 5725646 | 6186192 | 5972092 | 189403 | 1768671 | | 1788136 | 1744898 | 1767235 | 17681 |
|  |  | 2 | 2151385 | | 2008858 | 1959317 | 2039853 | 81417 | 322854 | | 298111 | 329513 | 316826 | 13510 |
|  |  | 3 | 1121156 | | 1093631 | 1022224 | 1079004 | 51062 | 287035 | | 264567 | 162669 | 238090 | 66276 |
|  |  | 4 | 888672 | | 1025609 | 858460 | 924247 | 89072 | 195709 | | 210897 | 176556 | 194388 | 17209 |
|  | siPSPC1 | 1 | 8045681 | | 8184503 | 8570314 | 8266832 | 221951 | 1521908 | | 1855397 | 1781743 | 1719683 | 143044 |
|  |  | 2 | 3486479 | | 3307271 | 3112770 | 3302173 | 152609 | 395606 | | 434495 | 392077 | 407392 | 19218 |
|  |  | 3 | 1119657 | | 1364902 | 1515406 | 1333321 | 199756 | 270801 | | 270678 | 185120 | 242200 | 49432 |
|  |  | 4 | 998170 | | 1230674 | 980030 | 1069625 | 139768 | 203261 | | 218574 | 153798 | 191878 | 33855 |

|  |  |  | Ratio LucF/LucR | | | | |  |  |  |  |  |
| --- | --- | --- | --- | --- | --- | --- | --- | --- | --- | --- | --- | --- |
| **FGF1 IRES** |  | **Exp num** | **A** | **B** | **C** | **Mean** | **SD** | **Normalisation** | **Mean** | **SD** | **Mann-Whitney p-value** | **Significance** |
| normoxia | siCtrl | 1 | 0.210 | 0.246 | 0.278 | 0.245 | 0.034 | 1.000 | 1.000 | 0.209 | *0.0432* | * |
|  |  | 2 | 0.173 | 0.159 | 0.178 | 0.170 | 0.010 | 1.000 |  |  |  |  |
|  |  | 3 | 0.158 | 0.141 | 0.173 | 0.157 | 0.016 | 1.000 |  |  |  |  |
|  |  | 4 | 0.213 | 0.234 | 0.221 | 0.223 | 0.011 | 1.000 |  |  |  |  |
|  | siPSPC1 | 1 | 0.240 | 0.177 | 0.189 | 0.202 | 0.034 | 0.825 | 0.805 | 0.183 |  |  |
|  |  | 2 | 0.119 | 0.134 | 0.120 | 0.124 | 0.009 | 0.731 |  |  |  |  |
|  |  | 3 | 0.143 | 0.137 | 0.129 | 0.136 | 0.007 | 0.866 |  |  |  |  |
|  |  | 4 | 0.186 | 0.163 | 0.183 | 0.177 | 0.013 | 0.796 |  |  |  |  |
| Hypoxia | siCtrl | 1 | 0.295 | 0.312 | 0.282 | 0.296 | 0.015 | 1.000 | 1.000 | 0.262 | *0.0537* | ns |
|  |  | 2 | 0.150 | 0.148 | 0.168 | 0.156 | 0.011 | 1.000 |  |  |  |  |
|  |  | 3 | 0.256 | 0.242 | 0.159 | 0.219 | 0.052 | 1.000 |  |  |  |  |
|  |  | 4 | 0.220 | 0.206 | 0.206 | 0.211 | 0.008 | 1.000 |  |  |  |  |
|  | siPSPC1 | 1 | 0.189 | 0.227 | 0.208 | 0.208 | 0.019 | 0.702 | 0.792 | 0.198 |  |  |
|  |  | 2 | 0.113 | 0.131 | 0.126 | 0.124 | 0.009 | 0.795 |  |  |  |  |
|  |  | 3 | 0.242 | 0.198 | 0.122 | 0.187 | 0.061 | 0.856 |  |  |  |  |
|  |  | 4 | 0.204 | 0.178 | 0.157 | 0.179 | 0.023 | 0.852 |  |  |  |  |

|  |  |  | Renilla luciferase | | | | | Firefly luciferase | | | | |
| --- | --- | --- | --- | --- | --- | --- | --- | --- | --- | --- | --- | --- |
| **FGF2 IRES** |  | **Exp num** | **A** | **B** | **C** | **Mean** | **SD** | **A** | **B** | **C** | **Mean** | **SD** |
| normoxia | siCtrl | 1 | 629345 | 620635 | 639942 | 629974 | 9669 | 125772 | 127427 | 104526 | 119242 | 12771 |
|  |  | 2 | 536863 | 551760 | 514298 | 534307 | 18861 | 107176 | 122524 | 107326 | 112342 | 8818 |
|  |  | 3 | 192661 | 139882 | 163766 | 165437 | 26429 | 49933 | 47191 | 45737 | 47620 | 2131 |
|  | siPSPC1 | 1 | 737879 | 856918 | 801706 | 798834 | 59571 | 96448 | 119580 | 107428 | 107819 | 11571 |
|  |  | 2 | 696996 | 667848 | 735212 | 700019 | 33783 | 99149 | 88683 | 101143 | 96325 | 6693 |
|  |  | 3 | 198025 | 148189 | 184532 | 176915 | 25777 | 41765 | 38916 | 42210 | 40964 | 1788 |
| Hypoxia | siCtrl | 1 | 680776 | 726150 | 752265 | 719730 | 36174 | 176583 | 188387 | 180124 | 181698 | 6057 |
|  |  | 2 | 772499 | 915914 | 922373 | 870262 | 84727 | 181399 | 209640 | 191629 | 194223 | 14298 |
|  |  | 3 | 105830 | 128727 | 123293 | 119283 | 11964 | 28238 | 45322 | 36300 | 36620 | 8546 |
|  | siPSPC1 | 1 | 933123 | 981337 | 910931 | 941797 | 35995 | 191281 | 174390 | 151232 | 172301 | 20106 |
|  |  | 2 | 998304 | 1047005 | 972539 | 1005949 | 37817 | 152691 | 157337 | 160958 | 156996 | 4144 |
|  |  | 3 | 154734 | 154943 | 132869 | 147515 | 12685 | 36101 | 27403 | 28769 | 30758 | 4678 |

|  |  |  | Ratio LucF/LucR | | | | |  |  |  |  |  |
| --- | --- | --- | --- | --- | --- | --- | --- | --- | --- | --- | --- | --- |
|  |  | **Exp num** | **A** | **B** | **C** | **Mean** | **SD** | **Normalisation** | **Mean** | **SD** | **Mann-Whitney p-value** | **Significance** |
| normoxia | siCtrl | 1 | 0.200 | 0.205 | 0.163 | 0.189 | 0.023 | 1.000 | 1.000 | 0.229 | *0.0474* | * |
|  |  | 2 | 0.200 | 0.222 | 0.209 | 0.210 | 0.011 | 1.000 |  |  |  |  |
|  |  | 3 | 0.259 | 0.337 | 0.279 | 0.292 | 0.041 | 1.000 |  |  |  |  |
|  | siPSPC1 | 1 | 0.131 | 0.140 | 0.134 | 0.135 | 0.004 | 0.711 | 0.732 | 0.220 |  |  |
|  |  | 2 | 0.142 | 0.133 | 0.138 | 0.138 | 0.005 | 0.655 |  |  |  |  |
|  |  | 3 | 0.211 | 0.263 | 0.229 | 0.234 | 0.026 | 0.802 |  |  |  |  |
| Hypoxia | siCtrl | 1 | 0.259 | 0.259 | 0.239 | 0.253 | 0.012 | 1.000 | 1.000 | 0.163 | *0.0003* | **** |
|  |  | 2 | 0.235 | 0.229 | 0.208 | 0.224 | 0.014 | 1.000 |  |  |  |  |
|  |  | 3 | 0.267 | 0.352 | 0.294 | 0.304 | 0.043 | 1.000 |  |  |  |  |
|  | siPSPC1 | 1 | 0.205 | 0.178 | 0.166 | 0.183 | 0.020 | 0.724 | 0.702 | 0.112 |  |  |
|  |  | 2 | 0.153 | 0.150 | 0.166 | 0.156 | 0.008 | 0.698 |  |  |  |  |
|  |  | 3 | 0.233 | 0.177 | 0.217 | 0.209 | 0.029 | 0.686 |  |  |  |  |

|  |  |  | Renilla luciferase | | | | | Firefly luciferase | | | | |
| --- | --- | --- | --- | --- | --- | --- | --- | --- | --- | --- | --- | --- |
| **VEGFAa IRES** |  | **Exp num** | **A** | **B** | **C** | **Mean** | **SD** | **A** | **B** | **C** | **Mean** | **SD** |
| normoxia | siCtrl | 1 | 1004059 | 961239 | 1082186 | 1015828 | 61326 | 9676 | 9919 | 10680 | 10091 | 524 |
|  |  | 2 | 858315 | 1006460 | 1131780 | 998852 | 136891 | 10177 | 10903 | 9795 | 10292 | 563 |
|  |  | 3 | 234523 | 254948 | 311971 | 267147 | 40139 | 2790 | 3237 | 4956 | 3661 | 1143 |
|  | siPSPC1 | 1 | 1074839 | 1201565 | 1237390 | 1171265 | 85406 | 11143 | 10556 | 10470 | 10723 | 366 |
|  |  | 2 | 1388730 | 1330000 | 1266025 | 1328252 | 61371 | 1289172 | 2652301 | 1266025 | 1735833 | 793769 |
|  |  | 3 | 362066 | 475749 | 428052 | 421956 | 57086 | 3371 | 4835 | 4230 | 4145 | 736 |
| Hypoxia | siCtrl | 1 | 1733484 | 1540310 | 1514623 | 1596139 | 119636 | 21593 | 19968 | 21705 | 21089 | 972 |
|  |  | 2 | 1201671 | 1239986 | 1269291 | 1236983 | 33910 | 21334 | 25229 | 24602 | 23722 | 2091 |
|  |  | 3 | 241300 | 357533 | 405013 | 334615 | 84228 | 7744 | 9020 | 9048 | 8604 | 745 |
|  | siPSPC1 | 1 | 1554638 | 1354818 | 866881 | 1258779 | 353794 | 17426 | 16509 | 15533 | 16489 | 947 |
|  |  | 2 | 1354727 | 1241900 | 1583299 | 1393308 | 173939 | 1356270 | 2145236 | 1583299 | 1694935 | 406157 |
|  |  | 3 | 388321 | 411510 | 398435 | 399422 | 11626 | 6566 | 6841 | 7877 | 7095 | 691 |

|  |  |  | Ratio LucF/LucR | | | | |  |  |  |  |  | |
| --- | --- | --- | --- | --- | --- | --- | --- | --- | --- | --- | --- | --- | --- |
|  |  | **Exp num** | **A** | **B** | **C** | **Mean** | **SD** | **Normalisation** | **Mean** | **SD** | **Mann-Whitney p-value** | | **Significance** |
| normoxia | siCtrl | 1 | 0.010 | 0.010 | 0.010 | 0.010 | 0.000 | 1.000 | 1.000 | 0.190 | *0.0063* | | ** |
|  |  | 2 | 0.012 | 0.011 | 0.009 | 0.010 | 0.002 | 1.000 |  |  |  |  |  |
|  |  | 3 | 0.012 | 0.013 | 0.016 | 0.013 | 0.002 | 1.000 |  |  |  |  |  |
|  | siPSPC1 | 1 | 0.010 | 0.009 | 0.008 | 0.009 | 0.001 | 0.926 | 0.820 | 0.064 |  |  |  |
|  |  | 2 | 0.009 | 0.009 | 0.009 | 0.009 | 0.000 | 0.840 |  |  |  |  |  |
|  |  | 3 | 0.009 | 0.010 | 0.010 | 0.010 | 0.000 | 0.725 |  |  |  |  |  |
| Hypoxia | siCtrl | 1 | 0.012 | 0.013 | 0.014 | 0.013 | 0.001 | 1.000 | 1.000 | 0.323 | *0.0884* | | ns |
|  |  | 2 | 0.018 | 0.020 | 0.019 | 0.019 | 0.001 | 1.000 |  |  |  |  |  |
|  |  | 3 | 0.032 | 0.025 | 0.022 | 0.027 | 0.005 | 1.000 |  |  |  |  |  |
|  | siPSPC1 | 1 | 0.011 | 0.012 | 0.018 | 0.014 | 0.004 | 1.039 | 0.763 | 0.148 |  |  |  |
|  |  | 2 | 0.014 | 0.013 | 0.013 | 0.013 | 0.001 | 0.702 |  |  |  |  |  |
|  |  | 3 | 0.017 | 0.017 | 0.020 | 0.018 | 0.002 | 0.669 |  |  |  |  |  |

|  |  |  | Renilla luciferase | | | | | Firefly luciferase | | | | |
| --- | --- | --- | --- | --- | --- | --- | --- | --- | --- | --- | --- | --- |
| **VEGFAb IRES** |  | **Exp num** | **A** | **B** | **C** | **Mean** | **SD** | **A** | **B** | **C** | **Mean** | **SD** |
| normoxia | siCtrl | 1 | 3953413 | 4461082 | 4955118 | 4456538 | 500868 | 636888 | 673362 | 711456 | 673902 | 37287 |
|  |  | 2 | 3617674 | 4224329 | 4116125 | 3986042 | 323572 | 597951 | 643573 | 666618 | 636047 | 34946 |
|  |  | 3 | 353069 | 379878 | 485489 | 406145 | 70009 | 45733 | 42142 | 54059 | 47311 | 6113 |
|  | siPSPC1 | 1 | 4899771 | 5347093 | 5652256 | 5299707 | 378474 | 761340 | 852226 | 883493 | 832353 | 63455 |
|  |  | 2 | 4923495 | 5294098 | 5487600 | 5235064 | 286648 | 680323 | 735719 | 713411 | 709817 | 27872 |
|  |  | 3 | 523779 | 465105 | 462089 | 483658 | 34779 | 49257 | 48436 | 44309 | 47334 | 2652 |
| Hypoxia | siCtrl | 1 | 5227968 | 5137726 | 5358864 | 5241519 | 111190 | 1455967 | 1495875 | 1461388 | 1471077 | 21646 |
|  |  | 2 | 5214317 | 6525712 | 4929908 | 5556646 | 851199 | 748890 | 981243 | 649085 | 793073 | 170430 |
|  |  | 3 | 350981 | 476613 | 454337 | 427310 | 67035 | 90388 | 93717 | 79327 | 87811 | 7533 |
|  | siPSPC1 | 1 | 5942477 | 6418246 | 6420607 | 6260443 | 275370 | 1481303 | 1800613 | 1778707 | 1686874 | 178367 |
|  |  | 2 | 6960135 | 6471371 | 5894089 | 6441865 | 533635 | 954043 | 1033952 | 1003397 | 997131 | 40322 |
|  |  | 3 | 440546 | 506296 | 499188 | 482010 | 36084 | 63614 | 90548 | 73343 | 75835 | 13639 |

|  |  |  | Ratio LucF/LucR | | | | |  |  |  |  |  |
| --- | --- | --- | --- | --- | --- | --- | --- | --- | --- | --- | --- | --- |
|  |  | **Exp num** | **A** | **B** | **C** | **Mean** | **SD** | **Normalisation** | **Mean** | **SD** | **Mann-Whitney p-value** | **Significance** |
| normoxia | siCtrl | 1 | 0.161 | 0.151 | 0.144 | 0.152 | 0.009 | 1.000 | 1.000 | 0.147 | *0.2307* | ns |
|  |  | 2 | 0.165 | 0.152 | 0.162 | 0.160 | 0.007 | 1.000 |  |  |  |  |
|  |  | 3 | 0.130 | 0.111 | 0.111 | 0.117 | 0.011 | 1.000 |  |  |  |  |
|  | siPSPC1 | 1 | 0.155 | 0.159 | 0.156 | 0.157 | 0.002 | 1.034 | 0.911 | 0.183 |  |  |
|  |  | 2 | 0.138 | 0.139 | 0.130 | 0.136 | 0.005 | 0.849 |  |  |  |  |
|  |  | 3 | 0.094 | 0.104 | 0.096 | 0.098 | 0.005 | 0.836 |  |  |  |  |
| Hypoxia | siCtrl | 1 | 0.278 | 0.291 | 0.273 | 0.281 | 0.009 | 1.000 | 1.000 | 0.305 | *0.6219* | ns |
|  |  | 2 | 0.144 | 0.150 | 0.132 | 0.142 | 0.009 | 1.000 |  |  |  |  |
|  |  | 3 | 0.258 | 0.197 | 0.175 | 0.210 | 0.043 | 1.000 |  |  |  |  |
|  | siPSPC1 | 1 | 0.249 | 0.281 | 0.277 | 0.269 | 0.017 | 0.958 | 0.920 | 0.277 |  |  |
|  |  | 2 | 0.137 | 0.160 | 0.170 | 0.156 | 0.017 | 1.097 |  |  |  |  |
|  |  | 3 | 0.144 | 0.179 | 0.147 | 0.157 | 0.019 | 0.748 |  |  |  |  |

|  |  |  | Renilla luciferase | | | | | Firefly luciferase | | | | |
| --- | --- | --- | --- | --- | --- | --- | --- | --- | --- | --- | --- | --- |
| **VEGFC IRES** |  | **Exp num** | **A** | **B** | **C** | **Mean** | **SD** | **A** | **B** | **C** | **Mean** | **SD** |
| normoxia | siCtrl | 1 | 1940200 | 1317594 | 1986534 | 1748109 | 373556 | 217056 | 137257 | 253247 | 202520 | 59346 |
|  |  | 2 | 1363206 | 1249203 | 1380695 | 1331034 | 71406 | 145917 | 137316 | 162033 | 148422 | 12547 |
|  |  | 3 | 448409 | 423742 | 553919 | 475357 | 69146 | 38149 | 39741 | 55704 | 44531 | 9709 |
|  | siPSPC1 | 1 | 2261492 | 2607191 | 2547477 | 2472053 | 184779 | 177855 | 262603 | 260378 | 233612 | 48300 |
|  |  | 2 | 1196580 | 1600569 | 1525023 | 1440724 | 214783 | 46171 | 85950 | 112355 | 81492 | 33316 |
|  |  | 3 | 552621 | 653237 | 669654 | 625170 | 63364 | 38782 | 47634 | 49944 | 45453 | 5891 |
| Hypoxia | siCtrl | 1 | 2035704 | 2389202 | 2247759 | 2224222 | 177921 | 281808 | 313489 | 279125 | 291474 | 19113 |
|  |  | 2 | 2763308 | 2109765 | 2969299 | 2614124 | 448766 | 128982 | 145698 | 226906 | 167195 | 52382 |
|  |  | 3 | 548838 | 425660 | 473070 | 482523 | 62131 | 48174 | 42953 | 55521 | 48883 | 6314 |
|  | siPSPC1 | 1 | 2904891 | 2858639 | 3277081 | 3013537 | 229404 | 328023 | 275547 | 293970 | 299180 | 26623 |
|  |  | 2 | 2975986 | 3429955 | 2396973 | 2934305 | 517751 | 131369 | 154116 | 146182 | 143889 | 11546 |
|  |  | 3 | 578088 | 560997 | 568108 | 569064 | 8586 | 50885 | 54865 | 50354 | 52035 | 2465 |

|  |  |  | Ratio LucF/LucR | | | | |  |  |  |  |  |
| --- | --- | --- | --- | --- | --- | --- | --- | --- | --- | --- | --- | --- |
|  |  | **Exp num** | **A** | **B** | **C** | **Mean** | **SD** | **Normalisation** | **Mean** | **SD** | **Mann-Whitney p-value** | **Significance** |
| normoxia | siCtrl | 1 | 0.112 | 0.104 | 0.127 | 0.115 | 0.012 | 1.000 | 1.000 | 0.118 | *0.0009* | **** |
|  |  | 2 | 0.107 | 0.110 | 0.117 | 0.111 | 0.005 | 1.000 |  |  |  |  |
|  |  | 3 | 0.085 | 0.094 | 0.101 | 0.093 | 0.008 | 1.000 |  |  |  |  |
|  | siPSPC1 | 1 | 0.079 | 0.101 | 0.102 | 0.094 | 0.013 | 0.820 | 0.695 | 0.188 |  |  |
|  |  | 2 | 0.039 | 0.054 | 0.074 | 0.055 | 0.018 | 0.496 |  |  |  |  |
|  |  | 3 | 0.070 | 0.073 | 0.075 | 0.073 | 0.002 | 0.779 |  |  |  |  |
| Hypoxia | siCtrl | 1 | 0.138 | 0.131 | 0.124 | 0.131 | 0.007 | 1.000 | 1.000 | 0.316 | *0.1983* | ns |
|  |  | 2 | 0.047 | 0.069 | 0.076 | 0.064 | 0.015 | 1.000 |  |  |  |  |
|  |  | 3 | 0.088 | 0.101 | 0.117 | 0.102 | 0.015 | 1.000 |  |  |  |  |
|  | siPSPC1 | 1 | 0.113 | 0.096 | 0.090 | 0.100 | 0.012 | 0.759 | 0.811 | 0.247 |  |  |
|  |  | 2 | 0.044 | 0.045 | 0.061 | 0.050 | 0.010 | 0.781 |  |  |  |  |
|  |  | 3 | 0.088 | 0.098 | 0.089 | 0.091 | 0.005 | 0.897 |  |  |  |  |

|  |  |  | Renilla luciferase | | | | | Firefly luciferase | | | | |
| --- | --- | --- | --- | --- | --- | --- | --- | --- | --- | --- | --- | --- |
| **VEGFD IRES** |  | **Exp num** | **A** | **B** | **C** | **Mean** | **SD** | **A** | **B** | **C** | **Mean** | **SD** |
| normoxia | siCtrl | 1 | 291674 | 287198 | 316877 | 298583 | 16000 | 19740 | 20522 | 20438 | 20233 | 429 |
|  |  | 2 | 567674 | 597668 | 541654 | 568999 | 28030 | 47796 | 48288 | 42436 | 46173 | 3246 |
|  |  | 3 | 546912 | 627624 | 776791 | 650442 | 116626 | 44748 | 51817 | 55327 | 50630 | 5388 |
|  | siPSPC1 | 1 | 352072 | 348952 | 328203 | 343076 | 12974 | 21662 | 18983 | 21014 | 20553 | 1398 |
|  |  | 2 | 552042 | 744632 | 688626 | 661767 | 99064 | 45207 | 53560 | 48649 | 49139 | 4198 |
|  |  | 3 | 781192 | 890044 | 980155 | 883797 | 99628 | 55585 | 54126 | 70395 | 60035 | 9001 |
| Hypoxia | siCtrl | 1 | 375774 | 397904 | 423655 | 399111 | 23963 | 40627 | 41173 | 42446 | 41415 | 934 |
|  |  | 2 | 525002 | 615275 | 500416 | 546898 | 60479 | 86899 | 103834 | 88966 | 93233 | 9239 |
|  |  | 3 | 878701 | 905268 | 897875 | 893948 | 13712 | 78430 | 97032 | 83797 | 86420 | 9574 |
|  | siPSPC1 | 1 | 428950 | 204448 | 344054 | 325817 | 113357 | 35543 | 28304 | 39439 | 34429 | 5650 |
|  |  | 2 | 561038 | 601141 | 684557 | 615578 | 63012 | 90108 | 100950 | 111935 | 100998 | 10913 |
|  |  | 3 | 1131210 | 1211175 | 1365269 | 1235884 | 118970 | 76199 | 94973 | 100224 | 90465 | 12631 |

|  |  |  | Ratio LucF/LucR | | | | |  |  |  |  |  |
| --- | --- | --- | --- | --- | --- | --- | --- | --- | --- | --- | --- | --- |
|  |  | **Exp num** | **A** | **B** | **C** | **Mean** | **SD** | **Normalisation** | **Mean** | **SD** | **Mann-Whitney p-value** | **Significance** |
| normoxia | siCtrl | 1 | 0.068 | 0.071 | 0.064 | 0.068 | 0.003 | 1.000 | 1.000 | 0.095 | *0.09* | ns |
|  |  | 2 | 0.084 | 0.081 | 0.078 | 0.081 | 0.003 | 1.000 |  |  |  |  |
|  |  | 3 | 0.082 | 0.083 | 0.071 | 0.079 | 0.006 | 1.000 |  |  |  |  |
|  | siPSPC1 | 1 | 0.062 | 0.054 | 0.064 | 0.060 | 0.005 | 0.884 | 0.891 | 0.108 |  |  |
|  |  | 2 | 0.082 | 0.072 | 0.071 | 0.075 | 0.006 | 0.922 |  |  |  |  |
|  |  | 3 | 0.071 | 0.061 | 0.072 | 0.068 | 0.006 | 0.865 |  |  |  |  |
| Hypoxia | siCtrl | 1 | 0.108 | 0.103 | 0.100 | 0.104 | 0.004 | 1.000 | 1.000 | 0.290 | *0.4283* | ns |
|  |  | 2 | 0.166 | 0.169 | 0.178 | 0.171 | 0.006 | 1.000 |  |  |  |  |
|  |  | 3 | 0.089 | 0.107 | 0.093 | 0.097 | 0.009 | 1.000 |  |  |  |  |
|  | siPSPC1 | 1 | 0.083 | 0.138 | 0.115 | 0.112 | 0.028 | 1.077 | 0.940 | 0.340 |  |  |
|  |  | 2 | 0.161 | 0.168 | 0.164 | 0.164 | 0.004 | 0.961 |  |  |  |  |
|  |  | 3 | 0.067 | 0.078 | 0.073 | 0.073 | 0.006 | 0.756 |  |  |  |  |

|  |  |  | Renilla luciferase | | | | | Firefly luciferase | | | | |
| --- | --- | --- | --- | --- | --- | --- | --- | --- | --- | --- | --- | --- |
| **IGF1R IRES** |  | **Exp num** | **A** | **B** | **C** | **Mean** | **SD** | **A** | **B** | **C** | **Mean** | **SD** |
| normoxia | siCtrl | 1 | 722018 | 972960 | 906771 | 867250 | 130055 | 159594 | 232929 | 214092 | 202205 | 38085 |
|  |  | 2 | 860625 | 880028 | 795415 | 845356 | 44325 | 202188 | 218830 | 189480 | 203499 | 14719 |
|  |  | 3 | 255284 | 263849 | 293168 | 270767 | 19867 | 103285 | 90152 | 119297 | 104245 | 14596 |
|  | siPSPC1 | 1 | 1060761 | 1289052 | 1353833 | 1234548 | 153951 | 155151 | 213263 | 201468 | 189961 | 30718 |
|  |  | 2 | 985309 | 1048775 | 905864 | 979983 | 71604 | 167280 | 199053 | 135802 | 167379 | 31625 |
|  |  | 3 | 427098 | 484160 | 423908 | 445055 | 33903 | 121673 | 123232 | 117490 | 120798 | 2969 |
| Hypoxia | siCtrl | 1 | 1058689 | 1125486 | 1292036 | 1158737 | 120175 | 304929 | 348329 | 355248 | 336169 | 27275 |
|  |  | 2 | 1158913 | 1300106 | 1178171 | 1212396 | 76567 | 212389 | 306379 | 271984 | 263584 | 47555 |
|  |  | 3 | 210319 | 236396 | 346932 | 264549 | 72527 | 98731 | 90687 | 136145 | 108521 | 24259 |
|  | siPSPC1 | 1 | 1597529 | 1714796 | 1755494 | 1689273 | 82017 | 258925 | 386756 | 318222 | 321301 | 63971 |
|  |  | 2 | 1594579 | 1752745 | 1758274 | 1701866 | 92954 | 276042 | 309622 | 288385 | 291350 | 16985 |
|  |  | 3 | 114079 | 278087 | 178890 | 190352 | 82603 | 71140 | 96693 | 81616 | 83150 | 12845 |

|  |  |  | Ratio LucF/LucR | | | | |  |  |  |  |  |
| --- | --- | --- | --- | --- | --- | --- | --- | --- | --- | --- | --- | --- |
|  |  | **Exp num** | **A** | **B** | **C** | **Mean** | **SD** | **Normalisation** | **Mean** | **SD** | **Mann-Whitney p-value** | **Significance** |
| normoxia | siCtrl | 1 | 0.221 | 0.239 | 0.236 | 0.232 | 0.010 | 1.000 | 1.000 | 0.268 | *0.0502* | ns |
|  |  | 2 | 0.235 | 0.249 | 0.238 | 0.241 | 0.007 | 1.000 |  |  |  |  |
|  |  | 3 | 0.405 | 0.342 | 0.407 | 0.384 | 0.037 | 1.000 |  |  |  |  |
|  | siPSPC1 | 1 | 0.146 | 0.165 | 0.149 | 0.154 | 0.010 | 0.661 | 0.695 | 0.201 |  |  |
|  |  | 2 | 0.170 | 0.190 | 0.150 | 0.170 | 0.020 | 0.706 |  |  |  |  |
|  |  | 3 | 0.285 | 0.255 | 0.277 | 0.272 | 0.016 | 0.708 |  |  |  |  |
| Hypoxia | siCtrl | 1 | 0.288 | 0.309 | 0.275 | 0.291 | 0.017 | 1.000 | 1.000 | 0.298 | *0.16* | ns |
|  |  | 2 | 0.183 | 0.236 | 0.231 | 0.217 | 0.029 | 1.000 |  |  |  |  |
|  |  | 3 | 0.469 | 0.384 | 0.392 | 0.415 | 0.047 | 1.000 |  |  |  |  |
|  | siPSPC1 | 1 | 0.162 | 0.226 | 0.181 | 0.190 | 0.033 | 0.652 | 0.907 | 0.534 |  |  |
|  |  | 2 | 0.173 | 0.177 | 0.164 | 0.171 | 0.007 | 0.791 |  |  |  |  |
|  |  | 3 | 0.624 | 0.348 | 0.456 | 0.476 | 0.139 | 1.146 |  |  |  |  |

|  |  |  | Renilla luciferase | | | | | Firefly luciferase | | | | |
| --- | --- | --- | --- | --- | --- | --- | --- | --- | --- | --- | --- | --- |
| **c-myc IRES** |  | **Exp num** | **A** | **B** | **C** | **Mean** | **SD** | **A** | **B** | **C** | **Mean** | **SD** |
| normoxia | siCtrl | 1 | 1376489 | 1656411 | 1634471 | 1555790 | 155667 | 241155 | 349763 | 295138 | 295352 | 54305 |
|  |  | 2 | 1216742 | 1371688 | 1207546 | 1265325 | 92227 | 325307 | 384619 | 326642 | 345523 | 33865 |
|  |  | 3 | 936981 | 912821 | 923416 | 924406 | 12111 | 353453 | 426897 | 451667 | 410672 | 51078 |
|  | siPSPC1 | 1 | 1959245 | 1729456 | 1762524 | 1817075 | 124228 | 332133 | 310274 | 335337 | 325915 | 13640 |
|  |  | 2 | 1430337 | 1754454 | 1669279 | 1618023 | 168027 | 322390 | 411266 | 351022 | 361559 | 45365 |
|  |  | 3 | 1297955 | 1402948 | 1494146 | 1398350 | 98176 | 494757 | 486234 | 539230 | 506740 | 28458 |
| Hypoxia | siCtrl | 1 | 2303679 | 2473031 | 2051451 | 2276054 | 212143 | 531197 | 646002 | 466514 | 547904 | 90903 |
|  |  | 2 | 1405500 | 1480296 | 1467212 | 1451002 | 39946 | 575691 | 601917 | 534370 | 570659 | 34053 |
|  |  | 3 | 802990 | 789755 | 806209 | 799651 | 8720 | 392506 | 481930 | 399124 | 424520 | 49828 |
|  | siPSPC1 | 1 | 2136726 | 973440 | 2184793 | 1764986 | 685921 | 411768 | 329971 | 419752 | 387164 | 49691 |
|  |  | 2 | 1802730 | 2187577 | 2061130 | 2017146 | 196157 | 587572 | 539105 | 606345 | 577674 | 34696 |
|  |  | 3 | 711911 | 903770 | 638287 | 751323 | 137060 | 355455 | 327717 | 305612 | 329595 | 24974 |

|  |  |  | Ratio LucF/LucR | | | | |  |  |  |  |  |
| --- | --- | --- | --- | --- | --- | --- | --- | --- | --- | --- | --- | --- |
|  |  | **Exp num** | **A** | **B** | **C** | **Mean** | **SD** | **Normalisation** | **Mean** | **SD** | **Mann-Whitney p-value** | **Significance** |
| normoxia | siCtrl | 1 | 0.175 | 0.211 | 0.181 | 0.189 | 0.019 | 1.000 | 1.000 | 0.388 | *0.3799* | ns |
|  |  | 2 | 0.267 | 0.280 | 0.271 | 0.273 | 0.007 | 1.000 |  |  |  |  |
|  |  | 3 | 0.377 | 0.468 | 0.489 | 0.445 | 0.059 | 1.000 |  |  |  |  |
|  | siPSPC1 | 1 | 0.170 | 0.179 | 0.190 | 0.180 | 0.010 | 0.951 | 0.845 | 0.277 |  |  |
|  |  | 2 | 0.225 | 0.234 | 0.210 | 0.223 | 0.012 | 0.819 |  |  |  |  |
|  |  | 3 | 0.381 | 0.347 | 0.361 | 0.363 | 0.017 | 0.816 |  |  |  |  |
| Hypoxia | siCtrl | 1 | 0.231 | 0.261 | 0.227 | 0.240 | 0.019 | 1.000 | 1.000 | 0.340 | *0.293* | ns |
|  |  | 2 | 0.410 | 0.407 | 0.364 | 0.393 | 0.025 | 1.000 |  |  |  |  |
|  |  | 3 | 0.489 | 0.610 | 0.495 | 0.531 | 0.068 | 1.000 |  |  |  |  |
|  | siPSPC1 | 1 | 0.193 | 0.339 | 0.192 | 0.241 | 0.085 | 1.006 | 0.839 | 0.285 |  |  |
|  |  | 2 | 0.326 | 0.246 | 0.294 | 0.289 | 0.040 | 0.734 |  |  |  |  |
|  |  | 3 | 0.499 | 0.363 | 0.479 | 0.447 | 0.074 | 0.841 |  |  |  |  |

|  |  |  | Renilla luciferase | | | | | Firefly luciferase | | | | |
| --- | --- | --- | --- | --- | --- | --- | --- | --- | --- | --- | --- | --- |
| **EMCV IRES** |  | **Exp num** | **A** | **B** | **C** | **Mean** | **SD** | **A** | **B** | **C** | **Mean** | **SD** |
| normoxia | siCtrl | 1 | 147772 | 179110 | 161146 | 162676 | 15725 | 107301 | 144501 | 143440 | 131747 | 21177 |
|  |  | 2 | 193872 | 193384 | 139866 | 175707 | 31040 | 159216 | 163871 | 110780 | 144622 | 29400 |
|  |  | 3 | 4196611 | 4747196 | 4813952 | 4585920 | 338799 | 3995099 | 4448880 | 4971984 | 4471988 | 488853 |
|  | siPSPC1 | 1 | 154112 | 256024 | 273750 | 227962 | 64567 | 128725 | 193675 | 239512 | 187304 | 55667 |
|  |  | 2 | 280515 | 360485 | 320666 | 320555 | 39985 | 233797 | 293037 | 235866 | 254233 | 33621 |
|  |  | 3 | 3779698 | 5201252 | 5024431 | 4668460 | 774751 | 3165164 | 4909784 | 4212893 | 4095947 | 717023 |
| Hypoxia | siCtrl | 1 | 177395 | 182885 | 198846 | 186375 | 11143 | 223538 | 216316 | 233362 | 224405 | 8556 |
|  |  | 2 | 289545 | 319434 | 335655 | 314878 | 23390 | 319218 | 329348 | 326687 | 325084 | 5251 |
|  |  | 3 | 8950028 | 9989663 | 7196947 | 8712213 | 1411465 | 7490989 | 18294662 | 14542755 | 13442802 | 5485186 |
|  | siPSPC1 | 1 | 255210 | 255153 | 241881 | 250748 | 7679 | 313768 | 293517 | 256449 | 287912 | 29068 |
|  |  | 2 | 515069 | 443759 | 489733 | 482854 | 36150 | 531615 | 444622 | 522848 | 499695 | 47896 |
|  |  | 3 | 9321714 | 9048404 | 9141799 | 9170639 | 138918 | 13646640 | 13487616 | 14622621 | 13918959 | 614555 |

|  |  |  | Ratio LucF/LucR | | | | |  |  |  |  |  |
| --- | --- | --- | --- | --- | --- | --- | --- | --- | --- | --- | --- | --- |
|  |  | **Exp num** | **A** | **B** | **C** | **Mean** | **SD** | **Normalisation** | **Mean** | **SD** | **Mann-Whitney p-value** | **Significance** |
| normoxia | siCtrl | 1 | 0.726 | 0.807 | 0.890 | 0.808 | 0.082 | 1.000 | 1.000 | 0.109 | *0.5346* | ns |
|  |  | 2 | 0.821 | 0.847 | 0.792 | 0.820 | 0.028 | 1.000 |  |  |  |  |
|  |  | 3 | 0.952 | 0.937 | 1.033 | 0.974 | 0.051 | 1.000 |  |  |  |  |
|  | siPSPC1 | 1 | 0.835 | 0.756 | 0.875 | 0.822 | 0.060 | 1.018 | 0.957 | 0.070 |  |  |
|  |  | 2 | 0.833 | 0.813 | 0.736 | 0.794 | 0.052 | 0.968 |  |  |  |  |
|  |  | 3 | 0.837 | 0.944 | 0.838 | 0.873 | 0.061 | 0.897 |  |  |  |  |
| Hypoxia | siCtrl | 1 | 1.260 | 1.183 | 1.174 | 1.205 | 0.048 | 1.000 | 1.000 | 0.313 | *0.909* | ns |
|  |  | 2 | 1.102 | 1.031 | 0.973 | 1.036 | 0.065 | 1.000 |  |  |  |  |
|  |  | 3 | 0.837 | 1.831 | 2.021 | 1.563 | 0.636 | 1.000 |  |  |  |  |
|  | siPSPC1 | 1 | 1.229 | 1.150 | 1.060 | 1.147 | 0.085 | 0.951 | 0.972 | 0.179 |  |  |
|  |  | 2 | 1.032 | 1.002 | 1.068 | 1.034 | 0.033 | 0.998 |  |  |  |  |
|  |  | 3 | 1.464 | 1.491 | 1.600 | 1.518 | 0.072 | 0.971 |  |  |  |  |

**Supplementary file 6. IRES activities in normoxic and hypoxic HL-1 cells after PSPC1 knock-down.**

HL-1 cells were transduced with Lucky Luke bicistronic lentivectors containing the IRES of FGF1, FGF2, VEGFA (a or b), VEGFC, VEGFD, IGF1R, c-myc or EMCV. Cells were then treated with siPSPC1 or siControl (siCtrl) smartpool during normoxia or hypoxia 1% O_2._ Renilla and firefly luciferase activities were measured (upper panel, or page 1 for FGF1 IRES) and the IRES activities evaluated with the ratio LucF/LucR (lower panel, or page 2 for FGF1 IRES). Mann-Whitney test was performed with n=9 (n=12 for FGF1 IRES). *p<0.05, **p<0.01, ***<0.001, ****p<0.0001. For each IRES the mean has been calculated with nine cell culture biological replicates (Experiments A, B, C correspond to experiments performed at different dates, while 1, 2, 3 are experiments performed in parallel at the same date, each of them being already the mean of three technical replicates (27 technical replicates in total).
