## Supplementary files 1-8 for "Long non-coding RNA Neat1 and paraspeckle components are translational regulators in hypoxia": V2 Supplementary file 8 Neat1 KD fluidigm .docx

| **A/ Neat1 knock-down** | | |  |  |  |  |  |  |  |
| --- | --- | --- | --- | --- | --- | --- | --- | --- | --- |
|  | **Total mRNA** | | | **Polysome bound mRNA** | | **Fold change** | | **Fold change Mean** | |
| **Gene name** | RQ 1 | | RQ 2 | RQ 1 | RQ 2 | RQ (polysomes)/ RQ (Total mRNA) 1 | RQ (polysomes)/ RQ (Total mRNA) 2 | RQ (ou -1/RQ) | SD |
| *Aanat* | ND | | 87.95 | 1.58 | ND | ND | ND | ND | ND |
| *Adar (Adar1)* | 1.35 | | 0.88 | 1.14 | 0.99 | 0.85 | 1.12 | -1.02 | 0.14 |
| *Alox15* | ND | | 0.22 | 4.17 | 0.61 | ND | 2.75 | ND | ND |
| *Ang* | 0.80 | | 1.00 | 0.50 | 0.82 | 0.63 | 0.82 | -1.38 | 0.09 |
| *Apaf1* | 1.21 | | 0.93 | 1.50 | 1.17 | 1.24 | 1.25 | 1.25 | 0.01 |
| *Apln* | 1.36 | | 0.99 | 1.07 | 1.23 | 0.79 | 1.24 | 1.02 | 0.22 |
| *Aplnr* | 1.15 | | 0.22 | 0.78 | 0.20 | 0.68 | 0.92 | -1.25 | 0.12 |
| *Bag1* | 1.19 | | 1.03 | 0.75 | 1.12 | 0.63 | 1.09 | -1.16 | 0.23 |
| *Bax* | 1.27 | | 1.05 | 1.00 | 1.18 | 0.78 | 1.12 | -1.05 | 0.17 |
| *Bcl2 (Bcl-2)* | 1.02 | | 0.96 | 0.56 | 0.63 | 0.55 | 0.65 | -1.66 | 0.05 |
| *Bcl2L1 (Bcl-XL)* | 1.25 | | 1.09 | 1.42 | 0.94 | 1.13 | 0.86 | -1.00 | 0.14 |
| *Bip(GRP78/Hspa5)* | 1.18 | | 0.64 | 0.95 | 0.92 | 0.80 | 1.45 | 1.13 | 0.33 |
| *BNP (Nppb)* | 3.36 | | 0.93 | 1.45 | 1.88 | 0.43 | 2.04 | 1.23 | 0.80 |
| *Casp2 (Caspase-2)* | 0.95 | | 0.42 | 1.08 | 0.82 | 1.14 | 1.94 | 1.54 | 0.40 |
| *Cdk1* | 0.97 | | 0.72 | 0.78 | 0.88 | 0.80 | 1.23 | 1.01 | 0.21 |
| *Cdkn1B (p27kip1)* | 0.87 | | 0.86 | 0.75 | 0.89 | 0.87 | 1.03 | -1.05 | 0.08 |
| *Cdkn2A (p16INK4)* | 1.08 | | 1.01 | 0.69 | 0.90 | 0.64 | 0.89 | -1.31 | 0.13 |
| *Col1A1* | 1.06 | | 0.69 | 0.95 | 0.73 | 0.90 | 1.06 | -1.02 | 0.08 |
| *Col3A1* | 0.80 | | 0.54 | 1.13 | 0.55 | 1.41 | 1.01 | 1.21 | 0.20 |
| *Csde1* | 1.07 | | 0.72 | 0.84 | 0.89 | 0.78 | 1.23 | 1.01 | 0.23 |
| *Ctgf* | 1.02 | | 0.79 | 1.05 | 1.09 | 1.03 | 1.39 | 1.21 | 0.18 |
| *Cyr61* | 1.60 | | 1.37 | 1.78 | 1.34 | 1.11 | 0.97 | 1.04 | 0.07 |
| *Dazap1* | 0.97 | | 1.11 | 0.71 | 0.88 | 0.74 | 0.79 | -1.31 | 0.02 |
| *Ddx17* | 1.11 | | 0.65 | 1.15 | 0.65 | 1.04 | 1.00 | 1.02 | 0.02 |
| *Egr1* | 2.50 | | 1.36 | 1.90 | 1.08 | 0.76 | 0.79 | -1.29 | 0.01 |
| *Eno3* | 1.12 | | 0.78 | 0.92 | 0.81 | 0.83 | 1.04 | -1.07 | 0.10 |
| *Epas1* | 1.53 | | 0.84 | 0.89 | 0.94 | 0.58 | 1.12 | -1.17 | 0.27 |
| *Fgf1* | 0.85 | | 0.51 | 0.81 | 0.69 | 0.95 | 1.36 | 1.16 | 0.20 |
| *Fgf10* | 1.53 | | 0.78 | 1.12 | 1.20 | 0.73 | 1.53 | 1.13 | 0.40 |
| *Fgf2* | ND | | 1.23 | 0.19 | 0.23 | ND | 0.19 | ND | ND |
| *Fgf21* | 1.33 | | 0.93 | 0.49 | 0.44 | 0.37 | 0.47 | -2.39 | 0.05 |
| *Fgf23* | ND | | ND | 2.38 | ND | ND | ND | ND | ND |
| *Fgf4* | 0.23 | | 193.41 | 32.32 | 8.24 | 139.51 | 0.04 | ND | ND |
| *Fgfr1* | 0.94 | | 0.51 | 0.58 | 0.48 | 0.62 | 0.94 | -1.28 | 0.16 |
| *Fgfr2* | 1.10 | | 0.78 | 0.87 | 0.67 | 0.79 | 0.87 | -1.21 | 0.04 |
| *Fgfr3* | 1.20 | | 0.76 | 1.07 | 1.04 | 0.89 | 1.38 | 1.14 | 0.24 |
| *Fmr1 (Fmrp)* | 1.21 | | 1.07 | 0.63 | 1.02 | 0.52 | 0.95 | -1.36 | 0.22 |
| *Fn1 (Fibronectine)* | 0.88 | | 0.73 | 1.03 | 0.69 | 1.18 | 0.95 | 1.06 | 0.12 |
| *Fus* | 1.04 | | 0.66 | 0.99 | 0.71 | 0.96 | 1.07 | 1.01 | 0.06 |
| *Gapdh* | 1.00 | | 1.00 | 1.00 | 1.00 | 1.00 | 1.00 | -1.00 | 0.00 |
| *Gusb* | 1.11 | | 0.72 | 1.19 | 1.00 | 1.07 | 1.39 | 1.23 | 0.16 |
| *Hgf* | ND | | ND | 0.37 | ND | ND | ND | ND | ND |
| *Hif1A* | 0.91 | | 0.73 | 0.98 | 0.79 | 1.08 | 1.08 | 1.08 | 0.00 |
| *Hnrnpa1* | 1.01 | | 0.67 | 0.83 | 0.69 | 0.83 | 1.02 | -1.09 | 0.09 |
| *Hnrnph3* | 1.10 | | 1.25 | 0.55 | 0.99 | 0.50 | 0.79 | -1.55 | 0.14 |
| *Hnrnpk* | 0.99 | | 0.90 | 0.91 | 0.88 | 0.92 | 0.98 | -1.06 | 0.03 |
| *Hnrnpm* | 1.02 | | 1.03 | 0.67 | 0.97 | 0.66 | 0.93 | -1.25 | 0.14 |
| *Hnrnpr* | 1.16 | | 0.90 | 0.69 | 1.02 | 0.59 | 1.13 | -1.16 | 0.27 |
| *Hoxa9* | 1.08 | | 2.35 | 9.51 | 1.41 | 8.83 | 0.60 | ND | ND |
| *HuR (Elavl1)* | 0.88 | | 0.76 | 1.06 | 0.83 | 1.20 | 1.09 | 1.15 | 0.05 |
| *Igf1* | 1.11 | | 0.92 | 0.74 | 0.69 | 0.67 | 0.75 | -1.41 | 0.04 |
| *Igf1R* | 0.92 | | 0.87 | 0.90 | 0.68 | 0.98 | 0.79 | -1.13 | 0.10 |
| *Igf2R* | 0.93 | | 0.74 | 1.08 | 0.88 | 1.16 | 1.19 | 1.17 | 0.02 |
| *Irf2* | 1.18 | | 1.05 | 0.95 | 0.95 | 0.80 | 0.91 | -1.17 | 0.05 |
| *Lamb1* | 0.74 | | 1.03 | 0.44 | 0.77 | 0.60 | 0.75 | -1.49 | 0.07 |
| *Lef1* | ND | | ND | 0.41 | 0.55 | ND | ND | ND | ND |
| *Mmp2* | 1.01 | | 0.92 | 1.84 | 0.86 | 1.83 | 0.94 | 1.39 | 0.44 |
| *Mycbp* | 0.91 | | 0.89 | 0.69 | 0.99 | 0.75 | 1.11 | -1.07 | 0.18 |
| *Mycl* | 1.93 | | 1.84 | 2.39 | 1.52 | 1.24 | 0.83 | 1.03 | 0.21 |
| *Ncl* | 1.12 | | 1.10 | 0.49 | 0.93 | 0.44 | 0.84 | -1.56 | 0.20 |
| *Neat1* | 0.54 | | 0.81 | 0.43 | 0.56 | 0.80 | 0.68 | -1.35 | 0.06 |
| *Neat1-2* | 0.61 | | 0.50 | 0.54 | 0.37 | 0.89 | 0.75 | -1.22 | 0.07 |
| *Nfil3* | 0.80 | | 1.08 | 1.16 | 0.86 | 1.45 | 0.80 | 1.12 | 0.33 |
| *Nkrf (NRF)* | 0.97 | | 1.25 | 0.84 | 1.08 | 0.87 | 0.86 | -1.15 | 0.01 |
| *Nono* | 1.21 | | 1.26 | 0.87 | 0.92 | 0.72 | 0.73 | -1.38 | 0.01 |
| *Nr1D1 (rev-erb-a)* | 1.23 | | 1.09 | 0.86 | 1.04 | 0.70 | 0.95 | -1.21 | 0.13 |
| *Pdgfa* | 1.12 | | 1.03 | 0.76 | 1.03 | 0.68 | 1.00 | -1.19 | 0.16 |
| *Pdgfb* | 1.07 | | 0.89 | 1.18 | 1.07 | 1.11 | 1.20 | 1.15 | 0.05 |
| *Per1 (Period-1)* | 1.18 | | 1.13 | 1.21 | 1.03 | 1.03 | 0.92 | -1.03 | 0.05 |
| *Pfkm (Pfk1)* | 0.81 | | 0.63 | 0.97 | 0.57 | 1.20 | 0.90 | 1.05 | 0.15 |
| *Pgf (Plgf))* | 1.02 | | 1.00 | 1.10 | 1.77 | 1.09 | 1.76 | 1.42 | 0.34 |
| *Prox1* | 0.85 | | 0.94 | 0.88 | 0.97 | 1.04 | 1.03 | 1.03 | 0.01 |
| *Pspc1* | 1.09 | | 1.05 | 0.65 | 0.93 | 0.59 | 0.89 | -1.35 | 0.15 |
| *Rbm14* | 1.11 | | 0.94 | 1.03 | 0.87 | 0.93 | 0.93 | -1.07 | 0.00 |
| *Rpl10A* | 0.91 | | 0.82 | 1.26 | 1.09 | 1.39 | 1.33 | 1.36 | 0.03 |
| *Rps2* | 1.12 | | 0.71 | 1.23 | 0.86 | 1.10 | 1.22 | 1.16 | 0.06 |
| *Rps25* | 1.12 | | 1.09 | 0.87 | 0.99 | 0.78 | 0.91 | -1.19 | 0.07 |
| *Rrbp1* | 0.94 | | 0.68 | 0.58 | 0.55 | 0.62 | 0.82 | -1.39 | 0.10 |
| *Serpine1 (PAI1)* | 1.03 | | 0.62 | 0.88 | 0.72 | 0.86 | 1.15 | 1.00 | 0.15 |
| *Setd7 (Set7)* | 1.23 | | 0.90 | 1.47 | 0.83 | 1.20 | 0.92 | 1.06 | 0.14 |
| *Sfpq* | 0.99 | | 1.07 | 0.71 | 0.89 | 0.72 | 0.83 | -1.29 | 0.05 |
| *Shmt1* | 1.14 | | 0.81 | 0.98 | 0.86 | 0.86 | 1.06 | -1.04 | 0.10 |
| *Slc7A1 (Cat-1)* | 1.07 | | 0.66 | 1.06 | 0.79 | 0.99 | 1.21 | 1.10 | 0.11 |
| *Srebf1 (Srebp1)* | 0.75 | | 0.70 | 0.84 | 0.78 | 1.12 | 1.11 | 1.12 | 0.00 |
| *Sstr2 (Sst2)* | 1.55 | | 1.51 | 3.86 | 1.94 | 2.49 | 1.28 | 1.89 | 0.60 |
| *Thbd (Thrombomodulin)* | 1.85 | | 0.91 | 1.05 | 0.96 | 0.57 | 1.06 | -1.23 | 0.24 |
| *Trp53* | 1.28 | | 1.14 | 1.08 | 1.09 | 0.84 | 0.95 | -1.11 | 0.06 |
| *Txnip* | 1.13 | | 0.74 | 1.22 | 0.86 | 1.07 | 1.15 | 1.11 | 0.04 |
| *Utrn (Utrophin)* | 1.04 | | 0.83 | 0.59 | 0.63 | 0.57 | 0.76 | -1.50 | 0.09 |
| *Vash1* | 1.06 | | 0.68 | 1.13 | 0.96 | 1.07 | 1.41 | 1.24 | 0.17 |
| *Vegfa* | 0.87 | | 0.95 | 0.83 | 1.02 | 0.96 | 1.07 | 1.01 | 0.06 |
| *Vegfb* | 0.93 | | 0.90 | 0.75 | 0.90 | 0.81 | 1.00 | -1.11 | 0.09 |
| *Vegfc* | ND | | ND | 1.16 | ND | ND | ND | ND | ND |
| *Vegfd (Figf)* | 0.84 | | 0.66 | 0.68 | 0.99 | 0.81 | 1.50 | 1.15 | 0.34 |
| *Xiap* | 0.97 | | 0.79 | 0.97 | 0.96 | 1.00 | 1.20 | 1.10 | 0.10 |

| **B/ Neat1_2 knock-down** |
| --- |

|  | **Total mRNA** | | **Polysome bound mRNA** | | **Fold change** | | **Fold change mean** | |
| --- | --- | --- | --- | --- | --- | --- | --- | --- |
| **Gene name** | RQ 1 | RQ 2 | RQ 1 | RQ 2 | RQ polysomes/ RQ Total mRNA 1 | RQ polysomes/ RQ Total mRNA 2 | RQ (or -1/RQ) | SD |
| *Aanat* | ND | 306.15 | ND | 0.73 | ND | 0.00 | ND | ND |
| *Adar (Adar1)* | 1.10 | 1.34 | 0.88 | 1.16 | 0.80 | 0.87 | -1.20 | 0.03 |
| *Alox15* | 0.45 | 2.70 | 1.98 | 0.36 | 4.37 | 0.13 | ND | ND |
| *Ang* | 0.83 | 1.49 | 0.86 | 1.09 | 1.04 | 0.73 | -1.13 | 0.16 |
| *Apaf1* | 1.04 | 1.16 | 0.93 | 1.04 | 0.90 | 0.90 | -1.11 | 0.00 |
| *Apln* | 1.23 | 1.53 | 0.76 | 1.54 | 0.62 | 1.00 | -1.23 | 0.19 |
| *Aplnr* | ND | 3.03 | 0.97 | 1.74 | ND | 0.57 | ND | ND |
| *Bag1* | 1.28 | 1.38 | 1.27 | 1.04 | 0.99 | 0.76 | -1.15 | 0.12 |
| *Bax* | 1.15 | 1.00 | 0.79 | 1.00 | 0.69 | 1.00 | -1.18 | 0.16 |
| *Bcl2 (Bcl-2)* | 1.04 | 1.38 | 0.91 | 1.16 | 0.88 | 0.84 | -1.16 | 0.02 |
| *Bcl2L1 (Bcl-XL)* | 1.19 | 1.20 | 0.96 | 1.12 | 0.81 | 0.93 | -1.15 | 0.06 |
| *Bip (GRP78/Hspa5)* | 0.78 | 1.16 | 0.67 | 0.92 | 0.87 | 0.79 | -1.21 | 0.04 |
| *BNP (Nppb)* | 0.83 | 0.60 | 0.61 | 0.88 | 0.74 | 1.48 | 1.11 | 0.37 |
| *Casp2 (Caspase-2)* | 1.02 | 1.30 | 0.99 | 0.99 | 0.97 | 0.76 | -1.15 | 0.10 |
| *Cdk1* | 1.10 | 1.19 | 1.00 | 1.08 | 0.91 | 0.91 | -1.10 | 0.00 |
| *Col1A1* | 1.52 | 1.35 | 0.76 | 1.00 | 0.50 | 0.74 | -1.61 | 0.12 |
| *Col3A1* | 1.47 | 1.08 | 0.38 | 1.05 | 0.25 | 0.98 | -1.62 | 0.36 |
| *Csde1* | 1.21 | 1.48 | 1.18 | 0.99 | 0.98 | 0.67 | -1.22 | 0.15 |
| *Ctgf* | 1.09 | 1.47 | 0.74 | 0.99 | 0.68 | 0.67 | -1.48 | 0.00 |
| *Cyr61* | 1.08 | 1.00 | 0.73 | 1.02 | 0.68 | 1.03 | -1.17 | 0.17 |
| *Dazap1* | 1.15 | 1.35 | 1.05 | 0.99 | 0.91 | 0.73 | -1.22 | 0.09 |
| *Ddx17* | 1.12 | 1.22 | 0.85 | 1.17 | 0.76 | 0.95 | -1.17 | 0.10 |
| *Egr1* | 0.96 | 1.16 | 0.80 | 0.90 | 0.83 | 0.78 | -1.24 | 0.03 |
| *Eno3* | 1.03 | 1.36 | 0.93 | 1.08 | 0.90 | 0.80 | -1.18 | 0.05 |
| *Epas1* | 1.17 | 1.58 | 0.95 | 1.02 | 0.82 | 0.64 | -1.37 | 0.09 |
| *Fgf1* | 1.09 | 1.45 | 1.00 | 0.95 | 0.92 | 0.65 | -1.27 | 0.13 |
| *Fgf10* | 0.90 | 1.26 | 0.25 | 2.94 | 0.27 | 2.33 | ND | ND |
| *Fgf2* | 4.34 | 2.07 | 0.85 | 2.41 | 0.20 | 1.16 | ND | ND |
| *Fgf21* | 0.82 | 0.97 | 0.67 | 1.01 | 0.82 | 1.03 | -1.08 | 0.10 |
| *Fgf23* | ND | ND | ND | 0.86 | ND | ND | ND | ND |
| *Fgf4* | 10.86 | 1.25 | 1.93 | 1.48 | 0.18 | 1.18 | -1.47 | 0.50 |
| *Fgfr1* | 1.19 | 1.86 | 0.66 | 1.04 | 0.56 | 0.56 | -1.79 | 0.00 |
| *Fgfr2* | 0.70 | 0.79 | 1.13 | 1.44 | 1.61 | 1.83 | 1.72 | 0.11 |
| *Fgfr3* | 0.96 | 1.39 | 0.77 | 1.06 | 0.80 | 0.77 | -1.28 | 0.02 |
| *Fmr1 (Fmrp)* | 1.30 | 1.67 | 1.17 | 1.31 | 0.90 | 0.78 | -1.19 | 0.06 |
| *Fn1 (Fibronectine)* | 0.96 | 1.41 | 0.86 | 1.46 | 0.89 | 1.04 | -1.04 | 0.07 |
| *Fus* | 0.99 | 1.26 | 0.89 | 1.09 | 0.90 | 0.86 | -1.14 | 0.02 |
| *Gapdh* | 1.00 | 1.00 | 1.00 | 1.00 | 1.00 | 1.00 | -1.00 | 0.00 |
| *Gusb* | 1.09 | 1.26 | 0.88 | 1.13 | 0.81 | 0.89 | -1.17 | 0.04 |
| *Hgf* | ND | ND | ND | ND | ND | ND | ND | ND |
| *Hif1A* | 0.99 | 1.18 | 1.01 | 1.03 | 1.02 | 0.87 | -1.06 | 0.08 |
| *Hnrnpa1* | 1.10 | 1.14 | 0.96 | 1.07 | 0.87 | 0.94 | -1.10 | 0.04 |
| *Hnrnph3* | 1.01 | 1.59 | 0.85 | 1.14 | 0.85 | 0.72 | -1.28 | 0.06 |
| *Hnrnpk* | 1.11 | 1.03 | 0.94 | 1.06 | 0.85 | 1.02 | -1.07 | 0.09 |
| *Hnrnpm* | 1.04 | 1.50 | 1.02 | 1.21 | 0.98 | 0.81 | -1.12 | 0.08 |
| *Hnrnpr* | 1.22 | 1.44 | 1.11 | 1.16 | 0.91 | 0.81 | -1.16 | 0.05 |
| *Hoxa9* | 0.65 | 0.60 | 0.52 | 0.19 | 0.80 | 0.31 | -1.80 | 0.24 |
| *HuR (Elavl1)* | 1.04 | 1.35 | 0.97 | 0.99 | 0.93 | 0.73 | -1.20 | 0.10 |
| *Igf1* | 1.98 | 2.38 | 0.54 | 1.10 | 0.27 | 0.46 | -2.72 | 0.10 |
| *Igf1R* | 0.83 | 1.28 | 0.67 | 0.97 | 0.81 | 0.76 | -1.27 | 0.02 |
| *Igf2R* | 0.88 | 1.43 | 0.88 | 1.11 | 1.00 | 0.78 | -1.13 | 0.11 |
| *Irf2* | 1.20 | 1.21 | 1.08 | 1.04 | 0.90 | 0.86 | -1.14 | 0.02 |
| *Lamb1* | 1.22 | 1.58 | 1.25 | 1.23 | 1.03 | 0.77 | -1.11 | 0.13 |
| *Lef1* | ND | ND | 2.27 | 2.18 | ND | ND | ND | ND |
| *Mmp2* | 0.84 | 2.21 | 0.81 | 0.97 | 0.96 | 0.44 | -1.42 | 0.26 |
| *Mycbp* | 1.33 | 1.22 | 0.87 | 1.03 | 0.65 | 0.85 | -1.33 | 0.10 |
| *Mycl* | 1.48 | 1.02 | 0.92 | 0.95 | 0.62 | 0.93 | -1.29 | 0.16 |
| *Ncl* | 1.18 | 1.63 | 0.99 | 1.01 | 0.84 | 0.62 | -1.37 | 0.11 |
| *Neat1* | 0.96 | 1.30 | 0.54 | 0.87 | 0.57 | 0.67 | -1.62 | 0.05 |
| *Neat1-2* | 0.37 | 0.78 | 0.41 | 0.59 | 1.09 | 0.75 | -1.08 | 0.17 |
| *Nfil3* | 1.47 | 1.35 | 1.28 | 1.10 | 0.87 | 0.82 | -1.18 | 0.03 |
| *Nkrf (NRF)* | 1.32 | 1.85 | 1.13 | 1.14 | 0.85 | 0.62 | -1.36 | 0.12 |
| *Nono* | 1.27 | 1.49 | 1.06 | 1.21 | 0.84 | 0.81 | -1.22 | 0.01 |
| *Nr1D1 (rev-erb-a)* | 1.04 | 1.18 | 0.87 | 1.03 | 0.83 | 0.87 | -1.17 | 0.02 |
| *p16INK4 (Cdkn2A)* | 1.07 | 0.98 | 0.83 | 1.31 | 0.78 | 1.33 | 1.06 | 0.28 |
| *p27kip1 (Cdkn1B)* | 1.17 | 1.39 | 0.99 | 1.06 | 0.85 | 0.77 | -1.24 | 0.04 |
| *Pdgfa* | 1.06 | 1.32 | 0.85 | 1.15 | 0.80 | 0.87 | -1.20 | 0.04 |
| *Pdgfb* | 1.46 | 1.16 | 0.86 | 1.20 | 0.59 | 1.03 | -1.23 | 0.22 |
| *Per1 (Period-1)* | 1.17 | 1.49 | 0.99 | 1.04 | 0.84 | 0.70 | -1.30 | 0.07 |
| *Pfkm (Pfk1)* | 0.96 | 1.18 | 0.89 | 1.02 | 0.92 | 0.86 | -1.12 | 0.03 |
| *Pgf (Plgf)* | 1.19 | 1.09 | 1.32 | 1.43 | 1.11 | 1.31 | -0.83 | 0.10 |
| *Prox1* | 1.26 | 1.68 | 1.23 | 1.25 | 0.98 | 0.74 | -1.16 | 0.12 |
| *Pspc1* | 1.07 | 1.17 | 0.97 | 1.13 | 0.91 | 0.97 | -1.07 | 0.03 |
| *Rbm14* | 1.12 | 1.44 | 0.91 | 1.11 | 0.82 | 0.77 | -1.26 | 0.02 |
| *Rpl10A* | 0.91 | 0.88 | 0.98 | 1.04 | 1.07 | 1.18 | 1.12 | 0.06 |
| *Rps2* | 0.94 | 1.14 | 0.86 | 0.96 | 0.92 | 0.84 | -1.14 | 0.04 |
| *Rps25* | 0.69 | 1.10 | 0.93 | 0.88 | 1.34 | 0.80 | 1.07 | 0.27 |
| *Rrbp1* | 0.90 | 1.11 | 0.65 | 1.13 | 0.72 | 1.02 | -1.15 | 0.15 |
| *Serpine1 (PAI1)* | 0.74 | 1.51 | 0.85 | 1.07 | 1.15 | 0.71 | -1.08 | 0.22 |
| *Setd7 (Set7)* | 1.19 | 1.36 | 1.02 | 0.99 | 0.86 | 0.73 | -1.26 | 0.06 |
| *Sfpq* | 1.13 | 1.90 | 1.00 | 1.30 | 0.88 | 0.69 | -1.27 | 0.10 |
| *Shmt1* | 1.14 | 1.39 | 0.99 | 1.19 | 0.87 | 0.86 | -1.15 | 0.00 |
| *Slc7A1 (Cat-1)* | 0.87 | 1.49 | 0.74 | 1.02 | 0.85 | 0.68 | -1.30 | 0.08 |
| *Srebf1 (Srebp1)* | 1.00 | 1.59 | 0.64 | 1.11 | 0.64 | 0.70 | -1.49 | 0.03 |
| *Sstr2 (Sst2)* | 0.65 | 0.98 | 0.87 | 1.12 | 1.34 | 1.14 | 1.24 | 0.10 |
| *Thbd (Thrombomodulin)* | 1.05 | 0.74 | 0.51 | 0.51 | 0.48 | 0.69 | -1.71 | 0.10 |
| *Trp53* | 1.14 | 1.25 | 0.95 | 0.98 | 0.83 | 0.79 | -1.24 | 0.02 |
| *Txnip* | 1.17 | 1.29 | 1.02 | 0.99 | 0.87 | 0.77 | -1.22 | 0.05 |
| *Utrn (Utrophin)* | 0.93 | 1.59 | 0.99 | 1.10 | 1.07 | 0.69 | -1.14 | 0.19 |
| *Vash1* | 1.41 | 1.82 | 1.05 | 1.02 | 0.74 | 0.56 | -1.54 | 0.09 |
| *Vegfa* | 1.02 | 1.64 | 0.88 | 1.01 | 0.86 | 0.62 | -1.35 | 0.12 |
| *Vegfb* | 1.40 | 1.58 | 1.02 | 1.06 | 0.73 | 0.67 | -1.43 | 0.03 |
| *Vegfc* | ND | ND | 1.20 | ND | ND | ND | ND | ND |
| *Vegfd (Figf)* | 1.23 | 1.13 | 1.06 | 1.04 | 0.86 | 0.93 | -1.12 | 0.03 |
| *Xiap* | 1.44 | 1.45 | 1.07 | 1.10 | 0.74 | 0.76 | -1.33 | 0.01 |

**Supplementary file 8. Change of mRNA recruitment into polysomes following Neat1 or Neat1_2 knock-down.**

HL-1 cardiomyocytes were transfected with gapmer Neat1, Neat1-2, or control. Polysomes were purified on sucrose gradient as described in Star Methods. RNAs were purified from cytoplasmic extracts and from pooled polysomal fractions and analyzed on a Fluidigm deltagene PCR array from two biologicals replicates (cell culture dishs and cDNAs), each of them measured in three technical replicates (PCR reactions). mRNA levels in polysomes (polysomal RNA/ total RNA) were analyzed for each gene. Relative quantification (RQ) of mRNA level was calculated using the 2–ΔΔCT method with normalization to GAPDH mRNA and to HL-1 tranfected by gapmer control, and is shown as fold change of expression. RQ1 and RQ2 correspond to two independent experiments. To measure the fold change of repression, the mean is expressed as -1/RQ for the values < 1 (yellow column).
